# Triggering and modulation of a complex behavior by a single peptidergic command neuron in *Drosophila*

**DOI:** 10.1101/2024.04.27.591209

**Authors:** Magdalena Fernandez-Acosta, Rebeca Zanini, Fabiana Heredia, Yanel Volonté, Juliane Menezes, Katja Prüger, Julieta Ibarra, Maite Arana, María Sol Perez, Jan A Veenstra, Christian Wegener, Alisson Gontijo, Andrés Garelli

## Abstract

At the end of their growth phase, *Drosophila* larvae remodel their bodies, firmly glue themselves to a substrate, and harden their cuticle in preparation for metamorphosis. This process is termed pupariation and it is triggered by a surge in the steroid hormone ecdysone. Substrate attachment is achieved by a recently-described pupariation subprogram called glue expulsion and spreading behavior (GSB). An epidermis-to-CNS Dilp8-Lgr3 relaxin signaling event that occurs downstream of ecdysone after pupariation initiation is critical for unlocking progression of the pupariation motor program towards GSB, but the factors and circuits acting downstream of Lgr3 signaling remain unknown. Here, we screened for such factors using cell type-specific RNA interference (RNAi) and behavioral monitoring. We identify Myoinhibiting peptide (Mip) and its highly conserved neuronal receptor, Sex peptide receptor (SPR), as a critical neuropeptidergic signaling pathway required to trigger and modulate multiple action components of GSB. In addition, we find that Mip is specifically required in a pair of descending neurons, whose optogenetic activation at a specific competence window triggers GSB-like behavior and whose neurogenetic silencing completely abrogates GSB without overtly affecting other pupariation components. This strongly suggests that these descending Mip neurons are developmentally-regulated GSB command neurons. Dissection of the GSB action components via muscle calcium-level monitoring coupled with cell-type specific RNAi indicates that Mip acts on multiple SPR-positive neuronal populations, which collectively define and pattern the sequence and timing of GSB actions. Hence, we have identified a pair of descending command neurons that utilize both synaptic transmission and neuropeptidergic signaling to trigger and modulate a complex innate behavior in *Drosophila*. Our results advance our molecular and cellular understanding of pupariation control, reveal the complexity of glue expulsion and spreading behavior control, provide insight into conserved aspects of Mip-SPR signaling in animals, and contribute to the understanding of how multi-step innate behaviors are coordinated in time and with other developmental processes through command neurons and neuropeptidergic signaling.

## INTRODUCTION

Neuronal circuits are the structural framework of interconnected neurons that form the basis of information processing and transmission within the nervous system. They are responsible for the integration and transformation of sensory inputs, motor outputs, and higher-order functions. However, the activity of neuronal circuits is not solely determined by their anatomical connectivity; it is profoundly influenced by the action of neuromodulators that alter intrinsic excitability, synaptic plasticity, and overall network dynamics (Marder, 2012; Marder *et al*., 2022; Burke *et al*., 2019). Together, these activities shape the output of neuronal circuits and ultimately drive specific behaviors.

The genetic tractability of *Drosophila melanogaster* and the relative simplicity of its central nervous system (CNS) facilitate the identification of neural circuits that control specific behaviors. *Drosophila* larvae display a wide variety of locomotor behaviors such as backward and forward crawling, burrowing, head sweep, turn, and rolling (Clark *et al*., 2018; Gowda *et al*., 2021). These result from the intra- and inter-segmental coordination of motor neuron activity mediated by local interneurons in the ventral nerve cord (VNC) (Kohsaka *et al*., 2019; Berni *et al*., 2012), while selection of the locomotor behavior is directed by higher order neurons influenced by internal and environmental stimuli (Carreira-Rosario *et al*., 2018; Fushiki *et al*., 2016; Jovanic *et al*., 2016, Takagi *et al.,* 2017). These neurons that mediate action selection are defined as command neurons if it can be experimentally demonstrated that they are both necessary and sufficient to initiate the behavior (Kupfermann and Weiss, 1978)

Some motor action patterns performed at specific developmental stages are controlled by hormones. One example is the ecdysis behavior, a typical innate behavior—those which are genetically-encoded and do not require previous learning (Lorenz, 1981)—which consists of a highly stereotyped sequence of movements that sheds the old cuticle at each molt. Ecdysis is triggered by the release of Ecdysis Triggering Hormone (ETH) and requires the concerted action of ETH, Eclosion Hormone and the Crustacean Cardioactive Peptide for proper execution (Clark *et al*., 2004; Ewer *et al*., 1997).

Pupariation is another example of a complex innate behavior that occurs after the last larval molt of *Drosophila* and other fly species, whereby larvae reshape and harden their cuticle to form a protective capsule within which they undergo metamorphosis (Denlinger, 1994; Fraenkel *et al*., 1973; Zdarek *et al*., 1972). The whole pupariation process is under the control of the steroid hormone Ecdysone, which acts parallelly on the epidermis to induce both cuticle sclerotization (Lam *et al*., 2000; Warren *et al*., 2006) and expression of *Drosophila insulin-like peptide 8* (*dilp8*) (Heredia *et al*., 2021; Blanco-Obregon *et al*., 2022), and on the CNS to initiate the pupariation motor program (PMP) that reshapes the body (Heredia *et al*., 2021). The PMP consists of three consecutive stages named after the most conspicuous one, the Glue-Spreading Behavior (GSB) (Heredia *et al*., 2021). PMP starts with pre-GSB, a series of whole-body contractions that shorten the larval body, evert the anterior spiracles, and irreversibly retract the most anterior larval segments. It is followed by GSB proper, which consists of the expulsion of glue—a proteinaceous solution produced by the larval salivary gland—and its spreading over the ventral surface of the body, which promotes adhesion of the future puparium to the underlying substrate. The motor program culminates with post-GSB, a sustained pulsatile muscle activity pattern that maintains the remodeled body shape achieved during the pre-GSB stage, while the cuticle is hardening (Figure 1A). Coordinated execution of the peripheral cuticle morphogenetic program and the centrally controlled motor program is achieved by the action of epidermally-derived Dilp8. This hormone promotes the progression of the PMP, which otherwise remains locked in the initial pre-GSB stage (Heredia *et al*., 2021).

**Figure 1.**
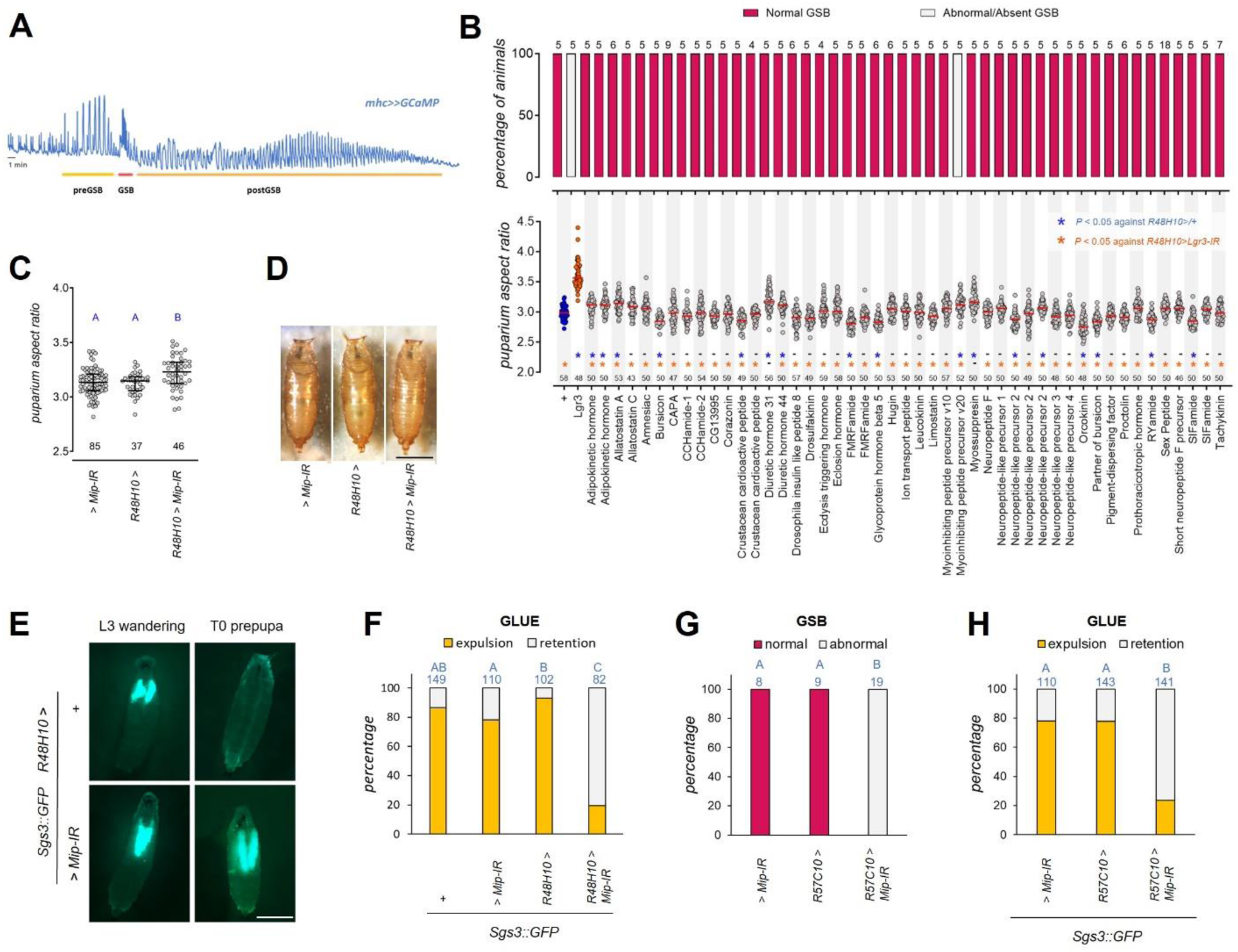
Proper Glue Spreading Behavior requires Mip in neurons. A) Representative profile of muscle calcium fluctuations during the stages of the pupariation motor program detected with muscle-expressed GCaMP (*mhc>>GCaMP*). B) Summary of the RNAi screen to detect neuropeptides required for the PMP by crossing *R48H10>* with RNAi lines targeting the indicated genes. Percentage of animals performing normal GSB (top) and puparium aspect ratio (bottom). Blue asterisk, *P* < 0.05 compared with the negative control (*R48H10>/+*). Orange asterisk, *P* < 0.05 against *R48H10>Lgr3-IR* positive control. Black numbers, N (number of animals). Dots represent one animal. Horizontal orange line, median. Kruskal-Wallis test, followed by Dunn’s test with correction for multiple comparisons. C) RNAi knockdown of *Mip* (*Mip-IR*) in *R48H10>* cells slightly increases puparium aspect ratio. Dot plots showing puparium aspect ratio of indicated genotypes. Dots represent one animal, horizontal bar, median, error bars 25 and 75 percentile. Black numbers, N. Same blue letter, *P* > 0.05, ANOVA and Tukey’s HSD test. D) RNAi knockdown of *Mip (Mip-IR)* in *R48H10>* cells does not induce gross defects in puparium shape. Scale bar, 1 mm. E) RNAi knockdown of *Mip* (*Mip-IR*) in *R48H10>* cells suppresses glue expulsion. All genotypes carry an Sgs3::GFP transgene that labels a glue protein (cyan). F) RNAi knockdown of *Mip* (*Mip-IR*) in *R48H10>* cells suppresses glue expulsion. Shown is the percentage of larvae of indicated genotypes that expel glue (labeled by Sgs3::GFP). Blue numbers, N. Same blue letter, *P* > 0.008, pairwise Fisher’s exact test with Bonferroni correction (alpha = 0.05, 6 tests). G) Pan-neuronal silencing of Mip alters GSB. Shown is the percentage of larvae of indicated genotypes that perform a normal GSB. Blue numbers, N. Same blue letter, *P* > 0.016, pairwise Fisher’s exact test with Bonferroni correction (alpha = 0.05, 3 tests). H) Pan-neuronal silencing of Mip suppresses glue expulsion. Percentage of animals of depicted genotypes releasing glue. Blue numbers, N. Same blue letter, *P* > 0.016, pairwise Fisher’s exact test with Bonferroni correction (alpha = 0.05, 3 tests). Data for F and H, and Figure 2H were collected in parallel as part of the same experiment and represented separately for clarity. For this reason, *> Mip-IR* values are the same in all cited panels.

Dilp8 is a relaxin-like peptidic hormone, initially identified as a systemic tissue homeostasis factor secreted from aberrantly-growing imaginal discs during the larval growth phase (Colombani *et al*., 2012; Garelli *et al*., 2012). Dilp8 acts via the neuronal Leucine-rich repeat-containing G protein-coupled receptor 3 (Lgr3) both during the larval stage (Garelli *et al*., 2015; Vallejo *et al*., 2015; Colombani *et al*., 2015; Jaszczak *et al*., 2016) and during pupariation (Heredia *et al*., 2021). In the latter stage, Lgr3 is required in up to six VNC Lgr3-positive (6VNC Lgr3+) interneurons whose cell bodies are organized in pairs around the posterior boundaries of thoracic segments 1 to 3, from where they project anteriorly, some reaching the brain (Heredia *et al*., 2021). The Dilp8-Lgr3 relaxin-like pathway is essential for the timely and proper execution of the PMP behaviors. However, it remains unclear how this single hormonal signal is translated into a succession of complex and diverse motor action routines. Similarly, the mechanism whereby the 6VNC Lgr3+ interneurons convey their message to the circuit that shapes the pupariation motor outputs has not yet been defined.

Here, we identify myoinhibitory peptide (Mip; also known as allatostatin B (AstB)) as a neuromodulating signal acting downstream of Dilp8 during the PMP. We find that Mip is required in a pair of descending neurons that traverse the whole VNC to shape the PMP behavioral routine associated with glue expulsion. When Mip is absent, the glue spreading behavior still initiates, but with an aberrant sequence of movements, whereas when Mip descending neurons are neurogenetically silenced or optogenetically activated, the behavior is completely abrogated or triggered, respectively. These results strongly suggest that these neurons are–apart from Mip neuromodulatory neurons–also GSB command neurons. We further show that Mip acts via its canonical receptor, the sex peptide receptor (SPR), during pupariation, and that it has a widespread neuromodulatory effect on SPR-positive glutamatergic, and cholinergic neurons, highlighting the importance of modulatory signals to reconfigure the activity of motor circuits and produce unique stereotyped outcomes.

## RESULTS

### An RNAi screen for neuropeptides acting downstream of the Dip8-Lgr3 pathway during pupariation

During pupariation, Dilp8 relaxin hormone produced by the cuticle epidermis acts on 6VNC Lgr3+ interneurons to unlock the progression of the PMP. RNA interference (RNAi)-mediated knockdown of *Lgr3* in this restricted cell population by means of intersectional genetics combining the *R18A01* and *R48H10 cis-*regulatory modules (CRMs; Pfeiffer *et al*., 2008; Jenett *et al*., 2012; Li *et al*., 2014) blocks the progression of the PMP and generates aberrant puparia with an increased length/width ratio (aspect ratio; Heredia *et al*., 2021). The neuroanatomy of the 6VNC Lgr3+ interneurons suggests that direct control of motor neuron activity is unlikely to explain the full spectrum of Dilp8-Lgr3 activity during pupariation (Heredia *et al*., 2021). Instead, it suggests that other neurons and/or neuromodulators should be acting downstream of the 6VNC Lgr3+ interneurons to organize the activity of motor pattern generators. To identify those putative mediators, we performed a genetic screen for neuropeptides acting downstream of Dilp8 in the PMP.

We reasoned that if a neuropeptide is required downstream of Dilp8/Lgr3 signaling in the 6VNC Lgr3+ interneurons, its encoding gene could be readily identified by silencing it in these interneurons and observing the effect this has on puparium shape and/or pupariation behavior. We screened for such neuropeptide-encoding genes using a cell-type restricted RNAi approach based on the GAL4-UAS system. A set of 47 UAS-driven RNAi lines from the TRiP collection targeting 40 neuropeptide-encoding genes (Supplementary Table 1) was crossed with the *R48H10-GAL4* (*R48H10>*) line, which drives expression in ∼180 cells in the CNS, including the 6VNC Lgr3+ interneurons, and, importantly, has no epistatic effect on GSB, in contrast to lines using the *R18A01* CRM, which would be an alternative line to target the same interneurons (Heredia *et al*., 2021). We evaluated the progeny of these crosses for the aspect ratio of their puparia and their performance of GSB by visual inspection of animals registered under white light in a custom-made pupariation device, as previously described (Heredia *et al*., 2021). Puparium aspect ratio was calculated in 40 to 60 individuals and the performance of GSB was classified as normal or abnormal in at least five larvae per genotype. The behavior was considered abnormal when it was either not present or present with an abnormal execution of the sequence of movements (Supplementary video 1). *R48H10>* crossed with *w^1118^* (*R48H10>*+) or *Lgr3* RNAi (*Lgr3-IR*) (*R48H10>Lgr3-IR*) was used as wild-type (negative control) or as positive control, respectively. Consistent with previous results (Heredia *et al.,* 2021), *R48H10>Lgr3-IR* animals had a statistically significantly-increased aspect ratio compared to controls (16.4% median increase, *P* < 0.0001, against *R48H10>+* control, Dunn’s test), and a complete suppression of GSB in all of the animals tested (*N* = 5) (Figure 1B).

### Identification of *Myoinhibiting peptide precursor* (*Mip*) as a critical factor for proper GSB

Of the 47 different RNAi lines screened in *R48H10>* neurons, only one, directed against *Myoinhibiting peptide precursor* (*Mip*; RNAi line *UAS-Mip-IR-V20* (*HMS02244*) (Ni *et al*., 2011), hereafter referred to as *UAS-Mip-IR*) affected the execution of GSB (0/5 animals performed normal GSB, as compared to 5/5 controls, as well as all the other tested RNAi lines) (Figure 1B, upper panel). *R48H10>Mip-IR* also induced a small increase in the aspect ratio of puparia, which was statistically-significant (1.8% median increase relative to *R48H10>+* controls, *P = 0.0016*, Dunn’s test) (Figure 1B, lower panel) and independently-reproducible (*P* = 0.002, against *R48H10>+* and *+>Mip-IR* controls, ANOVA and Tukey’s HSD test) (Figure 1C), but was smaller than the positive control (*R48H10>Lgr3-IR*) (*P* = 0.0004, compared to *R48H10>Mip-IR*, Dunn’s test; Figure 1B). Furthermore, the overall shape of *R48H10>Mip-IR* pupae did not resemble that of *dilp8* or *Lgr3* mutants (Figure 1D), which have severe defects in the retraction of the most anterior body segments and internalization of the mouth hooks (Heredia *et al*., 2021). A second RNAi line against *Mip* (*UAS-Mip-IR-V10* (*JF02145*) (Ni *et al*., 2009)) had no effect on the screen (Figure 1B), but produced the same result as the *UAS-Mip-IR-V20* RNAi line when driven together with a *UAS-Dicer2 (Dcr2)* cassette, which is routinely used to boost hairpin processing and gene silencing, particularly in neurons (Dietzl *et al*., 2007) (0 out of 5 animals performed normal GSB; *P = 0.0079, Fisher’s exact test*, 2.8% median increase in aspect ratio relative to *UAS-Mip-IR/+* controls, *P = 0.038*, Dunn’s test; Supplementary Figure 1). The Dcr2-dependent effect of the V10 RNAi construct is consistent with previous reports (Perkins *et al*., 2015). These results indicate that Mip is required in R48H10-expressing cells for proper GSB and puparium aspect ratio, albeit the effect on aspect ratio is weaker and categorically different from the effect of the Dilp8-Lgr3 pathway.

### Mip is required in a restricted population of neurons for glue expulsion

GSB starts with a strong and sustained contraction of the anterior segments that forces the release of the proteinaceous content of the salivary glands, which is then spread over the ventral surface of the animal. The result of our screen indicates that knockdown of *Mip* affects the behavior associated with glue vspreading, but this does not necessarily imply that the preceding release of glue is similarly impacted. To directly assess the role of Mip in glue synthesis and expulsion we used *Sgs3::GFP*, a transgene that encodes a glue protein tagged with GFP under the control of its own promoter (Biyasheva *et al.,* 2001). We monitored Sgs3::GFP fluorescence prior and after pupariation in wandering L3 larvae and in newly formed white prepupae, respectively, of controls and *Mip*-silenced animals (*R48H10>Mip-IR*). Prior to pupariation, we found similar levels of Sgs3::GFP in the salivary glands of control and *R48H10>Mip-IR* 3rd instar larvae, indicating that *Mip* is not required for glue synthesis. In contrast, after pupariation, we found that while 86.6% of control animals showed evidence of glue expulsion during pupariation, only 19.5% of *R48H10>Mip-IR* animals did so (*N* = 82 larvae; *P* < 0.00001 against controls, Fisher’s exact test) (Figure 1E-F). These results indicate that Mip is required in R48H10-positive cells for glue expulsion. This, together with the results described above, showed that *Mip* is critical for the proper execution of both behaviors that constitute GSB: glue expulsion and glue spreading.

The *Mip* gene of *Drosophila melanogaster* encodes a prepropeptide predicted to generate five mature myoinhibiting peptides of the B-type family of allatostatins with the shared W(X_6_)Wamide structure (Williamson *et al*., 2001), four of which have been confirmed by mass spectrometry (Predel *et al*., 2004; Baggerman *et al*., 2005; Reiher *et al*., 2011). B-allatostatins/myoinhibiting peptides were named after their ability to inhibit the synthesis of juvenile hormone in the corpora allata of the cricket *Gryllus bimaculatus* (Lorenz *et al*., 1995) and to suppress the spontaneous muscle contractions in the hindgut and oviduct of *Locusta migratoria* (Schoofs *et al*., 1991). *Mip* is expressed in the gut and CNS of *Drosophila* where it participates in the control of sleep (Oh *et al*., 2014), satiety (Min *et al*., 2016), mating (Jang *et al*., 2017; Shao *et al*., 2019), and olfaction (Hussain *et al*., 2016; Sizemore *et al*., 2023). No clear function during larval stages has been demonstrated, besides a proposed role in the control of ecdysis based on similarities in neuronal expression pattern and neuropeptide co-expression profile with *Manduca sexta* (Kim *et al*., 2006). The finding that *Mip* is required for the execution of GSB is the first genetically verified role for this neuropeptide in larvae.

As in adult flies, Mip is found both in the CNS and gut at the larval stage (Williamson *et al*., 2001; Veenstra, 2009; Chen *et al*., 2016). To test if neurons are the source of the Mip activity required for GSB, we used a pan-neuronal GAL4 line, *R57C10-GAL4* (*R57C10>*) (Pfeiffer *et al*., 2008), and *UAS-Mip-IR* to silence *Mip* expression in neurons (*R57C10>Mip-IR*). We found that neuronal-specific knockdown of *Mip* had the same effect as *R48H10>Mip-IR*, impairing both the execution of proper GSB (0 out of 19 performed normal GSB, as compared to 8 out of 8 and 9 out of 9 *UAS-Mip-IR/+* and *R57C10>+* controls, respectively, *P* < 0.00001, Fisher’s exact test) (Figure 1G) and glue expulsion (23.4% expelled glue, as compared to 78.2% and 77.6% for *UAS-Mip-IR/+* and *R57C10>+* controls, *P* < 0.00001, Fisher’s exact test, N = 141, 110, 143 respectively) (Figure 1H). These results demonstrated that Mip-positive (Mip+) neurons are critical for proper GSB and suggested that other possible sources of the neuropeptide are unlikely to contribute significantly to this behavior.

### Mip is neither expressed nor required in Lgr3+ VNC neurons for proper GSB

*Mip* expression in the *Drosophila* larval brain has been studied by *in situ* hybridization and immunostaining with slightly different results based on the technique and the reagents used (Williamson *et al.,* 2001; Park *et al*., 2008; Kim *et al*., 2006; Santos *et al*., 2007). Expression of *Mip* has been mapped to 4-5 pairs of neurons in the protocerebrum (Williamson *et al*., 2001, Park *et al*., 2008), a pair of bilateral neurons per hemisegment from thoracic segment T1 to abdominal A7, and four pairs in the tip of the VNC, in abdominal segments A8/9 (Kim *et al*., 2006; Santos *et al*., 2007). Mip*+* neurons emit longitudinal projections that travel across the VNC along the lateral and midline tracts, as well as transverse projections in each hemisegment that meet in the midline (Santos *et al*., 2007). This reported distribution of Mip+ cells within the VNC appeared to be different from that of the 6VNC Lgr3+ interneurons, raising the possibility that these cell populations do not overlap, which contrasted in principle with our initial objective of finding neuropeptides produced in the 6VNC Lgr3+ interneurons as a direct response to Dilp8.

To independently and directly confirm this observation, we labeled Mip+ cells using an anti-Mip antibody developed against the American cockroach Mip (*Periplaneta americana,* Predel *et al*., 2001) and assessed their distribution in the CNS of larvae expressing membrane-targeted red fluorescent protein, mCD8::RFP (*UAS-mCD8::RFP*) (Pfeiffer *et al*., 2010), in *R48H10>*-positive cells (*R48H10>mCD8::RFP)* (Figures 2A-D). In these animals, the subset of R48H10>mCD8::RFP-positive cells corresponding to the 6VNC Lgr3+ interneurons can be easily identified by their anatomical position in thoracic segments T1 to T3. Immunohistochemical analyses showed the same Mip expression pattern as already described by Santos *et al*. (2007), as expected, and some additional cells in the suboesophageal zone that had not been previously reported because the authors focused their analysis on the VNC (Figure 2A). Critically, we found no Mip+ cell that was also positive for *R48H10>mCD8::RFP* in segments T1-T3 (Figure 2B and D). Instead, a large bilateral neuron in the protocerebrum that expressed very high levels of Mip was also positive for *R48H10>mCD8::RFP* (Figure 2B-C). Apart from these brain neurons, a few smaller cells in the lateral posterior abdominal segments of the VNC also co-expressed both markers, but the pattern varied between preparations and it was not always possible to find the contralateral cell (Figure 2B, arrowheads). As the protocerebral neuronal pair appears to be the only one in the *R48H10>* expression field with consistent levels of Mip immunoreactivity, we hypothesized it is the only one strongly susceptible to *Mip* RNAi silencing and thus that it could coincidently be the neuron where Mip is required for proper GSB. If this were true, silencing of Mip in the VNC should have no effect on GSB. To test this, we targeted *Mip* RNAi to the VNC using *teashirt* (*tsh*)*-GAL4* (Roder *et al*., 1992; Calleja *et al*., 1996) (*tsh>Mip-IR*), which drives expression in all thoracic and abdominal segments except for A8/9. We found that, while *tsh>Mip-IR* completely eliminated detectable Mip immunoreactivity in the paired bilateral neurons of all thoracic and abdominal segments except for A8/9, demonstrating efficient RNAi in those cells (Figure 2E), it did not affect aspect ratio (Figure 2F), GSB execution (Figure 2G), or glue expulsion (Figure 2H). As hypothesized, 100% of *tsh>+* and *tsh>Mip-IR* larvae did GSB (N = 5 and 20 respectively, *P =* 1.000, Fisher’s exact test), and expelled glue as well or better than the controls (96.2% of *tsh>Mip-IR* larvae expelled glue, N = 79, as compared to 78.2% of *UAS-Mip-IR/+,* N = 110*, P* = 0.0006 and 93.7% of *tsh>+*, N = 126, *P* = 0.5358, Fisher’s exact tests) (Figure 2G-H). These results strongly suggested that the Mip activity required for proper GSB is not derived from the 6VNC Lgr3+ interneurons or any other neuron with cell bodies originating from VNC segments T1-A7. As the Mip+ and R48H10-positive protocerebral neuronal pair was the only candidate neuron left, we conclude it has a central role in the modulation of GSB via Mip, and also that Mip and Lgr3 are acting in two different neuronal populations.

**Figure 2.**
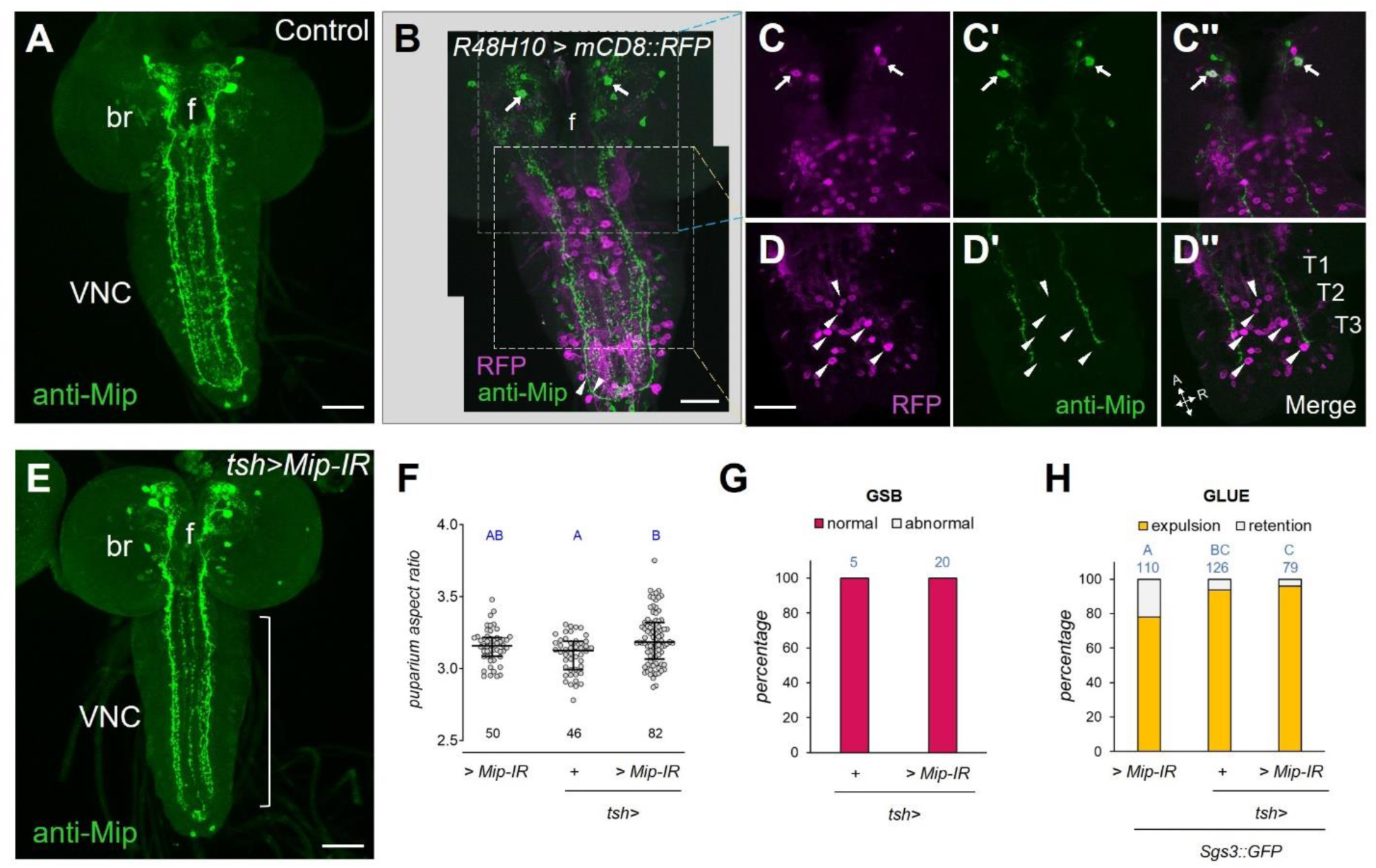
Mip is neither expressed nor required in 6VNC Lgr3+ cells. A) Mip expression pattern (green) in L3 larval CNS. f, esophageal foramen; br, brain; VNC, ventral nerve cord. B) Montage of full projections of two confocal stacks of a single CNS dissected from R48H10>CD8::RFP (magenta) larvae stained with an anti-Mip antibody (green). Arrows and arrowheads point to double-labeled cell bodies. Arrows, protocerebral pair. Arrowheads, 2 lateral posterior abdominal VNC cells. Scale bar, 50 µm. C) Details of the cell bodies of the protocerebral neuronal pair (arrows) depicted in B. Partial projections of a subset of the confocal sections in the regions highlighted as insets (dashed lines) in B. C (R48H10>CD8::RFP, magenta), C’ (anti-Mip antibody, green), and C’’ (merge). D) Details of 6VNC neuron cell bodies (arrowheads) from the confocal stack depicted in B. Partial projections of a subset of the confocal sections in the regions highlighted as insets (dashed lines) in B. D (R48H10>CD8::RFP, magenta), D’ (anti-Mip antibody, green), and D’’ (merge). A, anterior. R, right. T1-3, thoracic segments 1-3. f, esophageal foramen. E) Mip knockdown in the *tsh* expression domain (white bracket, *tsh>Mip-IR)* reduces Mip immunoreactivity (green) in VNC neurons, except for those in its posterior tip. f, esophageal foramen; br, brain; VNC, ventral nerve cord. F) Dot plot showing puparium aspect ratio of indicated genotypes. Dots: one animal. Horizontal bars, median, and 25-75 percentile. Blue numbers, N. Same blue letter, *P* > 0.05, ANOVA and Tukey’s HSD test. G) Mip is not required in the VNC neurons labeled by *tsh>* for proper GSB. Shown is the percentage of larvae of indicated genotypes that perform a normal GSB. Blue numbers, N. *P > 0.05*, Fisher’s exact test. H) Mip is not required in the VNC neurons labeled by *tsh>* for proper glue expulsion. Shown is the percentage of larvae of indicated genotypes that expel glue (labeled by Sgs3::GFP). Blue numbers, N. Same blue letter, *P* > 0.016, pairwise Fisher’s exact test with Bonferroni correction (alpha = 0.05, 3 tests). Data for H and Figure 1F and 1H were collected in parallel as part of the same experiment and represented separately for clarity. For this reason, *> Mip-IR* values are the same in all cited panels.

### Mip is required in a protocerebral pair of neurons for proper GSB

To further confirm that the *Mip-*expressing neuron labeled by *R48H10>* is the neuron where *Mip* is required, we carried out *Mip* RNAi mediated knockdown in restricted neuronal populations using different GAL4 drivers. Namely, we used three Mip-GAL4 drivers: *MipTH-GAL4* (*MipTH>*), and two other transgenic lines with more restricted patterns, *Mip3M-GAL4* (*Mip3M>*) and *Mip2M-GAL4* (*Mip2M>*), developed in this study (see Methods section). The latter two are random insertions of the same construct and label a reduced number of cell bodies in the protocerebrum and abdominal 8/9 segments, as well as the long projections that cross the VNC, but do not label any of the Mip+ bilateral neurons in the thoracic and abdominal segments (Figure 3A-D).

**Figure 3.**
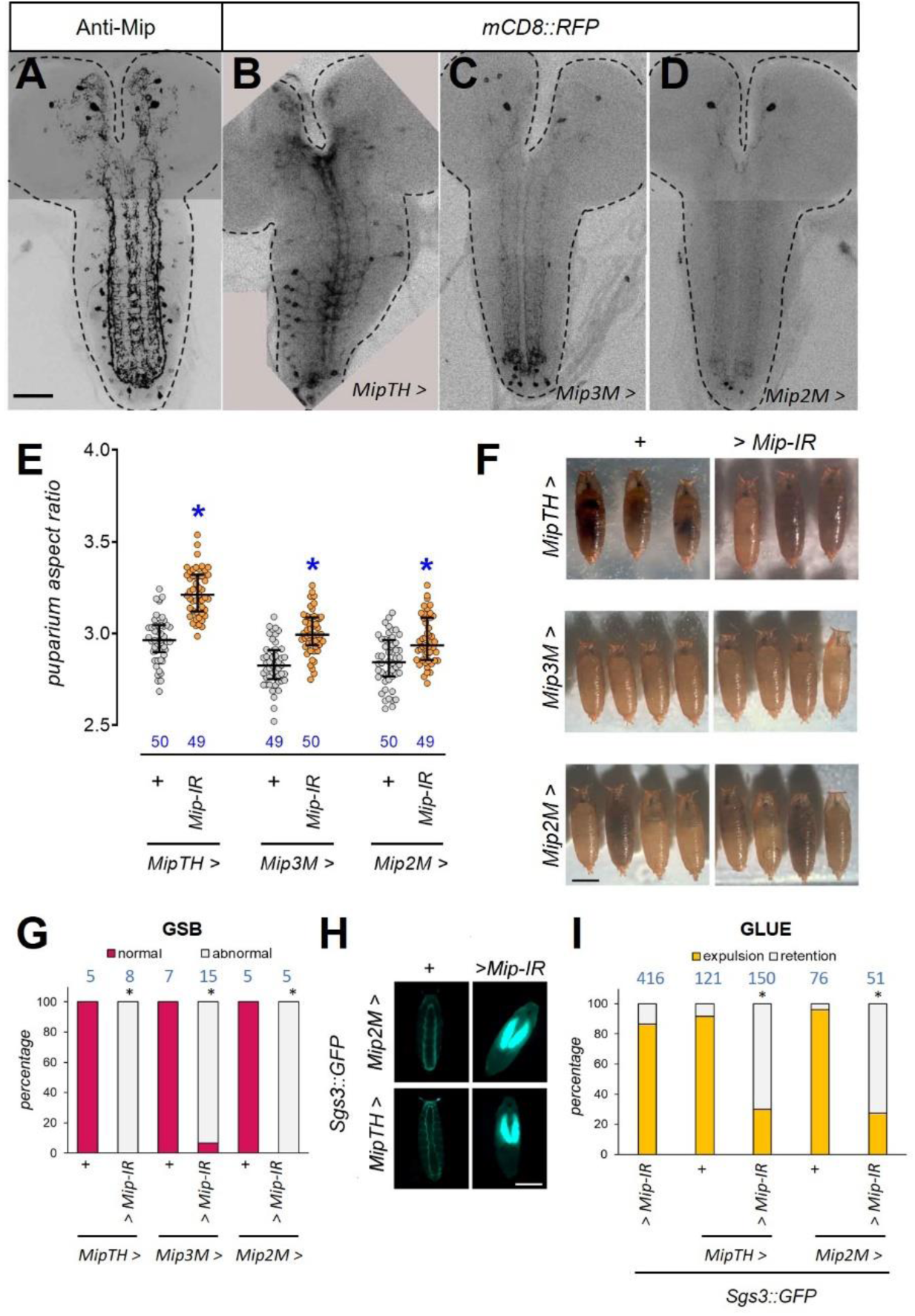
Mip is required in a pair of bilateral brain neurons labeled by *Mip2M>*. A) Montage of confocal slices of 3rd instar larva CNS stained with anti-Mip (black). Dotted lines, CNS contour. Scale bar, 50 µm. B-D) Montages of confocal slices of 3rd instar larva CNS expressing *UAS-CD8::RFP* (black) under the control of different *Mip>* lines (*MipTH>,* B; *Mip3M>,* C; *Mip2M>,* D). Dotted lines, CNS contour. E) RNAi-mediated downregulation of *Mip (Mip-IR*) with different *Mip-GAL4* lines slightly increases the aspect ratio of puparia. Depicted is a dot plot showing puparium aspect ratio of indicated genotypes. Dots: one animal. Horizontal bars, median, and 25-75 percentile. Blue numbers, N. Asterisk, *P* < 0.0001, unpaired t-test against respective GAL4-only controls. F) RNAi-mediated downregulation of *Mip (Mip-IR)* with different *Mip-GAL4* lines does not induce gross defects in puparium shape. Scale bar, 1 mm. G) RNAi-mediated downregulation of *Mip* with different *Mip-GAL4* lines alters the execution of GSB. Shown is the percentage of larvae of indicated genotypes that perform a normal GSB. Blue numbers, N. Asterisk, *P = 0.0008, P < 0.00005* and *P = 0.0008,* Fisher’s exact test. H) Mip is required in *Mip-GAL4-*expressing cells for proper glue expulsion. Displayed are representative larvae of the depicted genotypes carrying an Sgs3::GFP transgene labeling a glue protein (cyan). Scale bar, 1 mm. I) Mip is required in *Mip-GAL4-*expressing cells for proper glue expulsion. Shown is the percentage of larvae of indicated genotypes that expel glue (labeled by Sgs3::GFP). Blue numbers, N. Asterisk, *P* < 0.00001 against *>Mip-IR* and corresponding *GAL4>+* control, pairwise Fisher’s exact test with Bonferroni correction (alpha = 0.05, 3 tests for each driver).

Silencing of *Mip* using either of these three *Mip-GAL4* drivers slightly increased the AR of puparia (Figure 3E), but, similarly to what occurred in *R48H10>Mip-IR* animals, this was not linked to the gross defects in puparium shape characteristic of *dilp8* or *Lgr3* mutation (Figure 3F). We also found that all three manipulations equally negatively affected the execution of GSB (0/8 *MipTH>Mip-IR*, 1/15 *Mip3M>Mip-IR*, and 0/5 *Mip2M>Mip-IR* larvae performed normal GSB, as compared to 5/5, 7/7 and 5/5 of their respective controls, *P* = 0.00078, *P* = 0.00005, and *P* = 0.00794, respectively, Fisher’s exact test) (Figure 3G) and glue expulsion (30.0% and 27.5% of *MipTH>Mip-IR* and *Mip2M>Mip-IR* larvae expelled glue, *P* < 0.00001 and *P* < 0.00001, Fisher’s exact test with Bonferroni correction, as compared to >86% for all control conditions) (Figure 3H-I), consistent with our previous results that established a link between both processes. The fact that the outcome of *Mip* silencing was similar for the three GAL4 drivers tested, indicated that *Mip2M>* (the driver with the sparsest expression pattern) contains the critical cells where *Mip* is required for GSB. This prompted us to characterize the morphology of *Mip2M>* neurons in detail.

### Mip is required in a pair of descending Mip2M interneurons for proper GSB

Co-labeling of *Mip2M>* cells with a *UAS-mCD8::RFP* reporter (*Mip2M>RFP*) and with anti-Mip antibody, showed six Mip2M>RFP-positive cells, all of which also stained positively for anti-Mip (Figure 4A). These included the two large neurons in the protocerebrum that extend long descending projections through the VNC, and other four neurons in segments A8/9. The number of Mip2M neurons in the tip was variable, but all of them were always labeled with the anti-Mip antibody. Despite all of these cells having the potential to provide Mip for GSB execution, the latter four are not *R48H10*>-positive, minimizing their relevance for the behavior and strengthening the case for the protocerebral bilateral pair of neurons as the critical neurons requiring Mip for proper GSB behavior.

**Figure 4.**
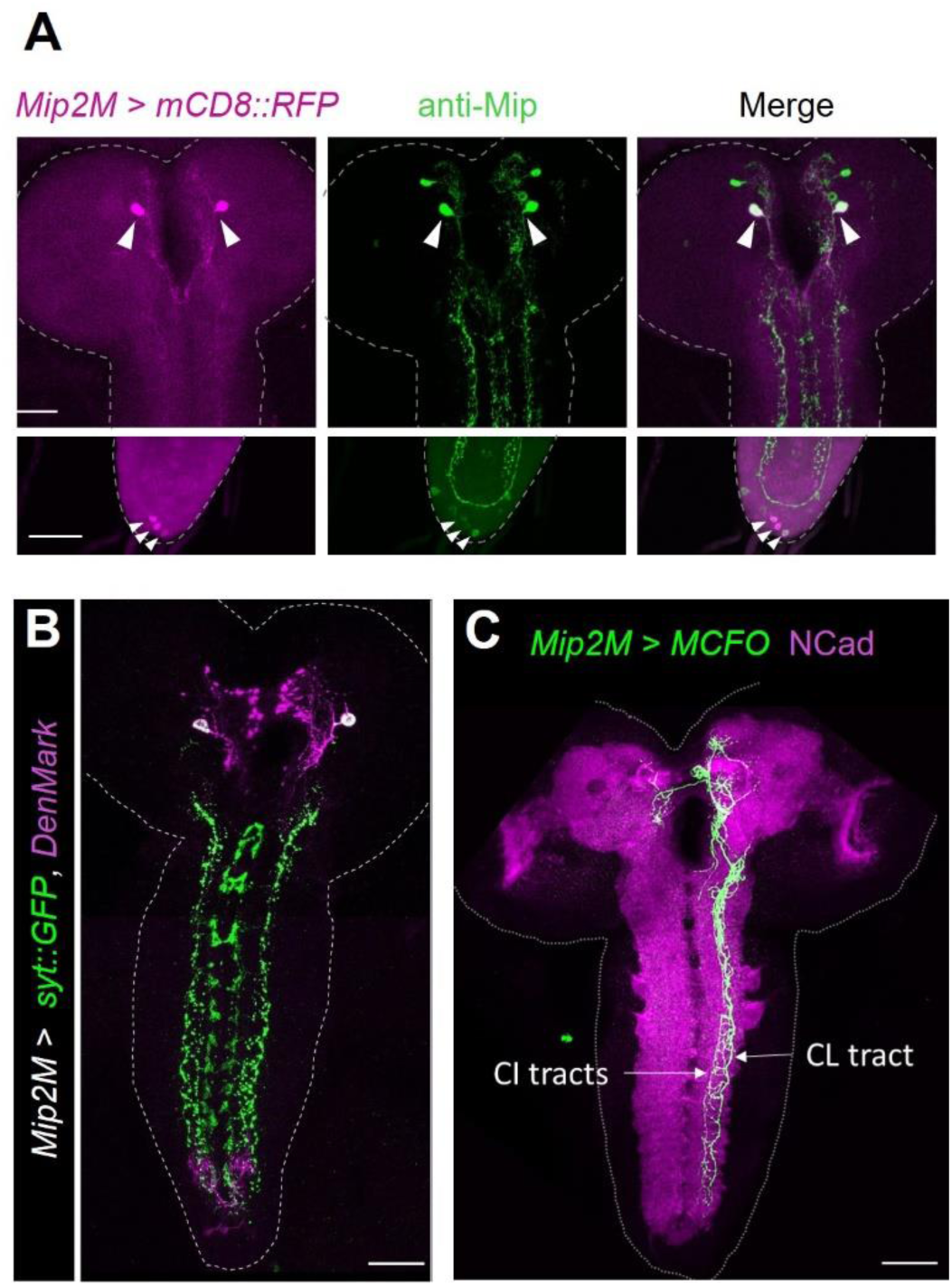
Morphology of the Mip descending neurons. A) All *Mip2M>-*positive cells express Mip (arrowheads). Depicted are confocal image of CNS preparation of 3rd instar larvae stained for *Mip2M>mCD8::RFP* (magenta), and anti-Mip (green). The top and bottom images are from different confocal stacks. Dotted lines, CNS contour. Scale bar, 50 µm. B) Dendritic and axonal compartments of Mip2M neurons labeled with Denmark (magenta) and syt::GFP (green) respectively. Dotted line, CNS contour. Scale bar, 50 µm. C) Single cell clone of a Mip descending neuron showing the contralateral ramification that crosses the midline in the protocerebrum and a long and ipsilateral axonal projection that traverses the ventral nerve cord between the central intermediate (CI) and lateral (CL) longitudinal tracts. Green, *Mip2M>MCFO* (anti-V5) and anti-NCad, magenta, neuropil counterstain. Dotted line, CNS contour. Scale bar, 50 µm.

To determine the neuroanatomy of the two large protocerebral Mip+ neurons and to identify their dendritic and axonal compartments, we expressed the presynaptic marker *UAS-syt::GFP* and the dendrite and cell body marker *UAS-DenMark* under control of *Mip2M>* in 3^rd^-instar-larva-CNS preparations (Figure 4B). We found that these neurons have extensive DenMark-positive dendritic arborizations in the medial compartments of the protocerebrum and in the subesophageal zone, and long axonal syt::GFP-positive projections that travel along the VNC, suggesting that these are descending interneurons. The cells that reside in the posterior end of the VNC extend their dendrites anteriorly, to abdominal segments A7/8 and also project long axons anteriorly that reach the brain lobes, medially to the axons of the protocerebral neurons. This suggested that at least one of these posterior cells are ascending interneurons. Labeling of individual Mip2M> cells using the multicolor flip-out (MCFO) methodology (Nern *et al*., 2015) confirmed these observations and allowed us to refine the anatomy of the Mip2M>-expressing descending and ascending interneurons, as well as to identify a third morphologically distinct neuronal type within the *Mip-2M* population (Supplementary Figure 2). The protocerebral descending interneuron has a large cell body that receives both ipsilateral and contralateral afferences through a small dendritic ramification that crosses the midline anteriorly to the esophagus foramen. The longitudinal axon, on the contrary, remains ipsilateral, with multiple ramified branches oriented towards the midline all along its way to the end of the VNC (Figure 4C and Supplementary Figure 2).

The morphology and location of Mip descending neuron within the VNC is consistent with the description made by Santos *et al*. (2007) of the Mip-positive longitudinal projections that extend adjacent to the central intermediate and central lateral Fasciclin2 tracts (Supplementary Figure 2). These axonal projections run in an intermediate region between the dorsal neuropil that contains motor neuron dendritic arborizations and the ventral neuropil containing sensory axonal projections. This places Mip axonal extensions in a region populated by interneurons and suggests that Mip shapes motor output via modulation of interneurons activity rather than direct contact with motor neurons.

We reasoned that if *Mip* is required exclusively in these neurons for GSB, then expression of *Mip* only in *Mip2M* cells in a *Mip* mutant context should rescue glue expulsion. To test this, we first phenotyped animals homozygous for *Mip^1^*, a genetic null allele in which the whole *Mip* coding sequence has been replaced by a *miniwhite* cassette using homologous recombination (Min *et al.,* 2016). Lack of *Mip* produced a slight, yet statistically significant, increase in the puparium aspect ratio (6.5% increase, *P* < 0.001 ANOVA and Tukey HSD test against homozygous and heterozygous background controls), which, similarly to *Mip* silencing, did not resemble the puparium shape defects emerging from a deficit in the Dilp8/Lgr3 signaling pathway (Figure 5A). Furthermore, *Mip* mutation also perturbed GSB (0 out of 25 of *Mip^1^* homozygous animals did a normal GSB, *P* < 0.00001, Fisher’s exact test with Bonferroni correction, as compared to 20 out of 20 and 5 out of 5 of *Mip^+/+^* and *Mip^1/+^* controls) (Figure 5B) and caused glue retention in the salivary glands in all animals tested (0 out of 20 expelled glue, *P* < 0.00001, Fisher’s exact test, compared to 20 out of 20 of *Mip^1/+^* controls) (Figure 5C-D). To perform the genetic rescue experiment, we then generated a *UAS-Mip* transgene carrying the *Mip* coding sequence devoid of its endogenous 5’ and 3’ untranslated regions (UTRs) and induced its expression in *Mip2M* neurons of *Mip^1^* animals. As a consequence of this manipulation, glue was observed over the surface of the puparium in one quarter of the animals evaluated, indicating that Mip activity in the Mip+ descending neurons can rescue the glue retention phenotype, albeit with low penetrance, and to a level which did not reach statistical significance against one out of three controls assayed (28 out of 106, *P* = 0.0011, *P* = 0.0639, and *P =* 0.0001, Fisher’s exact tests, as compared to 0 out of 28, 0 out of 11, and 0 out of 38 for the three control *Mip^1^* conditions, ++, +>*Mip*, and *Mip2M>+*, respectively) likely due to the relatively small sample size (N = 11) of the second control condition (Figure 5E).

**Figure 5.**
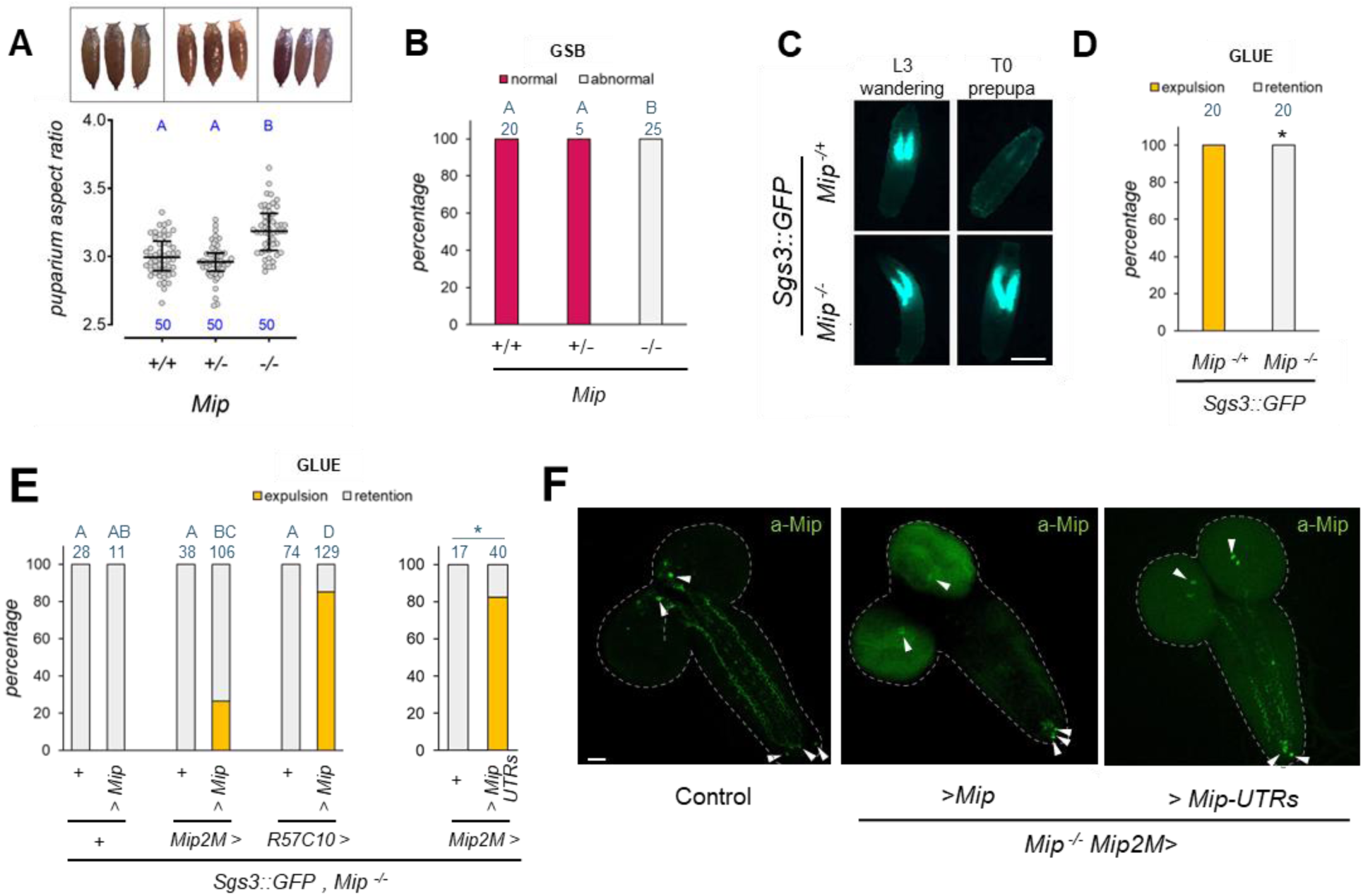
The Mip mutant glue retention phenotype is rescued by neuronal expression of *Mip*. A) *Mip^1^* null mutation increases puparium aspect ratio. Dot plot showing puparium aspect ratio of indicated genotypes (+/+, controls. +/-, *Mip^1^* heterozygotes. *-/-, Mip^1^* homozygotes). Representative images of the respective genotypes are shown in the top. Dots: one animal. Horizontal bars, median, and 25-75 percentile. Blue numbers, N. Same blue letter, *P* > 0.05, ANOVA and Tukey’s HSD test. B) *Mip^1^* mutant larvae perform an abnormal GSB. Shown is the percentage of larvae of indicated genotypes (same as in A) that perform a normal GSB. Blue numbers, N. Same blue letter, *P* > 0.016, pair-wise Fisher’s exact test with Bonferroni correction (alpha = 0.05, 3 tests). C) Abnormal GSB execution causes glue retention in the salivary glands of *Mip^1^* mutant larvae. Shown are representative photos of L3 wandering larvae and white prepupa (T0) carrying a Sgs3::GFP glue reporter (cyan). D) *Mip^1^* mutants do not expel glue during pupariation. Shown is the percentage of larvae of indicated genotypes that expel glue (labeled by Sgs3::GFP). Blue numbers, N. Asterisk, *P <* 0.00001, Fisher’s exact test. E) Left panel: Panneuronal expression of *Mip* rescues glue release to levels similar to control conditions (compare with Figure 1F). Expression of *Mip* in *Mip2M* neurons also rescues glue expulsion, albeit to a lesser extent. Blue numbers, N. Same letter, *P >* 0.008, pair-wise Fisher’s exact test with Bonferroni correction (alpha = 0.05, 6 tests). Right panel: *Mip2M* neuron specific expression of an alternative *Mip* construct carrying endogenous *Mip UTR* sequences rescues glue expulsion in a higher proportion of animals than controls. Blue numbers, N. Asterisk, *P* < 0.0001, Fisher’s exact test. F) Confocal stack projections of L3 larval brains showing Mip immunoreactivity (green) in control (left) and *Mip^1^, Mip2M>Mip* (no UTRs) (center) and *Mip^1^, Mip2M>Mip-UTRs (*with *Mip* with endogenous *Mip* UTR sequences) samples. Mip detection in rescued animals is mostly limited to the cell bodies with the cDNA devoid of endogenous *Mip* UTR sequences and is fully rescued when the complete mRNA sequence is used. Dotted line, CNS contour. Scale bar, 50 µm.

The fact that glue was still retained in the salivary glands of three quarters of the *Mip^1^*, *Mip2M>Mip* rescued pupae could be the consequence of a weak expression of the transgene or a defective transport along the axonal projections of transgenically-derived *Mip* mRNA that lacks endogenous UTR sequences that could have evolved to promote axonal transport (Sahoo *et al*., 2018; Vargas *et al.,* 2022). To verify these alternatives, we evaluated the expression of the transgene by immunocytochemistry in rescued larvae (of the genotype *Mip^1^*, *Mip2M>Mip*). Antibody staining showed that, in contrast to the endogenous Mip protein that is distributed throughout the descending Mip+ neurons, including cell body, dendrites, and the descending longitudinal axon and its varicosities, Mip2M-induced transgenic Mip protein was detectable only at a low level in the cell bodies of the descending neurons (Figure 5F) with occasional faint staining along the ventral nerve cord that resembled Mip distribution along the descending axons (Supplementary Figure 3). These results are consistent with the partial rescue observed, and hinted that Mip is required in the Mip2M descending axon for proper GSB.

To further support this claim, we attempted two other rescue strategies. First, we reasoned that pan-neuronal expression of *Mip* should be sufficient to rescue glue expulsion in *Mip* mutants by providing Mip from the Mip2M descending neurons cell-autonomously and from other nearby VNC neurons, non-cell-autonomously, which is possible as *Mip* encodes secreted peptides. Indeed, when expressed pan-neuronally, *Mip* expression rescued glue retention in 110 out of 129 (85.3%) of *Mip^1^* individuals (*P* < 0.00001, Fisher’s exact test with Bonferroni correction, as compared to 0% for the respective control conditions) (Figure 5E), similar to control conditions (see Figures 1F and H). Second, based on these findings, we hypothesized that the reason why *Mip2M>Mip* gave only partial rescue was because the *UAS-Mip* transgene was giving weak Mip expression. To test this, we generated a new *Mip* transgene carrying both its endogenous 5’ and 3’ UTR sequences (*UAS-Mip-UTRs*), which should facilitate Mip expression. Indeed, immunofluorescence assays using the anti-Mip antibody showed that Mip expression levels in Mip2M-descending-neuron cell bodies and neurites in *Mip2M>Mip-UTRs* animals were similar to the wild-type pattern (Figure 5F). Furthermore, restricted expression of this new construct in Mip2M neurons was sufficient to rescue glue retention to the same levels of controls (Figure 5E) (33 out of 40 larvae expelled glue, 82.5%, *P* < 0.00001, Fisher’s exact test with Bonferroni correction, as compared to 0% for the respective control condition). Taken together, these results are strongly consistent with the model that Mip is required in the descending Mip2M neurons for proper GSB.

### Descending Mip2M interneuron activity is critical for GSB initiation

We next asked if Mip2M neurons played additional roles in GSB regulation beyond being the source of neuromodulatory Mip. To test this, we monitored GSB execution in animals where Mip2M neuronal activity was suppressed by inhibiting synaptic transmission neurogenetically. This was achieved using the GAL4-UAS system to express the light chain of the tetanus toxin (TNT) in Mip2M neurons (*Mip2M>TNT*). TNT inhibits vesicle-membrane fusion events by cleaving members of the vesicle-associated membrane protein (VAMP)/synaptobrevin family, such as presynaptic VAMP2, which is critical for synaptic transmission.

Expression of an inactive form of *TNT* (*TNT(-)*), served as negative control for TNT (Sweeney *et al*., 1995). Results showed that animals carrying *Mip2M>TNT* initiate GSB, but do so with an abnormal pattern of movements in 15 out of 23 (65.2%) individuals (*P* < 0.001, Fisher’s exact test) (Figure 6A). Not surprisingly, a similar percentage of *Mip2M>TNT* animals [45 out of 147 (67.3%)] did not expel glue (*P* < 0.0001, Fisher’s exact test) (Figure 6B), reminiscent of *Mip* mutants. In contrast, 9 out of 9 (100%) of *Mip2M>TNT(-)* control animals performed a normal GSB and 54 out of 67 (80.6%) expelled glue, similar to wild-type animal levels (Figure 6A-B). These results suggest that vesicle-membrane fusion events mediated by TNT-sensitive VAMP/synaptobrevin-like proteins are critical for proper GSB execution. Whereas TNT is typically used as a means to block synaptic transmission (Sweeney *et al*., 1995), its effects might be broader as there are other relevant cellular processes that might be sensitive to it. For instance, TNT might also directly negatively affect the secretion of neuropeptides such as Mip, which often occurs from dense core vesicles throughout the neuronal body via VAMP-mediated membrane fusion events (Hoogstraaten *et al.,* 2020). Hence, as both synaptic transmission and dense core vesicle secretion can be compromised by TNT expression, and the extent of the effect obtained is similar to that observed when silencing or mutating *Mip*, these results would be consistent with a single function for Mip2M neurons in GSB: to secrete Mip.

**Figure 6.**
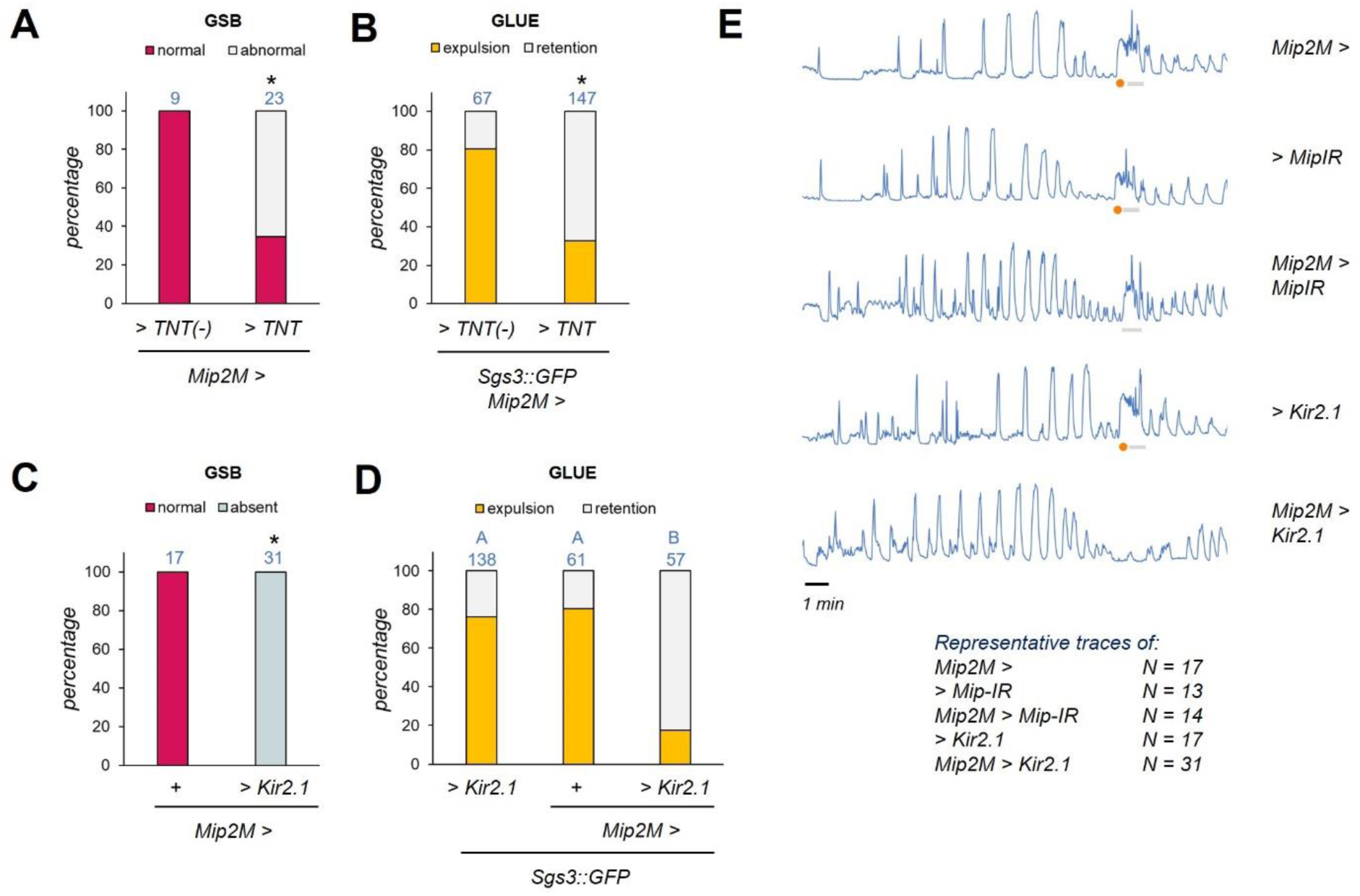
Silencing Mip descending neuron activity blocks GSB. A) Silencing of Mip2M neuronal activity with tetanus toxin (TNT) altered GSB execution. An inactive version of TNT [TNT(-)] served as control. Shown is the percentage of animals that perform a normal GSB. Blue numbers, N. Asterisk, *P = 0.001*, Fisher’s exact test B)Suppression of Mip2M neuronal activity with tetanus toxin (TNT) altered glue expulsion. An inactive version ot TNT served as control. Shown is the percentage of animals that expelled glue (marked by Sgs3::GFP). Blue numbers, N. Asterisk, *P < 0.0001*, Fisher’s exact test C) Silencing Mip2M neurons with the hyperpolarizing Kir2.1 channel completely blocked GSB. Shown is the percentage of larvae of indicated genotypes (same as in A) that perform a normal GSB. Blue numbers, N. Asterisk, *P <* 0.00001, Fisher’s exact test. D) Silencing Mip2M neurons with the hyperpolarizing Kir2.1 channel impaired glue expulsion. Shown is the percentage of larvae of indicated genotypes that expel glue (labeled by Sgs3::GFP). Blue numbers, N. Same blue letter, *P* > 0.016, pairwise Fisher’s exact test with Bonferroni correction (alpha = 0.05, 3 tests). E) Representative traces of muscle GCaMP fluctuations of the indicated genotypes. GSB is labeled with a grey horizontal bar. A strong and sustained increase in fluorescent signal marks the initiation of GSB and is the result of the tetanic contraction that expels the glue out of salivary glands (orange dot). GSB ends with one or two strong peaks that correspond to head waving/whole body contractions. All these elements are present in control conditions. *Mip2M>MipIR* animals execute an abnormal GSB that lacks the tetanic contraction. *Mip2M>Kir2.1* larvae do not show the characteristic pattern of GSB. Time scale bar, 1 min.

If the above were true, silencing Mip2M neurons by other means, such as by membrane hyperpolarization, would also induce similar results: a strong, yet incompletely prevalent disruption of proper GSB execution and glue expulsion. To test this, we used the GAL4-UAS system to express the inwardly rectifying potassium channel Kir2.1 in Mip2M neurons (*Mip2M>Kir2.1*). Kir2.1 expression hyperpolarizes cell membranes, which directly reduces neuronal excitability (Owal *et al.,* 2015), and hence prevents neuronal communication through chemical and electrical synapses. In contrast to TNT expression and *Mip* mutation or RNAi, which disturb GSB yet do not block its initiation, *Kir2.1* expression in Mip2M neurons completely blocked the GSB stage (0 out of 31 larvae did GSB, *P* < 0.00001, Fisher’s exact test, while 17 out of 17 *UAS-Kir2.1* controls did a normal GSB) (Figure 6C). These results suggest that Mip2M descending neurons play an additional and critical role in GSB beyond secreting Mip.

To further characterize and investigate these roles, we followed muscle contraction patterns more closely using a *LexAop2-GCaMP6f* calcium reporter transgene driven by a muscle-specific driver, *mhc-LexA* (*mhc>>GCaMP*), as previously described (Heredia *et al*., 2021). mhc>>GCaMP signals were monitored in animals expressing *Mip2M>Kir2.1* and compared to *Mip2M>Mip-IR* and control animals carrying each transgene alone (*Mip2M>/+, UAS-Mip-IR/+*, or *UAS-Kir2.1/+*). The most salient feature of the mhc>>GCaMP profile associated with GSB in control animals is a sustained increase of the fluorescence intensity that corresponds to the strong tetanic contraction that precedes glue expulsion, and a pair of sharp peaks related to the head waving movements that mark the end of the behavior (Heredia *et al*., 2021) (Figure 6E, top). *Mip2M>Mip-IR* larvae clearly lacked the peak associated with glue expulsion, but apart from that, had a similar profile to control animals, at least at this level of resolution (*N* = 14). In stark contrast, *Mip2M>Kir2.1* animals completely lacked any evidence of muscle contraction between pre- and post-GSB phases (the moment in which GSB is supposed to occur) (*N* = 31) (Figure 6E). As expected, the lack of GSB in *Mip2M>Kir2.1* animals was coupled to strong glue retention. Only 10 out of 57 (17.5%) *Mip2M>Kir2.1* larvae expelled glue (*P* < 0.0001, Fisher’s exact test with Bonferroni correction) compared to 76.1% (105/138) and 80.3% (49/61) of *UAS-Kir2.1* and *Mip2M>* control genotypes, respectively (Figure 6D). Interestingly, however, 17.5% of animals still expelled some glue. As *Mip* mutants showed a complete blockage of glue expulsion, one might speculate that Mip neuropeptide has an additional role on some aspect of glue release that is independent of GSB per se.

In summary, the neuronal silencing experiments described above showed that the inhibition of either vesicle release or neuronal excitability in Mip2M neurons negatively affected GSB, but to different extents. Such differences might be explained by the cell-type-specific susceptibility to manipulation of the underlying molecular processes targeted by each technique (Thum *et al*., 2006; Pauls *et al*., 2015). In this case, considering the hypothesis that both manipulations are completely efficient (a scenario where vesicle release or neuronal excitability are completely blocked by TNT or Kir2.1 expression, respectively), the results indicate that Mip2M neurons have at least two separable functions during pupariation: 1) to activate the GSB program via depolarizations, independently of Mip, and 2) to secrete Mip to modulate the execution of the GSB and glue expulsion programs.

### Descending Mip2M interneuron are command neurons

To test the hypothesis that Mip2M descending interneurons are GSB command neurons, we used an optogenetic approach to depolarize Mip2M neurons of late 3^rd^ instar larvae while monitoring them for GSB-like behaviors in the pupariation arenas. For this, we expressed a channelrhodopsin, *ReaChr*, under the control of *Mip2M>* (*Mip2M>ReaChR*)*. ReaChr* increases membrane potential in response to red or green light stimulation in the presence of all-trans retinal (Inagaki *et al*., 2014). We therefore exposed *Mip2M>ReaChR* larvae previously fed or not 100 µM all-trans retinal to a series of 10x 0.5-s pulses of green (525 nm) light at 1 Hz every 5 min for up to 24 h or until all larvae had pupariated. Importantly, following this exposure regimen, similar amounts of both control and all-trans-retinal-fed larvae pupariated within the experimental timeframe displaying all the normal stages of the motor program: pre-GSB, GSB, and post-GSB. This indicated that neither the all-trans retinal treatment nor the repeated light stimulation regimen altered the progression of the endogenous PMP, allowing us to focus directly on the effects of the light stimuli on behavior. Visual inspection of larvae following the light pulses clearly indicated that GSB-like behaviors were induced exclusively in *Mip2M>ReaChR* larvae that were fed all-trans retinal, but not in unfed animals (Supplementary video 2). The elicited behavior typically occurred ∼10 s after the light-pulse series went off and frequently consisted of a tetanic contraction followed by repeated (two or more) cycles of back-and-forth peristaltic waves. Apart from indicating that Mip2M neurons indeed play a fundamental command role on GSB, this experiment also suggested that the behavior initiation needs the accumulation of a slow-acting second messenger in Mip2M or downstream neurons.

Visual inspection of the larval recordings also indicated that not all light pulses induced GSB-like behaviors. One hypothesis is that the competence of the Mip2M neurons to respond to optogenetic stimuli could be developmentally regulated. The fact that the optogenetic procedure did not interfere with normal PMP progression meant that we could test the effects of developmental time on the competence of Mip2M neurons to generate GSB-like behaviors upon optogenetic stimulation. This was done by synchronizing each animal at a precise PMP stage (namely, the beginning of the endogenous post-GSB stage, see methods) and individually backtracking the analysis of the response of the animal to the light stimuli at specific developmental intervals; specifically, between 3-2 h before post-GSB, 1-0 h before post-GSB, and during pre-GSB (an ∼10-min interval). This strategy allowed us to distinguish between two hypothetical optogenetic-response scenarios: I) where depolarization of Mip2M neurons at any developmental timepoint would be sufficient to elicit GSB, or II) where Mip2M neurons would only become licensed to trigger GSB closer to the time of the endogenous GSB. Quantitative analysis of the results showed that between 3-2 h before post-GSB, only half of the larvae fed all-trans retinal (8 out of 16) responded to the light stimuli by performing a GSB-like behavior, while the rest showed no response (Figure 7A-B). Furthermore, those that responded, did so only to a fraction of the stimuli [10.6% ± 12.7% (10-40%); mean ± SD (Range) of total light stimuli, *N* = 16 larvae] (Figure 7B). In contrast, between 1-0 h before post-GSB, all larvae (20 out of 20) responded to at least one of the light stimuli (Figure 7A), and the average percentage of positive responses significantly increased to 57.3% ± 29.6% (*N* = 20 larvae, P = 0.0016, Dunn’s test against “3-2 h before post-GSB”) (Figure 7B). Critically, in the absence of all-trans retinal, 0/139 light stimuli (*N* = 11 larvae at “1-0 h before post-GSB”) elicited this type of response, strongly indicating specificity of the optogenetically-induced GSB-like response (Figure 7A). Finally, during the short ∼10-min pre-GSB interval, which directly precedes endogenous GSB, all 20 out of 20 larvae surveyed also responded to the optogenetic stimuli and the percentage of positive responses further increased to 93.3% ± 14.7% (*N* = 20 larvae, P = 0.0068, Dunn’s test against “3-2 h or 1-0 h before post-GSB”) (Figure 7B). These results are highly consistent with the second hypothetical scenario raised above, demonstrating that the ability of optogenetically-induced Mip2M neuron depolarizations to evoke a GSB-like response increases as the animal approaches the time of its endogenous GSB, indicating that the competence of Mip2M neurons to act as GSB command neurons is developmentally regulated (Figure 7A). Interestingly, none of the 11 light pulses given to different larvae after the execution of the endogenous GSB produced a GSB-like behavior (Figure 7A), suggesting that the GSB command neuron competence is lost during post-GSB, and hence it is tightly restricted to a specific developmental window.

**Figure 7.**
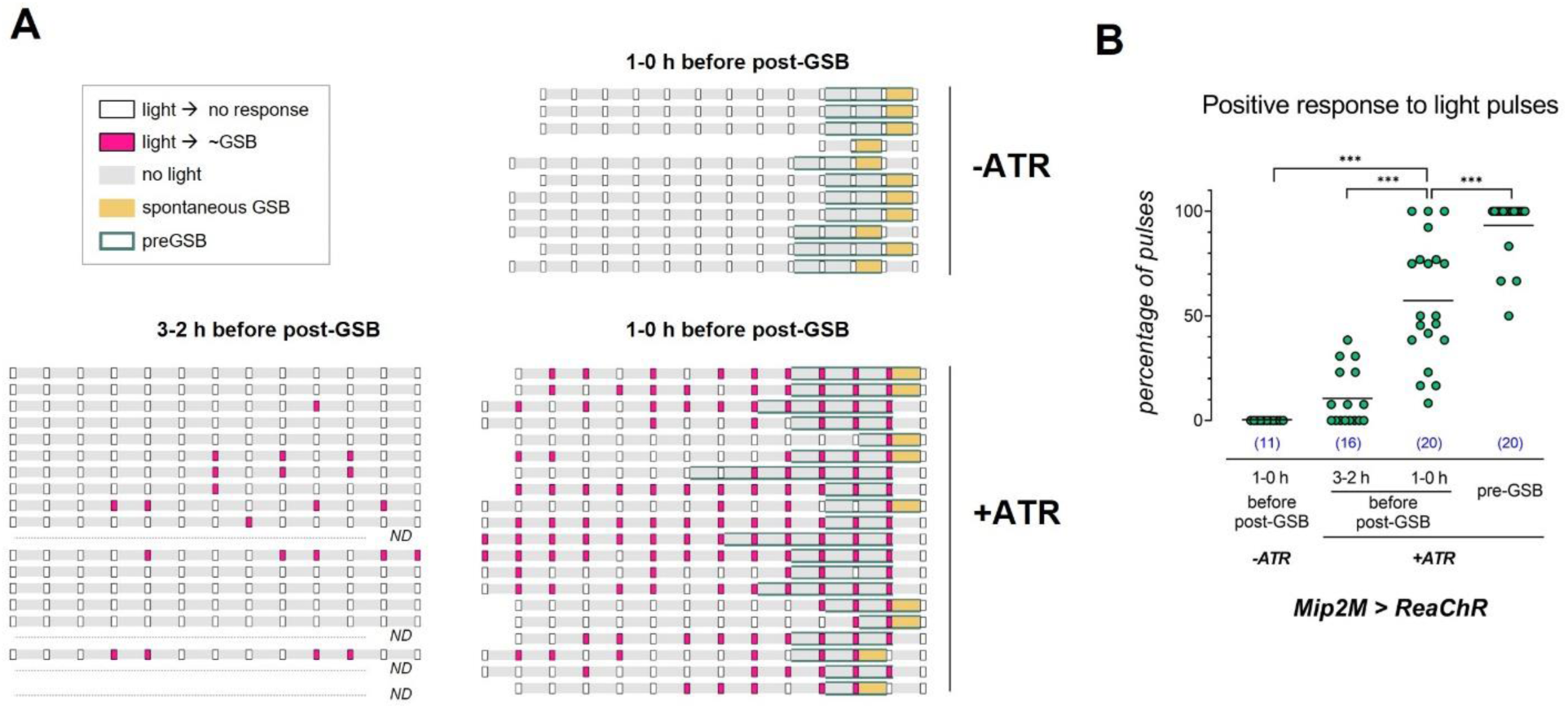
Optogenetic stimulation of Mip2M neurons induces GSB-like behaviors during a specific developmental window. A) Wandering larvae fed with 100 µM all-trans retinal (ATR) were stimulated every 5 min with 10 pulses of green light of 0.5 s of duration at 1 Hz. Response to light was evaluated off-line in 1-h time windows beginning 180 and 60 min before post-GSB (“3-2 h before post-GSB” and “1-0 h before post-GSB” respectively). Rectangular boxes represent light stimuli, filled with white color if the larva did not respond to light and pink if the stimulation induced a GSB-like response. The period when larvae performed pre-GSB contractions is shaded in dark green. 5-min intervals without light stimulation are represented in gray, or in orange when larvae executed a spontaneous GSB, which was interpreted as the “endogenous” (non-light-induced) GSB. Larvae that were not fed with ATR did not respond to light (-ATR). On the other hand, ATR supplementation conferred responsivity to light. The frequency of positive responses to stimuli increased as the larvae approached GSB. Larvae became unresponsive to light as soon as post-GSB was initiated. ND, not done. B) Quantification of the percentage of light pulses that were followed by a GSB-like behavior per larva, without (-ATR) and supplemented with all-trans retinal (+ATR), at the indicated time periods.

Given that the GSB-like behavior is not elicited simultaneously with light stimuli or immediately after the light turns off, and despite the absence of such elicited behaviors in all-trans retinal-negative controls, it could be argued that the behavior observed a few seconds later is merely the fortuitous occurrence of a spontaneous GSB shortly after light stimuli. Against this argument is the fact that endogenous GSB is a “once-in-a-lifetime” behavior under normal circumstances (Heredia *et al*., 2021 and Figure 7A), but when *Mip2M* neurons were experimentally manipulated, most larvae repeatedly executed GSB-like motions in response to light. Furthermore, shortly after performing a light-induced GSB-like movement, 9 out of 20 larvae exhibited the spontaneous GSB (Figure 7A and Supplementary video 3).

Altogether, these results indicate that Mip2M descending neurons are not only necessary, but also sufficient for GSB induction, thus fulfilling the requirements to be considered command neurons. Continuous optogenetic depolarization of Mip2M neurons is not required for the complete execution of the behavior. Instead, it initiates a behavioral response that outlasts its own activity, acting as trigger neurons (Marder *et al*., 2005).

### The Mip receptor, SPR, is critical for proper GSB modulation

To further understand how Mip modulates behavior during the PMP, we examined the effect of its known receptor, SPR (Kim *et al.,* 2010, Poels *et al.,* 2010), on GSB. SPR was initially described as the receptor for the Sex Peptide (SP), a secreted peptide that is expressed only in the male accessory gland (Chen *et al*., 1988). SP is transferred during copulation to the female in seminal fluid and induces postmating changes in reproductive and feeding behavior (Yapici *et al*., 2008; Chapman *et al*., 2003; Liu *et al*., 2003; Carvalho *et al*., 2006). However, SPR is also expressed in the larval and adult male brain (Yapici *et al*., 2008), as well as in other insect species that lack a clear ortholog of SP (Kim *et al*., 2010). This indicated that SPR should have another ligand and led to the identification of Mip as the ancestral ligand for the SPR (Kim *et al*., 2010; Poels *et al*., 2010). Here, we used the deficiency *Df(1)Exel6234* that which removes *SPR* and four other genes (hereafter called *SPR(-/-)*; Yapici *et al*., 2008), as well as RNAi-mediated *SPR* knockdown [*UAS-SPR-IR* (*GD3236*) expressed together with *Dcr2* (hereafter, *>SPR-IR>Dcr2*) (Dietzl *et al*., 2007)] to evaluate the role of SPR in GSB. Consistent with our previous observations reducing *Mip* function, animals homozygous for *SPR(-/-)* also performed an abnormal GSB (0 out of 51 animals tested did a normal GSB, *P* < 0.00001, Fisher’s exact test) (Figure 8A), were unable to expel glue (0 out of 12, expelled glue, *P* < 0.00001, Fisher’s exact test) (Figure 8B), and had a puparium with no obvious morphological defects (Supplementary Figure 4). As expected, larvae lacking the other SPR ligand, SP [*SP^Δ130^* (*SP(-/-)*) (Liu and Kubli, 2003)] performed GSB normally (100% of *SP(-/-)* and background control animals did a normal GSB, *N* = 5 for both conditions) (Figure 8A). These results strongly suggest that Mip acts via its known receptor, SPR, during GSB.

**Figure 8.**
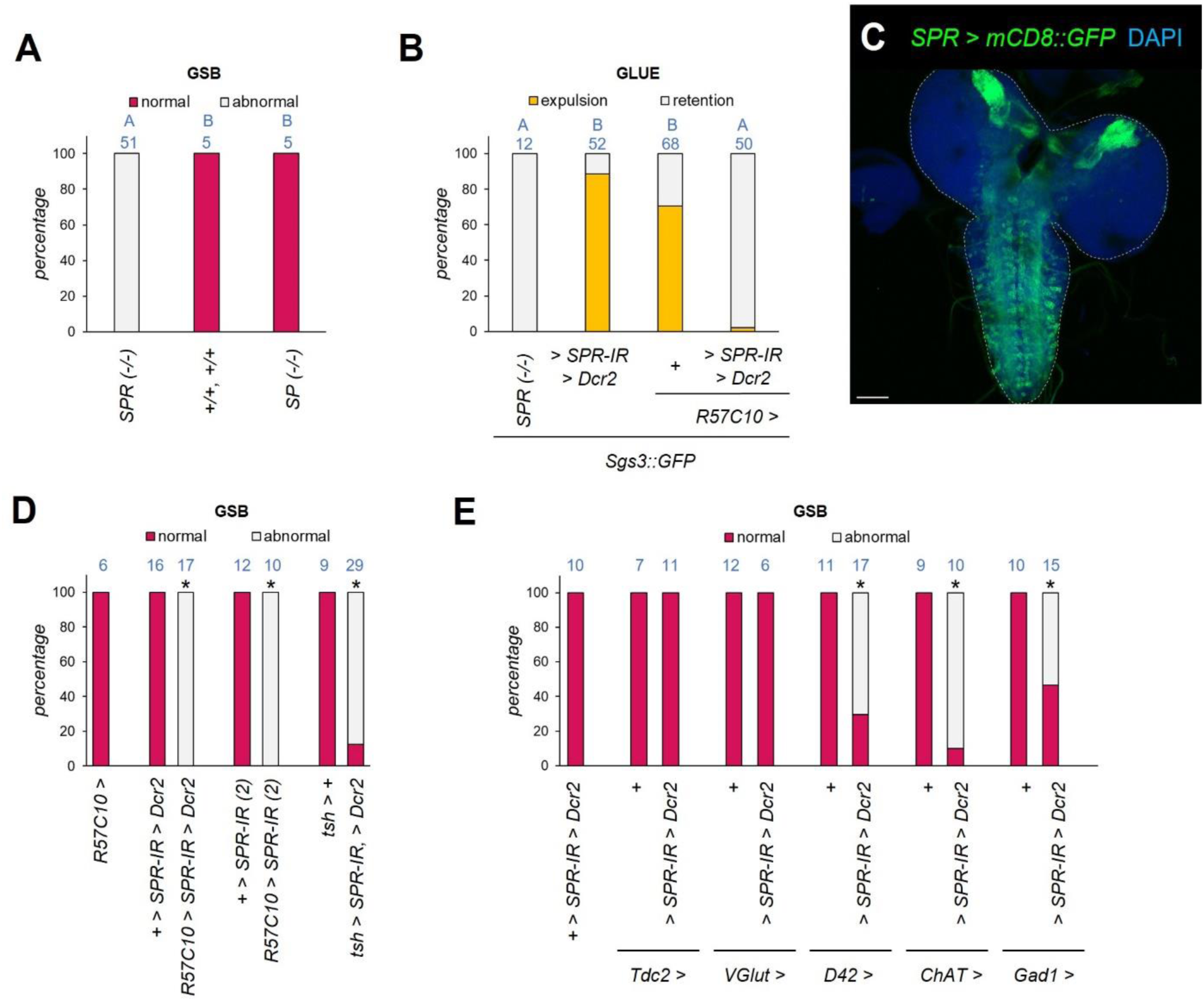
Sex peptide receptor is required in multiple VNC neurons for proper GSB execution. A) The *sex peptide receptor* (*SPR)* gene is required for proper GSB, while the *sex peptide* (*SP*) gene is not. Shown is the percentage of animals of the depicted genotypes [*SPR (-/-): Df(1)Exel6234. SP (-/-): Δ^130^*;(+/+, +/+), *SP[Δ^130^]* background controls] that perform a normal GSB. Blue numbers, N. Same blue letter, *P* > 0.016, pairwise Fisher’s exact test with Bonferroni correction (alpha = 0.05, 3 tests). B) Panneuornal RNAi knockdown of *SPR* (*R57C10*>*SPR-IR>Dcr2*) abrogates glue expulsion. Shown is the percentage of larvae of the depicted genotypes that expelled glue (marked by Sgs3::GFP). Blue numbers, N. Same blue letter, *P* > 0.008, pairwise Fisher’s exact test with Bonferroni correction (alpha = 0.05, 6 tests). C) Confocal stack projections of dissected L3 larval CNS from *SPR>mCD8::GFP* animals. DAPI counterstain, nuclei (blue). Dotted line, CNS contour. Scale bar, 50 µm. D) Pan-neuronal (*57C10>*) or VNC-specific (*tsh>*) RNAi knockdown of *SPR* alters the execution of GSB. The graph shows the percentage of animals of the indicated genotypes that execute a normal GSB. Two different *SPR* RNAi constructs were tested (>*SPR-IR>Dcr2* or >*SPR-IR (2)*). Blue numbers, N. Asterisk, *P* < 0.0001 against corresponding control conditions, pairwise Fisher’s exact test with Bonferroni correction (alpha = 0.05, 3 tests). E) *SPR* is required in multiple neuronal subtypes for GSB. Shown is the percentage of animals that execute a normal or abnormal GSB upon *SPR* silencing (>*SPR-IR>Dcr2*) in octopaminergic (*Tdc2>*), glutamatergic (*VGlut>*), motor (*D42>*), cholinergic (*ChAT>*) and GABAergic (*Gad1>*) neurons. Blue numbers, N. Asterisk, *P* < 0.0001 against corresponding control conditions, pairwise Fisher’s exact test with Bonferroni correction (alpha = 0.05, 3 tests).

### SPR is required in VNC neurons for proper GSB modulation

Transcriptomic studies show that in larvae, *SPR* is weakly expressed in the gut and CNS alone (FlyAtlas, Chintapalli *et al*., 2007). To confirm this and learn more about its expression pattern we crossed an *SPR-GAL4* knock-in where *GAL4* is expressed under the control of *SPR* endogenous regulatory sequences (Deng *et al*., 2019), with a *20x-UAS-IVS-mCD8::GFP* (mCD8::GFP) (Pfeiffer *et al*., 2010) reporter and dissected larval brains. We find strong *SPR>mCD8::GFP* expression in the CNS, consistent with transcriptomic data, most notably in the mushroom bodies, subesophageal zone, and VNC (Figure 8C). We thus hypothesized that *SPR* is required in the CNS for proper GSB. To test this, we expressed two different *SPR-IR* lines [either *>SPR-IR>Dcr2* or *UAS-SPR-IR(2)* (*HMC06520*, hereafter >*SPR-IR(2)*) (Ni *et al*., 2011)] in the nervous system using the pan-neuronal GAL4 line *R57C10>*. Pan-neuronal *SPR* knockdown using either RNAi lines compromised the execution of GSB in all animals evaluated (None of the larvae did a normal GSB, *N* = 17 and 10 for *R57C10>SPR-IR>Dcr2* and *R57C10>SPR-IR(2)*, respectively; *P* < 0.0005 for both conditions, Fisher’s exact test with Bonferroni correction) (Figure 8D) and only 1 out of 50 larvae tested expelled glue (2%, *P* < 0.00001, Fisher’s exact test with Bonferroni correction), as expected, and in contrast to the control conditions (88.5% and 70.6% expelled glue, *N* = 52 and 68, for *>SPR-IR>Dcr2* and *R57C10>* controls, respectively) (Figure 8B). We conclude that the Mip receptor, SPR, is required in neurons for proper GSB and glue expulsion.

Considering the morphology of the Mip2M descending neurons, which extend their axons down the entire length of the VNC, we hypothesized that the critical *SPR-*positive neurons for GSB would lie in the VNC, so that knockdown of *SPR* exclusively in the VNC should mimic the effects of *SPR* deficiency. To do this, we expressed *SPR RNAi* restricted to the VNC using *tsh>* (*tsh>SPR-IR>Dcr2*) and monitored GSB. We found out that, while 9 out of 9 (100%) control *tsh>* animals performed a normal GSB, only 3 out of 29 (∼10%) *tsh>SPR-IR>Dcr2* larvae did so (*P* < 0.0001, Fisher’s exact test with Bonferroni correction) (Figure 8D). These data confirm our hypothesis that *SPR* is required in VNC neurons, but since the percentage of animals presenting with abnormal GSB is lower than the one observed for pan-neuronal *SPR* RNAi or *SPR* mutation, these experiments do not formally exclude the possibility that *SPR* is also required in cells from other regions of the nervous system.

### SPR is required in multiple neuronal subtypes in the VNC for proper GSB modulation

We next attempted to further functionally characterize the population of neurons where SPR is required for the proper execution of GSB. Muscles and the motor neurons that innervate them are arranged in a segmental fashion in larvae. There are thirty muscles in each hemisegment innervated in a highly stereotypical pattern by thirty-six motor neurons (reviewed in Kohsaka *et al*., 2012; Clark *et al*., 2018) that provide glutamatergic input to individual muscles (Type Ib motor neurons) or large groups of muscles (type Is motor neurons). In addition, three type II unpaired motor neurons provide octopaminergic innervation to most muscles (Monastirioti *et al*., 1995; Zarin *et al*., 2019). We manipulated these motor neurons using three GAL4 drivers: two commonly used motor-neuron drivers, *D42-GAL4* (*D42>*) and *OK371-GAL4*, and a driver that labels tyraminergic and octopaminergic neurons, *Tdc2-GAL4* (*Tdc2>*) (Cole *et al*., 2005; Vömel *et al*., 2008). *D42>* is inserted upstream of the *Toll6* sequence and is widely expressed in the VNC, including in at least 30 out of 36 motor neurons (Sanyal, 2009), as well as in the brain and in sensory neurons. *OK371-Gal4* is an enhancer trap inserted upstream of *Drosophila vesicular glutamate transporter* and is also known as *VGlut*-*GAL4* (*VGlut>*). It marks glutamatergic neurons in the brain and VNC, also labeling motor neurons (Mahr and Aberle, 2006). Behavioral monitoring of animals where *SPR* was knocked-down in each of these neuronal populations using *>SPR-IR>Dcr2* showed that while *SPR* knockdown in *VGlut>* and *Tdc2>* cells (*VGlut>SPR-IR>Dcr2* and *Tdc2>SPR-IR>Dcr2,* respectively) did not affect the execution of GSB, knockdown of *SPR* in *D42*> cells (*D42>SPR-IR>Dcr2*) negatively affected GSB performance in 12 out of 17 (∼71%) animals monitored (*P* < 0.005, Fisher’s exact test with Bonferroni correction) (Figure 8E). These results suggested that *SPR* is not required in motor neurons themselves, but in other neurons within the *D42>* expression pattern.

Interestingly, RNAi-mediated silencing *SPR* in other neuronal populations, such as GABAergic and cholinergic neurons, using *Glutamic acid decarboxylase 1 (Gad1)-GAL4* (*Gad>*) (Ng *et al*., 2002) and *Choline acetyltransferase (ChAT)-GAL4* (*ChAT*>) (Diao *et al*., 2015), respectively, also impaired GSB performance in a significant fraction of animals (only 7 out of 15 (∼47%) *Gad>SPR-IR(1)>Dcr2* and 1 out of 10 (10%) *ChAT>SPR-IR>Dcr2* animals did a normal GSB, *P* < 0.01 for both conditions, Fisher’s exact test with Bonferroni correction) (Figure 8E). This indicates that Mip-dependent activation of SPR is required in multiple neuronal subtypes in the VNC. One possibility that arises from such a model is that each population of SPR-positive neurons is responsible for a specific action component of GSB in response to Mip.

### High resolution definition of the GSB fixed action pattern

To test the hypothesis above, we recorded larvae expressing *mhc>>GCaMP* under a fluorescence-equipped stereomicroscope and characterized the behavioral routine, or fixed action pattern, executed during GSB. This routine is a complex sequence of contractions that begins with a strong and sustained activation of the muscles in the anterior third of the larva that expels glue, followed by interspersed backward and forward peristaltic waves that gradually reach more posterior segments of the animal. The end of the behavior is marked by a whole body contraction accompanied by a left and right movement of the anterior end of the larva, or “head waving” (Heredia *et al*., 2021).To quantify these GSB subprograms, we first manually classified the *mhc>>GCaMP* contractions of control larvae (*Mip^1^* background control) into one of these categories: tetanic contraction, backward and forward peristaltic movements, whole body contraction, and whole body contraction with head waving (head waving). Backward movement was further classified as short if it stopped before reaching the posterior third of the body or long if they passed that boundary (Figure 9A). Behavioral subunits were color coded and graphically represented in ethograms.

**Figure 9.**
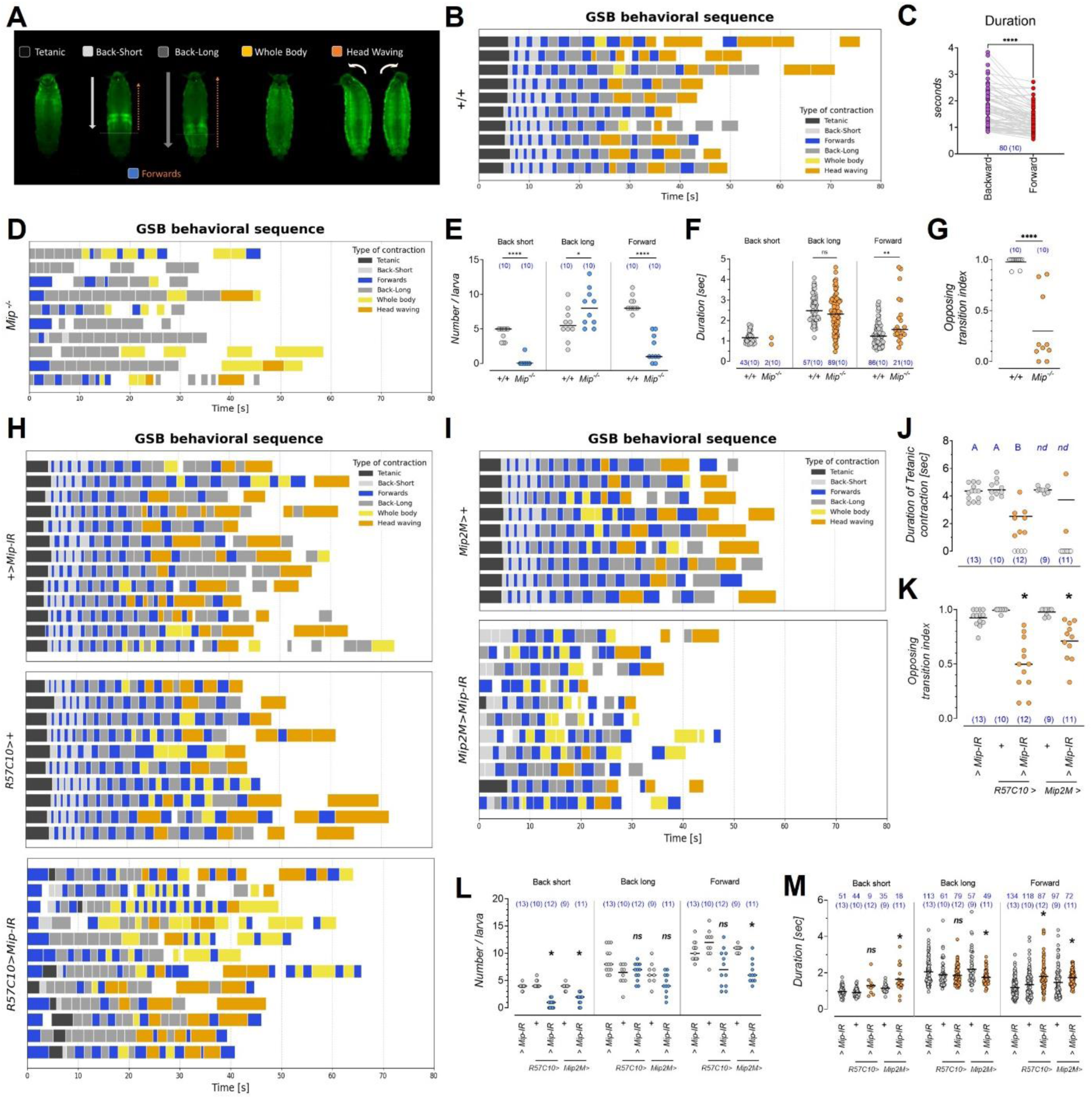
Mip modulates neuronal activity to produce a stereotyped motor pattern during GSB. A) Categories used to classify behavioral subunits that comprise GSB. B) Ethograms displaying the sequence of behaviors and duration of each element of GSB in wild type animals (+/+). Each row represents the color-coded behavioral sequence of one animal expressing *mhc>>GCaMP*. C) Forward peristaltic waves travel the same distance as the preceding backward wave in a shorter time. The graph shows the duration of pairs of backward and forward consecutive peristaltic waves. Dots represents one wave. Blue numbers, N (number of larvae). Asterisk, *P* < 0.0001, Wilcoxon matched-pairs signed rank test. D) Ethogram of *Mip^1^* (*Mip^-/-^*) mutant animals. The initial tetanic contraction is absent, the number of forward peristaltic waves is reduced, and the back- and-forth pattern of peristaltic waves is lost in larvae that lack Mip neuropeptide. E) Quantification of the occurrence of backwards and forwards peristaltic waves in wild type and *Mip^1^* mutant animals (*Mip^-/-^*). Dots represent one larva, Horizontal bar, median value. Blue numbers, number of larvae. ****, *P* < 0.0001, *, *P* < 0.05, Mann Whitney test (for Back short and Forward) and unpaired t-test (Back long). F) Duration of each type of contraction in wild type and *Mip^1^* mutant larvae (*Mip^-/-^*). Each dot represents one contraction. Blue numbers, N (number of larvae). NS, *P* > 0.05, unpaired t-test (Back long). **, *P* = 0.0027, Mann Whitney test (Forward). G) The back-and-forth altern of peristaltic waves is lost in *Mip^1^* mutants, resulting in a low opposing transition index value. See text for description. Dots represents one larva. Blue numbers, N. Horizontal line, mean value. ****, *P* < 0.0001, Mann Whitney test. H-I) Silencing *Mip* expression pan-neuronally or in Mip2M neurons affects GSB behavioral sequence. The initial tetanic contraction is absent or misplaced and the consistency of the alternate back-and-forth peristaltic waves is reduced. Ethograms of *UAS-Mip-IR/+, R57C10>+* and *Mip2M*>+ control conditions and *R57C10>Mip-IR* and *Mip2M>Mip-IR* silenced larvae. J) The duration of the tetanic contraction in those *R57C10>Mip-IR* larvae that show the behavior, is reduced. Dots represents one larva. Blue numbers, N. Horizontal bar, mean value. Empty circles represent animals that did not perform the tetanic contraction and were excluded from the statistical analysis and mean calculation. Same blue letter, *P* > 0.05, ANOVA, Tukey’s HSD test. nd: not determined. K) A reduction of the opposing transition index indicates that the consistency of the back-and-forth transitions is reduced upon pan-neuronal or Mip2M *Mip* knockdown. Blue numbers, N. Asterisk, *P* < 0.01 against respective controls, Kruskal Wallis and Dunn’s test. L) Quantification of the occurrence of backwards and forwards waves in control *R57C10>Mip-IR* and *Mip2M>Mip-IR* silenced larvae. Dots represents one larva. Horizontal bar, median value. Blue numbers, N. Asterisk, *P* < 0.005 against respective controls, Kruskal Wallis and Dunn’s test. M) Duration of each type of contraction in control *R57C10>Mip-IR* and *Mip2M>Mip-IR* silenced larvae. Each dot represents one contraction. Horizontal bar, mean value. Blue numbers, N (number or larvae). Asterisk, *P* < 0.05 against respective controls. Kruskal Wallis and Dunn’s test.

Control animals displayed a highly stereotyped sequence of behaviors that was extremely consistent for the initial 25 seconds and differed only slightly between individuals from that point until completion (Figure 9B, Supplementary video 4). Control animals always initiated GSB with a tetanic contraction that lasted 5.6 ± 0.5 s (mean ± SD, *N* = 10), which was immediately followed by an uninterrupted series of backward and forward peristaltic waves, typically five short and three long backwards movements that gradually reduced their strength with time. Forward waves originated in the same body segment where the preceding backward wave ended. Consequently, the distance traveled by any forward wave is comparable to that of the preceding backward movement, and the speed of propagation of each wave can be considered inversely proportional to its duration. We found that backward waves of muscle contraction propagated on average 23.4% slower than the successive forward counterpart (*N* = 80 cycles observed in 10 control animals; *P* < 0.0001, Wilcoxon matched-pairs signed rank test) (Figure 9C). After the initial 25 s, when whole body and head waving movements occurred, the transition between backward and forward waves became less consistent, and sequences of two backward waves with a head-waving in between them were frequently observed.

However, it is important to note that these sequences of behavior were not expressed with a robust transition pattern and head-wavings could be observed before and after both types of peristaltic waves. A brief pause marked the end of GSB and the initiation of post-GSB. As post-GSB is characterized by whole-body contractions with occasional head-wavings interrupted by periods of muscle relaxation, it can be challenging to ascertain whether a particular contraction is the last one of GSB or the first one of post-GSB. Therefore, we arbitrarily defined the end of GSB as the occurrence of a pause lasting at least 10 s.

### Mip is critical for the correct nature and sequence of GSB component acts

Having determined the fixed action pattern of GSB at a higher resolution, we then first characterized it in *Mip* mutants and then in different *Mip* RNAi knockdown conditions. We found that the GSB fixed action pattern was severely affected by the lack of *Mip*, both in the repertoire of component actions and in their sequential arrangement. Specifically, none of the *Mip^1^* mutant animals evaluated displayed the initial sustained tetanic contraction or the rapid cycles of back-and-forth short peristaltic waves observed in wild-type animals (*N* = 10) (Figure 9D Supplementary video 4). Instead, half of the observed *Mip^1^* mutants initiated GSB with a slow and prolonged forward peristaltic wave, while the rest began with a backwards peristaltic movement that traveled along the entire length of the body. Next, independently of the initial GSB component action used by *Mip^1^* mutants, their GSB proceeded with a prolonged series of long backward peristaltic waves followed by whole-body or head-waving movements, only occasionally interrupted by forward peristaltic contractions. Quantitative analysis showed that short peristaltic waves were absent in 9 out of 10 larvae evaluated (P < 0.0001, Mann Whitney test) and the number of forward waves was also reduced (P < 0.0001, Mann Whitney test) (Figure 9E). The average duration of the long backward peristaltic contractions in *Mip^1^* mutant animals was not different from control animals (*P* = 0.1191, unpaired t-test), but the fewer forward ones lasted longer than in controls (*P* = 0.0027, Mann Whitney test) (Figure 9F). The consistent back-and-forth alternation of peristaltic waves found in control animals was severely affected by *Mip^1^* mutation. We quantified the extent of this effect by assigning a value of 1 to transitions between consecutive peristaltic waves if they moved in opposite directions and 0 otherwise. We then calculated the “opposing transition index” dividing the sum of transition values by the total number of transitions between consecutive peristaltic waves. An index of 1 means there is a perfect back-and-forth alternation and, on the contrary, an index of 0 means that all consecutive waves move in the same direction. Control animals had an index of 0.98 ± 0.05 (mean ± SD, *N* = 10) while *Mip^1^* mutants had 0.30 ± 0.34 (mean ± SD, *N* = 10), indicating a reduction in the alternation between back-and-forth peristaltic waves (*P* < 0.0001, Mann Whitney test) (Figure 9G). Reduced back-and-forth alternation could explain why *Mip^1^* mutants have on average longer forward peristaltic waves (Figure 9F), as the initial short forward waves (reciprocal to the short backward waves) are virtually absent in these mutants, biasing the forward waves towards longer ones. Finally, *Mip^1^* mutants have pauses in the first 20-25 s of GSB, which are not observed in controls. We concluded that Mip modulates different aspects of GSB, throughout the duration of the behavior. Both the elements that are part of it, their duration, and their sequence require the action of Mip to be executed correctly. Of particular note is the finding that *Mip* is absolutely critical for the initial tetanic contraction that is tightly associated with glue expulsion (Heredia *et al.,* 2021). It is likely therefore that this is the critical action component requiring Mip activity for efficient glue expulsion.

### Neuronal Mip determines the nature and sequence of GSB component acts

Next, we analyzed the structure of GSB in animals with pan-neuronal *Mip* RNAi silencing (*R57C10>Mip-IR*). We found that *57C10>Mip-IR* altered the structure of GSB in a similar fashion to *Mip^1^* mutation (Figure 9H,L-M). The most prominent consequence of this was a change in the initiating GSB component act, which instead of the tetanic contraction, was a slow forward peristaltic wave in all but one of the larvae evaluated (1 out of 10 versus 13 out of 13 and 10 out of 10, for *R57C10>Mip-IR* vs *UAS-Mip-IR/+* and *R57C10>+* controls) (Figure 9H). Despite not being the first element of the behavioral sequence, the tetanic contraction was still observed in two thirds of the animals, but executed with a defective timing and duration, lasting on average 65% less than the tetanic contraction of control larvae (*P* < 0.002, ANOVA and Tukey’s HSD test) (Figure 9J). Importantly, albeit present, the tetanic contraction was ineffective in expelling glue (less than one third of *R57C10>Mip-IR* animals expelled glue; see Figure 1H). The number of short backward peristaltic waves was reduced and the duration of forward movements increased (Figure 9L-M). More importantly, another fully penetrant difference with control animals was the lack of the initial highly-organized alternation cycle between short backward and forward peristaltic waves. Instead, and similarly to *Mip^1^* mutant animals, *R57C10>Mip-IR* animals immediately executed the long backward peristaltic waves organized in stretches of self-transitions or transiently interrupted by waves in the opposite direction. The loss of the alternation of back-and-forth peristaltic waves was reflected by a significantly lower opposing transition index (0.50 ± 0.24; mean ± SD, *N* = 12) than controls (0.92 ± 0.10 and 0.99 ± 0.01 for *>Mip-IR* and *R57C10>*; *P* = 0.0268 and P < 0.0001, respectively, Dunn’s test) (Figure 9K). Apart from indicating that neuronal Mip modulates both the components and the sequence of GSB throughout its duration, these results also implicate neuronal Mip as a critical factor in the initial tetanic contraction. The slight difference between the effect of *Mip* mutation and pan-neuronal *Mip* RNAi on the tetanic contraction [fully penetrant absence vs. fully penetrant problems (absence or abnormality), respectively] could be explained by the nature of the *Mip* loss of function manipulations (knockout x knockdown).

### Mip from descending Mip2M interneurons determines the nature and sequence of GSB component acts

Silencing of *Mip* in *Mip2M* neurons (*Mip2M>Mip-IR*) produced the same type of alterations as ubiquitous neuronal silencing; lack of the tetanic contraction and of the short-range alternating back-and-forth peristaltic waves. The initial tetanic contraction was missing in 82% of *Mip-2M>Mip-IR* larvae (9 out of 11) and the number of peristaltic waves, both in the backward and forward direction, was significantly reduced (Figure 9I). Self-transitions between backward peristaltic waves present in *Mip^1^* and *57C10>MipIR* individuals were also found in *Mip2M>Mip-IR* animals but, in this case, in short stretches of only two or three consecutive events (8 out of 11 larvae), with an average opposing transition index of 0.71 ± 0.18 (mean ± SD, *N* = 11) (Figure 9K). As we had previously concluded that the large descending Mip2M interneurons are the critical neurons modulating GSB (Figures 2B and 3G-I), these results are strongly consistent with the model where Mip produced and released from the descending Mip2M interneurons is critical for the presence and correct sequence of different GSB action components throughout its duration. Moreover, it strongly implicates the descending Mip2M interneurons in the modulation of the initial tetanic contraction that is tightly linked to glue expulsion. If this model were correct, then the pan-neuronal expression of *Mip* in *Mip^1^* mutant animals—a condition that rescued glue retention (Figure 5E)–should rescue the nature and sequence of the GSB fixed action pattern. To test this, we monitored mhc>>GCaMP in “*Mip^1^, R57C10>Mip*”-rescued mutant animals at high resolution as described above. We found that, even though there were some quantitative differences in the number and/or duration of the GSB action components, all of them were restored and in their correct (“wild type”) order in genetically-rescued *Mip^1^, 57C10>Mip* animals (Figure 10). For example, despite the tetanic contraction being 60% shorter than that of wild-type individuals (*P* = 0.0001, Mann Whitney test), its presence (in contrast to its complete absence in *Mip^1^* mutant larvae) was fully rescued by pan-neuronal expression of *Mip* (Figure 10A), highly consistent with our previous finding that animals of this genotype efficiently expulse glue (Figure 5E). Glue expulsion was followed by peristaltic waves in both directions that began as one or two short range backwards movements (5 out of 7 animals) and then transitioned to long range contractions with a robust alternate pattern of back-and-forth changes in direction and an average opposing transition index of 0.97 ± 0.07 (mean ± SD, *N* = 7, *P* = 0.0057 and *P* = 0.0005 against *Mip^1^, UAS-Mip/+* and *Mip^1^, R57C10>+*, respectively, Dunn’s test) (Figure 10C). Hence, we conclude that widespread expression of *Mip* within the nervous system rescued the overall profile of GSB. The subtle differences with the wild-type sequence suggest that the smooth transition through the stages might be the result of gradual or locally regulated release of the neuropeptide that was not reproduced in our experimental (pan-neuronal) rescue approach.

**Figure 10.**
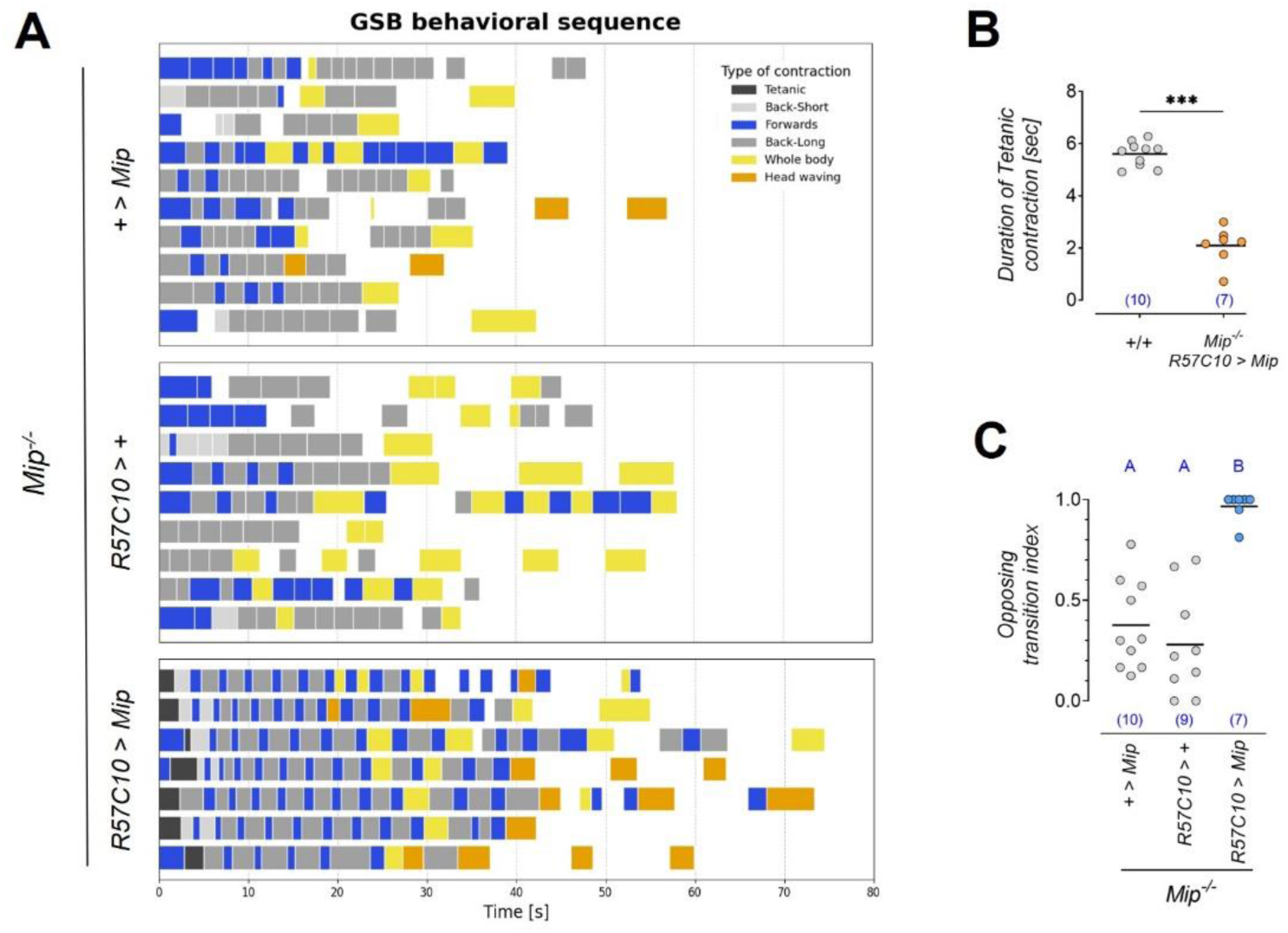
Pan-neuronal expression of Mip rescues GSB behavioral sequence. A) Ethograms displaying the sequence and duration of GSB component acts in *Mip^-/-^, >Mip/+* and *Mip^-/-^, R57C10>+* mutant controls and when *Mip* expression is rescued in the whole CNS in *Mip^-/-^ R57C10>Mip* animals. Pan-neuronal expression of *Mip* rescues the tetanic contraction and the alternate back and forth peristaltic waves. Each row represents the color-coded behavioral sequence of one animal expressing *mhc>>GCaMP*. B) Pan-neuronal expression of *Mip* rescues tetanic contraction presence, but not its duration, relative to wild-type control levels (+/+). Shown are dot plots, where each dot represents one larva of the indicated genotypes. Horizontal bar, mean value. Blue number between parenthesis, N larvae. Asterisk, *P* = 0.0001, Mann Whitney test. C) The alternating pattern of back-and-forth peristaltic waves (opposing transition index) is rescued in *Mip^-/-^, R57C10>Mip* larvae. Shown are dot plots, where each dot represents one larva of the indicated genotypes. Horizontal bar, mean value. Blue number between parenthesis, N larvae. Same blue letters, *P* > 0.05, Kruskal Wallis and Dunn’s test.

### Interaction between different SPR+ neuronal populations determines specific GSB component acts and their sequences

If Mip2M-neuron-derived Mip is required to specify multiple GSB action components, we expect that the silencing of SPR should have the same effects. Furthermore, we hypothesized that SPR activity controls different components in the different neuronal populations we have identified above. To test this, we first characterized the GSB profile of animals with pan-neuronally silenced *SPR*. We found out that, similarly to *Mip* mutant or *Mip* RNAi animals, *R57C10>SPR-IR>Dcr2* animals lacked the initial tetanic contraction of GSB and consisted mostly in backward peristaltic waves terminated by whole body contractions or head waving movements (Figure 11A), with only a few forward waves poorly organized in a back-and-forth pattern, giving an average opposing transition index of 0.49 ± 0.27 (mean ± SD, *N* = 7, *P* < 0.0001 and *P* = 0.0012 against *R57C10>+* and *>SPR-IR>Dcr2/+* controls, respectively, Dunn’s test) (Figure 11B). SPR silencing also reduced the number of peristaltic movements (Figure 11C) and increased their length (Figure 11D). These results corroborate the model where Mip acts via neuronal SPR to regulate multiple GSB component acts and their sequences.

**Figure 11.**
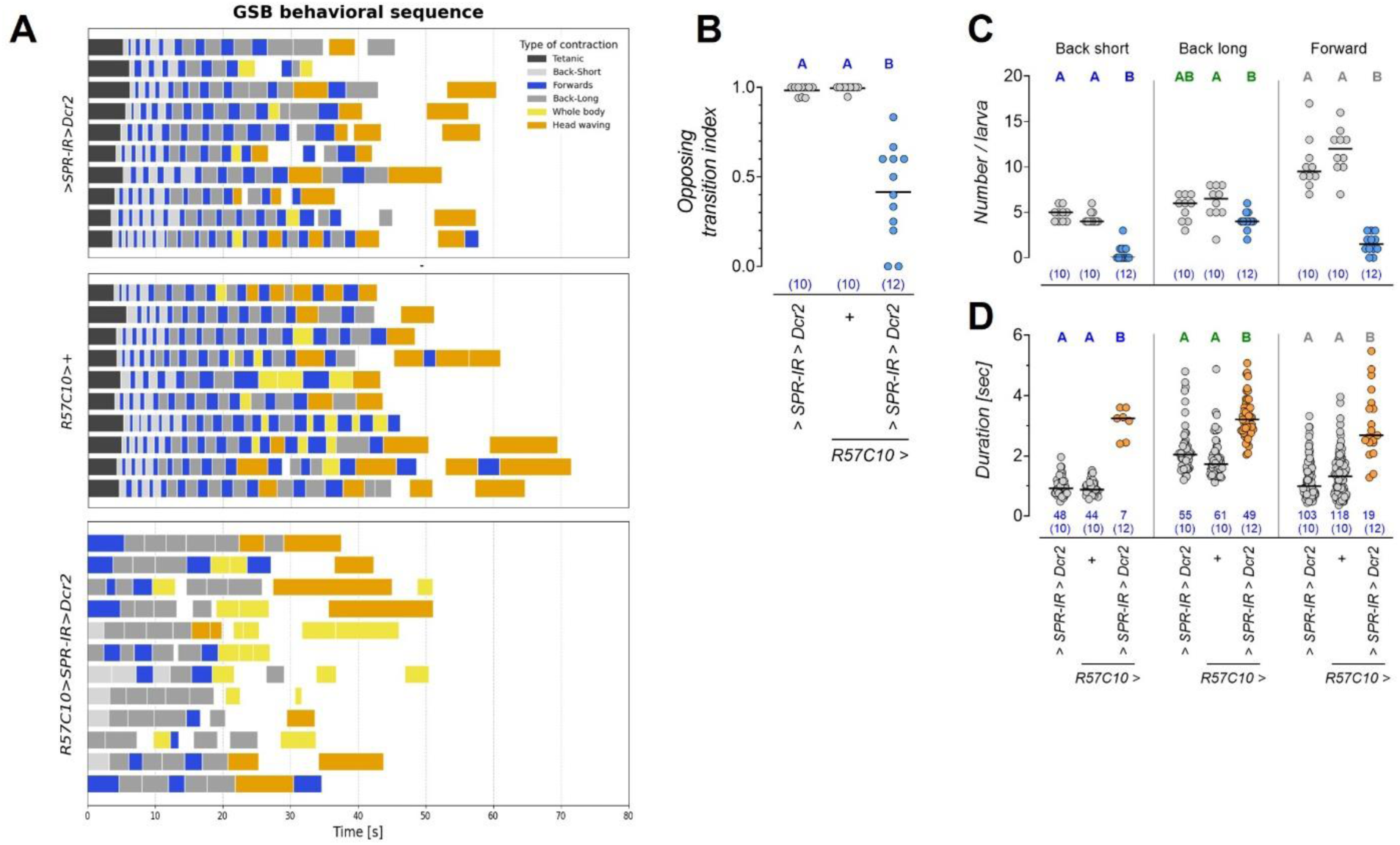
Pan-neuronal silencing of SPR altered the pattern of GSB components. A) Ethograms displaying the sequence and duration of GSB component acts in *UAS-SPR-IR/+* UAS-Dcr2/+ and *R57C10>+* controls and when the SPR is pan-neuronally silenced with *R57C10>SPR-IR>Dcr2.* Pan-neuronal knockdown of SPR results in the same type of alterations than *Mip* mutation. The *R57C10>+* control dataset corresponds to the same dataset analyzed in Figure 9. B) Pan-neuronal silencing of *SPR* disorganizes the pattern of peristaltic waves and reduces the opposing transition index. Each dot represents on larva. Horizontal bar, mean value. Blue numbers, N. Same blue letters, *P* > 0.05, Kruskal Wallis and Dunn’s test. C) Quantification of the occurrence of peristaltic wave movements in the indicated genotypes. Silencing of *SPR* reduces the number of backward short and forward peristaltic waves. Each dot represents one larva. Horizontal bar, median value. Blue numbers, N. Same letters, *P* > 0.05, Kruskal Wallis and Dunn’s test. D) Quantification of the duration of peristaltic wave movements in the indicated genotypes. Silencing of *SPR* increases the duration of all types of movements. Horizontal bar, mean value. Blue numbers: number of contractions and number of larvae between parentheses. Same letters, *P* > 0.05, Kruskal Wallis and Dunn’s test.

Next, we analyzed the structure of GSB when *SPR* was silenced in non-overlapping subpopulations of neurons using *D42>*, *ChAT>*, or *Gad>*, which visibly altered GSB in previous lower resolution analyses in the pupariation monitor, and *VGlut>*, which did not have a visible effect (Figure 8E). Interestingly, all of these manipulations affected particular aspects of GSB, but none of them individually fully phenocopied the effect of pan-neuronal manipulation (Figure 12), and in the case of *D42>*, it even resulted in change in the opposite direction. Specifically, rather than blocking the tetanic contraction, knockdown of *SPR* in *D42>* neurons (*D42>SPR-IR>Dcr2*) doubled its duration from 4.8 ± 1.0 and 4.8 ± 0.5 s, in *+>SPR-IR>Dcr2* and *D42>+* control animals (mean ± SD, *N* = 10 for both genotypes), respectively, to 10.3 ± 1.8 s (mean ± SD, *N* = 11) (*P* < 0.0001, ANOVA and Tukey’s HSD test) (Figure 12E). It also reduced the number of backward and forward waves, while increasing their duration (Figure 12B-C). One possible interpretation of this finding is that SPR is required in D42> neurons for animals to achieve the proper speed of propagation of muscle peristaltic contractions during GSB. Knockdown of SPR in cholinergic and GABAergic neurons with *ChAT>* and *Gad1>* respectively, reduced or completely abolished the number of short range and forward peristaltic waves, increased the duration of the latter and disorganized the alternate back-and-forth pattern leading to self-transitions in both directions (Figure 12F-M). Direct observation of larvae filmed in the pupariation monitor suggested that GSB was not grossly altered by *VGlut>SPR-IR>Dcr2* (Figure 8E). However, this higher resolution analysis with *mhc>>GCaMP* detected subtle deviations from normal behavior, such as the execution of consecutive peristaltic waves, in this case in the forward rather than backward direction, that resulted in an opposing transition index of 0.87 ± 0.11 (mean ± SD, *N* = 16) (*P* = 0.0049 and P = 0.011 against *+>SPR-IR>Dcr2* and *VGlut>+* respectively, Dunn’s test), and a decrease in the duration of the backward contractions and head waving movements (Supplementary Figure 5). As a whole, this data suggests that the sequence of behaviors and their characteristics emerge as the result of the modulation of the interaction between multiple SPR+ neuronal types at the circuit level by Mip neuropeptide derived from the descending axons of the Mip2M interneuron. Also, the data indicates that none of the neuronal populations evaluated herein controls the execution of a single specific component under Mip/SPR signaling by itself, even though we cannot exclude the possibility that this type of control is exerted by other neuronal types not yet assayed.

**Figure 12.**
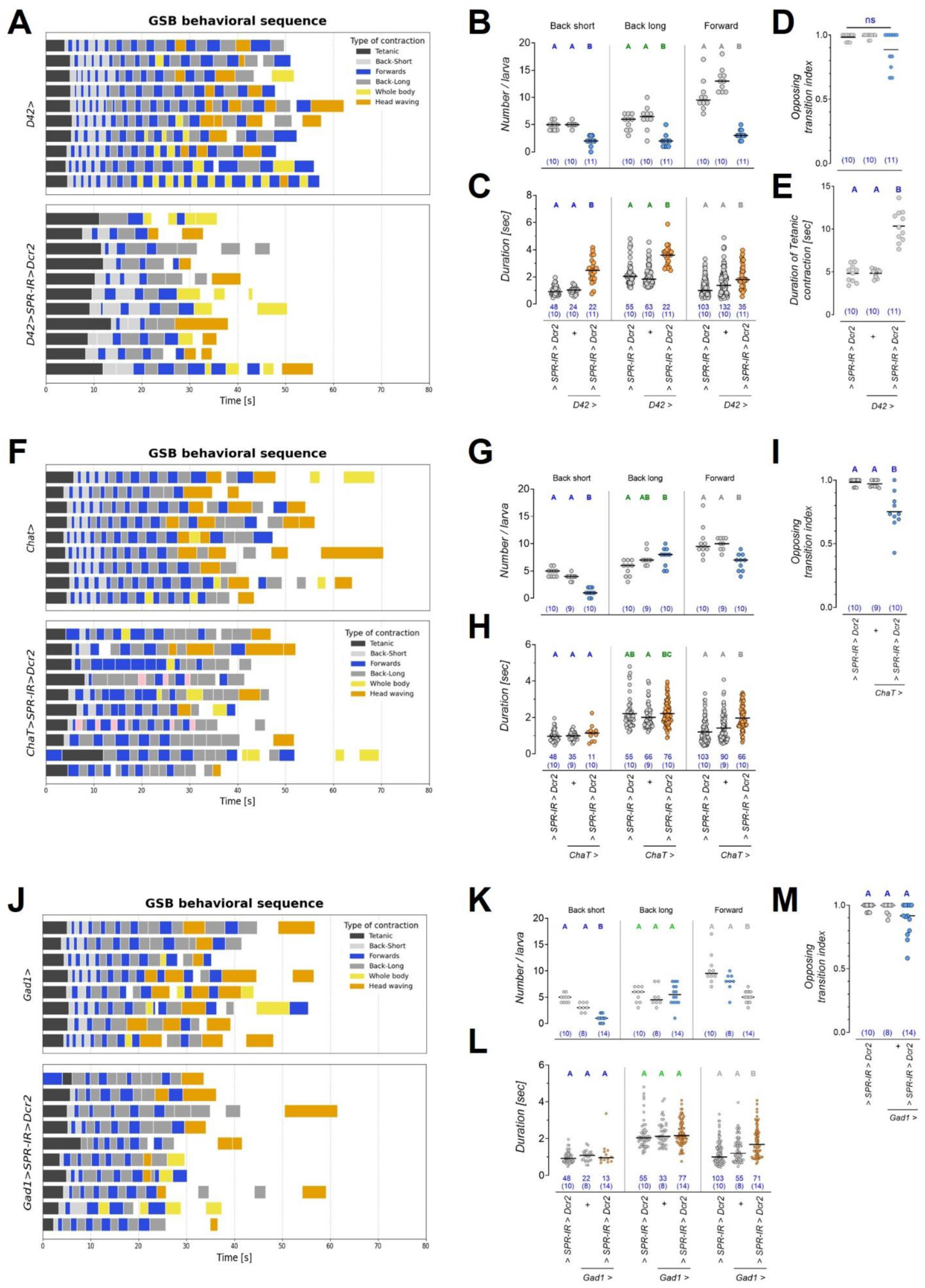
SPR is simultaneously required in multiple neuronal populations to shape GSB. Ethograms displaying the sequence and duration of GSB component acts, and quantification of the occurrence and duration of peristaltic waves and the robustness of the propagation of peristaltic waves in alternat directions with the opposing transition index in motor and sensory neurons (*D42>, A-E)*, cholinergic (*ChAT>, F-I)* and GABAergic (*Gad1>, J-M)* neurons. A) Silencing *SPR* expression with *D42>SPR-IR* extends the tetanic contraction and reduces the number of peristaltic waves. *D42>* labels motor and sensory neurons. These behavioral alterations are attributable to the effect on non-motor neurons, since they are not observed when *SPR* is knockdown with *VGlut>*, a different motor neuron driver (Supplementary figure 5). B) Quantification of the occurrence of backwards and forwards peristaltic waves showing that the number of all types of peristaltic movements is reduced in *D42>SPR-IR* larvae. Each dot is an animal. Horizontal bar, mean value. Blue numbers, N. Same colored letters, *P* > 0.05, Kruskal Wallis and Dunn’s test. C) Peristaltic waves propagate at a lower speed in *D42>SPR-IR* larvae. Each dot is a contraction. Horizontal bar, median value. Blue numbers, N (number of larvae). Same colored letters, *P* > 0.05, Kruskal Wallis and Dunn’s test. D) The pattern of back-and-forth peristaltic waves is preserved upon silencing with D42>. Each dot is an animal. Horizontal bar, mean value. Blue numbers, N. Same letters, *P* > 0.05, Kruskal Wallis and Dunn’s test. E) The tetanic contraction doubles its duration in *D42>SPR-IR* larvae. F) Ethograms of *ChAT>SPR-IR>Dcr2* larvae. Knockdown of *SPR* in cholinergic neurons reduces the number of back-short peristaltic waves and affects the alternate pattern of peristaltic waves. G) The number of backwards short and forward peristaltic waves is reduced upon *SPR* silencing in cholinergic neurons. Each dot is an animal. Horizontal bar, mean value. Blue numbers, N. Same colored letters, *P* > 0.05, Kruskal Wallis and Dunn’s test. H) Forward peristaltic waves last longer in *ChAT>SPR-IR>Dcr2* larvae, mostly as the consequence of the absence of the initial short forward ones. Each dot is a contraction. Horizontal bar, median value. Blue numbers, N (number of larvae). Same colored letters, *P* > 0.05, Kruskal Wallis and Dunn’s test. I) A reduction in the opposing transition index reflects an alteration of the back-and-forth peristaltic pattern when *SPR* is silenced in cholinergic neurons. Each dot is an animal. Horizontal bar, mean value. Blue numbers, N. Same letters, *P* > 0.05, Kruskal Wallis and Dunn’s test. J) Ethograms displaying the behavioral sequence during the execution of GSB in control larvae and when *SPR* is silenced in GABAergic neurons. Knockdown animals do fewer back-short peristaltic waves than controls. K) The number of backward short and forward peristaltic movements is lower in *Gad1>SPR-IR>Dcr2* animals when compared with controls. However, this tendency is not statistically significant. Each dot is an animal. Horizontal bar, mean value. Blue numbers, N. Same colored letters, *P* > 0.05, Kruskal Wallis and Dunn’s test. L) Knockdown of *SPR* in GABAergic neurons does not affect the duration of peristaltic waves. Each dot is a contraction. Horizontal bar, median value. Blue numbers, N (number of larvae). Same colored letters, *P* > 0.05, Kruskal Wallis and Dunn’s test. M) The opposing transition index of *Gad1>SPR-IR>Dcr2* larvae is not statistically different from control animals. Each dot is an animal. Horizontal bar, mean value. Blue numbers, N. Same letters, *P* > 0.05, Kruskal Wallis and Dunn’s test. Dot plots: dots represent one contraction (duration plots) and one larva (rest of the plots), Horizontal bar is median number of contractions, mean duration time and mean opposing transition index. Blue numbers: number of contractions and number of larvae between parentheses. Same letters, *P* > 0.05, Kruskal Wallis and Dunn’s test.

Together, these results indicate that Mip neuropeptide plays a crucial role in shaping the PMP downstream or in parallel to the Dilp8/Lgr3 signaling event that unlocks PMP progression (Figure 13). The results presented above further show that Mip does not act as a triggering mechanism for GSB, but, instead, it is triggered by depolarizing events originating from descending Mip2M neurons. These same neurons secrete Mip, which functions as a neuromodulator of multiple SPR+ neuronal types located in the larval VNC that are essential for both the proper selection of fixed action component acts and their execution in a highly ordered sequence to achieve all the behavioral complexity that comprise GSB and glue expulsion.

**Figure 13.**
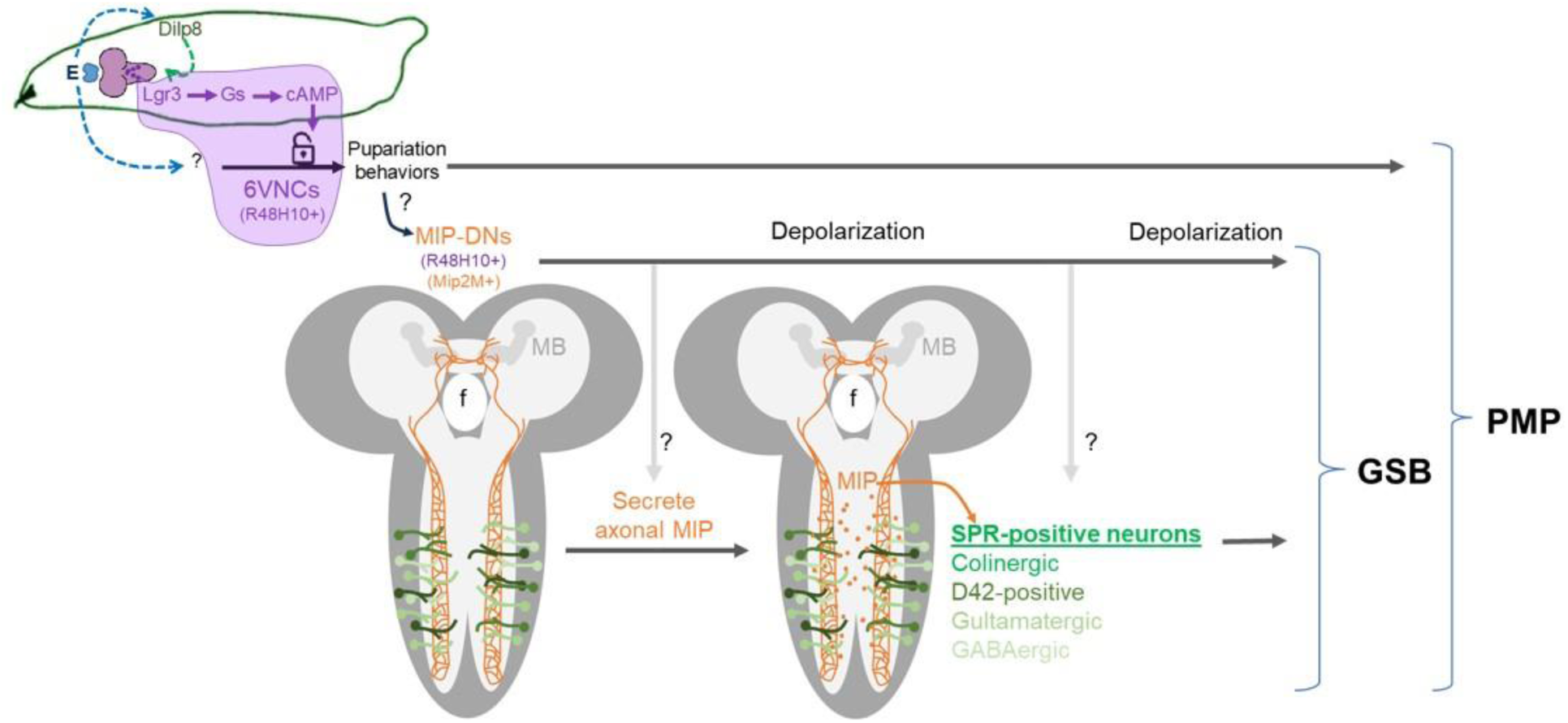
Model of Mip and SPR activity during GSB. A surge in ecdysone (E) in the prothoracic gland of the larva (blue organ in larva) during the end of the 3rd instar larval growth period, leads to Dilp8 production in the larval cuticle epidermis and concomitantly to the initiation of pupariation behaviors. However, Dilp8 signaling to 6 ventral nerve cord (6VNC) neurons positive for Lgr3 (dark magenta cells in the small magenta central nervous system (CNS) in the larva cartoon) is required to unlock the progression of the pupariation motor program beyond the initial short pre-GSB contractions. Importantly GSB, the glue expulsion and spreading behavior, is completely abrogated in the absence of Dilp8 or Lgr3 (Heredia *et al*., 2021). 6VNC neurons are positive for the R48H10 enhancer activity. We identified the Myoinhibiting peptide (Mip) in an RNAi screen in R48H10 cells. However, Mip is required in cells other than 6VNC, namely in two descending neurons (Mip-DNs, orange in enlarged CNS model), which are positive for both R48H10 and Mip2M enhancer activity. Mip is required for the modulation of multiple GSB action components via its SPR receptor, which is expressed and required in multiple populations of VNC neurons, including Cholinergic, D42-positive, Glutamatergic, and GABAergic neurons, for proper modulation of GSB action components. Independently of Mip neuropeptide, depolarization of Mip-DNs (Mip2M neurons) is also absolutely required for the triggering of GSB, albeit GSB is aberrant and restricted to an uncoordinated subset of the GSB components in this situation. In the absence of Mip or Mip-DN neuron activation the rest of the PMP occurs and is mostly normal, although subtle alterations are detected in pre-GSB in Mip mutants, suggesting that Mip might also modulate other PMP components.

## DISCUSSION

We have found that a pair of descending command neurons that reside in the brain lobes and project long axons across the larval VNC acquire competence during a tightly-controlled developmental window to act as command neurons of the GSB stage, a defining stage of the PMP. These descending interneurons express and require the neuropeptide Mip to modulate the activity of SPR+ neurons within the VNC to generate the motor pattern characteristic of the GSB stage.

Innate behaviors consist of a series of highly stereotyped motor actions produced without prior learning and executed until completion once initiated. They rely on the existence of a preconfigured neural circuit that is activated in response to external or internal stimuli. The PMP of *D. melanogaster* is an innate behavior triggered by a surge of the molting hormone Ecdysone at the end of the larval period that, through three stages of patterned muscle contractions, reshapes the larval cuticle into the protective puparium. While the program as a whole is developmentally regulated by Ecdysone, progression through behavioral stages is promoted by the action of the epidermally-derived relaxin hormone Dilp8 on a minimal population of six neurons in the VNC. Other innate behaviors in *Drosophila*, as well as in other species, are also initiated by neuropeptidergic signaling (Dewey *et al*., 2004; *et al.,* 2005; Scheller *et al*., 1982). One well-studied example is the ecdysis behavior of holometabolous insects. Injection of ETH elicits the complete ecdysis routine in *Drosophila*, *Bombyx,* and *Manduca* (Kim *et al*., 2006; Adams *et al*., 1997; Zitnan *et al.,* 1996) by sequentially activating downstream neuropeptidergic neurons that cooperatively shape motor output (Kim *et al*., 2006; 2015). These ETH-responsive neurons express a cocktail of neuropeptides that include kinin, FMRFa, CCAP, Mip, and Bursicon. Silencing the activity of subsets of peptidergic neurons produces behavioral changes, affecting the duration, rhythmicity, or completely abolishing specific stages of ecdysis (Kim *et al*., 2015).

We hypothesized that the complex motor pattern of the PMP would similarly emerge from the modulatory activity of neuropeptides downstream of Dilp8 and identified Mip as one of these signals. This is, to the best of our knowledge, the first role described for Mip in *Drosophila* larva supported by direct genetic evidence. Mip has profound effects on the sequence of behaviors during the GSB stage of the PMP. First, it is an absolute requirement for the inclusion in the repertoire of behaviors of the initial tetanic contraction that expels salivary glue. Secondly, it organizes the series of peristaltic movements in an alternate back-and-forth fashion and, finally, it is required for the transient initial restriction of peristaltic waves to the anterior segments of the larva and for the gradual recruitment of more posterior segments into the behavior. The data presented here is consistent with Mip being required to generate the peristaltic waves that propagate forward. The observation that *Mip-*mutant animals have fewer forward waves, but always intercalated with movements in the opposite direction supports this idea. Nonetheless, since about 30% of *Mip2M>Mip-IR* animals displayed consecutive forward waves, the possibility that Mip might alternatively, or simultaneously, act to facilitate a switch in direction should be considered.

The neural circuits that control larval locomotion have been extensively studied, and many components of the underlying neural circuits that participate in motor neuron coordination within segments and between contiguous segments have been identified (reviewed in Clark *et al*., 2018; Gowda *et al*., 2021). However, the role of most of the 250 interneurons per VNC hemisegment, as well as that of a large number of ascending and descending interneurons that traverse the VNC, and how they integrate in the circuits for executing distinct locomotor behaviors are only now beginning to be understood. One example is the mooncrawler descending neuron (MDN) that simultaneously blocks forward and elicits backward locomotion by activating the A18b premotor neuron, which is active during backward movements, and inhibiting the forward-active A27h premotor neuron via the GABAergic inhibitory neuron Pair1, another descending neuron. Acting this way, the MDN controls the activity of two parallel circuits that ultimately activate the same group of motor neurons, and hence, muscles, to produce antagonistic outputs (Carreira-Rosario *et al.,* 2018). The main source of Mip that modulates GSB is a pair of descending neurons with command activity. These likely represent the large Dimmed(+) Tachykinin(+) Mip(+) descending neuron labeled as #38 by Park (Park *et al*., 2008; Nässel and Winther, 2010). However, Tachykinin does not seem to contribute to the behavior as revealed by a negative result in our screen (Figure 1B) using an RNAi line of proven efficacy (Song *et al*., 2014), at least at the resolution level of the screen. On the other hand, Mip plays a key modulatory role, but it is not itself the trigger of the behavior, evidenced by the timely execution of a defective GSB even in *Mip^1^* null mutant larvae. Descending neurons, like the aforementioned MDN, connect higher order centers with the local circuits in the VNC that produce repetitive action patterns (Guo *et al.,* 2022; Namiki *et al.,* 2022; Suver *et al*., 2016). The data presented here indicates that a similar role is played by the Mip2M descending neuron, with profuse dendritic arborizations in the brain lobes and a long axon that reaches the end of the VNC. Mip2M descending neuron silencing (via Mip2M>Kir2.1 expression) completely abolishes GSB, and the optogenetically-induced depolarization triggers GSB, demonstrating that it exerts a command function on the behavior. Furthermore, we found out that this command function is developmentally regulated. Specifically, the probability of triggering GSB-like behaviors is increased as the animal approaches pupariation and abruptly ends as the animal enters post-GSB. The factor(s) that participate in this command-neuron competence acquisition remain to be identified, but are likely to involve either ecdysone itself, whose effects culminate at pupariation, or ecdysone-dependent factors, such as those dependent on the Dilp8-Lgr3 pathway, which is triggered by ecdysone at pupariation (Heredia *et al.,* 2021). Regardless, our findings on the Mip2M descending neurons offer a well-defined paradigm to study the developmental licensing of command neuron activity.

The Mip2M descending neurons could have multiple targets along the VNC. Co-transmission of neurotransmitters and neuropeptides, as in this case, provides flexibility for circuit modulation (Nusbaum *et al*., 2017) and increases the number of possible targets, given that neuropeptides can be released at synapses and non-synaptic sites, diffuse from its source, and act far from the release site (van den Pol, 2012). We have not pursued direct postsynaptic targets of the Mip2M descending neurons in the present work. Instead, we concentrated our efforts on the functional characterization of neurons modulated by Mip via SPR and found that the circuit underlying GSB is complex and is composed of at least cholinergic, GABAergic, and some neurons within the *D42>* population. Three main conclusions about the circuit can be drawn from our data. First, the circuit has no apparent hierarchical organization. Silencing of SPR in individual populations did not fully recapitulate the effect of the lack of *Mip*, and even had opposite effects depending on the population targeted. Second, no particular element of the behavioral sequence was associated with a specific neuronal subtype. Third, motor neurons are not the primary target of Mip. We propose that Mip modulates the activity of interneurons in the intermediate neuropil to reconfigure motor circuits properties and produce a stage specific motor output pattern. While the Mip descending neuron is consistent with the logic of one neuron = one behavior, the circuit downstream of it does not seem to operate under the same logic, at least as action components are concerned, nor does it seem to directly control the initiation of central pattern generators that produce each of the motor patterns. Rather, Mip/SPR appear to act on an overlying layer that dictates how those neuronal circuits interact between themselves, orchestrating the transitions between action components and the order of events of the behavioral sequence. This in-depth characterization of GSB behavioral sequence will allow us to identify circuit components in detail.

Interestingly, the inhibition of GSB execution by means of the hyperpolarization of Mip2M descending neurons did not block PMP progression; it still progressed to post-GSB but only after a short period of inactivity similar to GSB duration. This suggests that, different from other innate behaviors in *Drosophila*, such as egg laying, which is subjected to sensory feedback that guides progression to subsequent component actions when the task has been accomplished (Cury and Axel, 2023), the PMP is executed without (or with negligible) feedback control, at least at the GSB to post-GSB transition. The spreading behavior is not gated by the effective expulsion of glue, nor the post-GSB by the spreading of glue over the ventral surface of the body, confirming previous results (Heredia *et al*., 2021). Once the PMP is released by epidermis-derived Dilp8, it proceeds until completion. Behavioral progression in ecdysis results from the orderly secretion of an array of neuropeptides, capable of promoting the phase specific motor patterns and suppressing the previous one (Truman, 2005). It will be relevant to determine if other neuropeptides besides Mip similarly act downstream or in parallel to Dilp8 to orchestrate the PMP behavioral sequence, as well as the extent of their modulatory effect. A thorough analysis showed that the effect of Mip is not limited to the GSB stage. mhc>>GCaMP monitoring confirmed that the three main stages of the PMP (pre-GSB, GSB, and post-GSB) were present with the typical sequential organization in *Mip^1^* animals. However, the three stages were on average ∼60, 30, and 10% shorter, respectively, than those in control siblings. Further differences in motor-activity patterns were unveiled by the characterization of pre-GSB contractions, which were fewer, shorter, and more frequent (Supplementary Figure 6). This data suggests that the Mip/SPR pathway is likely to modulate multiple aspects of the PMP in addition to GSB and indicates a fertile field of study.

Insect Mips belong to the superfamily of Wamide neuropeptides, characterized by a conserved amidated tryptophan residue at the C-terminal end. The ancient origin of Wamide signaling is supported by its presence in the last common ancestor to cnidarians and bilaterians (Jékely, 2013). Nevertheless, clear members of the Wamide family have not been identified in some protostome phyla like hymenopterans and in deuterostomes genomes (Mirabeau *et al*., 2013; Hauser *et al*., 2010), suggesting that they have been lost multiple times during evolution. Vertebrates encode at least two putative orthologs of the *Drosophila Mip receptor,* SPR, represented by human orphan receptors GPR139 and GPR142 (Mirabeau *et al.,* 2013). Interestingly, similar to SPR, GPR139 is also expressed in the nervous system and mice lacking GPR139 activity show a series of neuropsychiatric behavior abnormalities (Dao *et al*., 2021), indicating an ancient and conserved role for SPR-family receptors in behavioral control. Even though the physiological endogenous ligands of vertebrate SPR orthologs remain to be firmly established, they have been shown to respond to essential aromatic amino acids (like L-Tryptophan and L-Phenylalanine), to peptides derived from melanocyte stimulating hormone (MSH), and to the core motif HFRW (Isberg *et al.,* 2014; Liu *et al*., 2015; Nohr *et al*., 2017).

Some strikingly similar functions have been described for distant members of the Wamide superfamily. Life cycle of many marine invertebrate species, like corals and hydroids (Cnidarians) and annelids (protostomes) consist of mobile and sessile or pelagic and benthic forms respectively. Transition from one to the other is mediated by environmental cues and can be experimentally stimulated by synthetic Wamides that induce a change in swimming behavior (Leitz *et al*., 1994; Erwin *et al.,* 2010; Lechable *et al.,* 2020). The Mip orthologue from the annelid *Platynereis* is expressed in chemosensory and neurosecretory neurons in the apical organ, a specialized sensory cluster that regulates larval settlement. Similar as to how the fly Mip plays a critical role in defining–via glue expulsion and spreading behavior–the final location of the larval life history stage (the puparium attachment site), *Platynereis* Mip also orchestrates larval settlement behavior. Treatment of *Platynereis* larvae with synthetic neuropeptide triggers larval sinking and exploratory crawling behavior, an effect that is mediated by an *SPR* ortholog (Conzelmann *et al.,* 2013). Together, these findings suggest that one of the ancestral functions of Mip signaling may have been the modulation of neuronal circuits that shape behaviors at developmental transitions.

## MATERIALS AND METHODS

### *Drosophila* husbandry and stocks

*sfGFP::Lgr3^ag^*^5^ and *mhc-LexA* (*mhc-LHV2*) were previously described (Garelli *et al*., 2015; Heredia *et al*, 2021). *Mip^1^* (Min *et al*., 2016) and its background control, *DfExel6234* and *Δ^130^* (Liu and Kubli, 2003) were gifts from Carlos Ribeiro, *UAS-Kir2.1* on III (*w*; P{w[+mC]=UAS-Hsap\KCNJ2.EGFP}7*) was a gift from Nara Muraro, *UAS-SPR-IR* (*w^1118^; P{w[+mC]=GD3236}v7061* from VDRC), *VGlut-GAL4* (*w^1118^; P{w+mW.hs]=GawB}VGlut[OK371]*), *ChAT-GAL4* (*w*; Mi{Trojan-GAL4.0}ChAT[MI04508-TG4.0] CG7715[MI04508-TG4.0-X]/TM6B, Tb[1]*) (Diao *et al*., 2015), *Gad-GAL4* (*P{Gad1-GAL4.3.098}*) (Ng *et al*., 2002), and *tsh-GAL4* (*P{w[+mW.hs]=GawB}tsh[md621]*) (Calleja *et al*., 1996) were gifts from Fernanda Ceriani, *D42-GAL4* (*P{GawB}D42*) was a gift from Rafael Pagani, *Sgs3::GFP* (Biyasheva *et al*., 2001) was a gift from Mariana Melani.

The following lines were obtained from the Bloomington Drosophila Stock Center at Indiana University:

*BL41680 y^1^ sc* v^1^ sev^21^; P{y^+t7.7^ v^+t1.8^=TRiP.HMS02244}attP2 (Mip-IR-V20)*

*BL26246 y^1^ v^1^; P{y^+t7.7^ v^+t1.8^ =TRiP.JF02145}attP2 (Mip-IR-V10)*

*BL77389 y^1^ sc* v^1^ sev^2^; P{y^+t7.7^ v^+t1.8^ =TRiP.HMC06520}attP40 (UAS-SPR-IR(2))*

*BL51984 w^1118^; P{w^+mC^=Mip-GAL4.TH}2M (Mip-GAL4.TH)*

*BL28838 w*; P{w^+mC^=UAS-TeTxLC.tnt}G2 (Tetanus toxin)*

*BL28839 w*; P{w^+mC^=UAS-TeTxLC.(-)Q}A2 (Mutated tetanus toxin)*

*BL6596 w*; P{w^+mC^=UAS-Hsap\KCNJ2.EGFP}1 (UAS-Kir2.1)*

*BL50395 w^1118^; P{y^+t7.7^ w^+mC^=GMR48H10-GAL4}attP2*

*BL39171 w^1118^; P{y^+t7.7^ w^+mC^=GMR57C10-GAL4}attP2*

*BL84692 w[*] TI{2A-GAL4}SPR[2A-GAL4]*

*BL24646 P{w^+mC^=UAS-Dcr-2.D}1, w[1118]*

*BL33064 w^1118^; P{w^+mC^=UAS-DenMark}2, P{w^+mC^=UAS-syt.eGFP}2; In(3L)D, mirr^SaiD1^ D^1^/TM6C, Sb^1^*

*BL9313 w*; P{w^+mC^=Tdc2-GAL4.C}2*

*BL44277 w^1118^; P{y^+t7.7^ w^+mC^=13XLexAop2-IVS-GCaMP6f-p10}su(Hw)attP5*

*BL32218 w*; P{y^+t7.7^ w^+mC^=10XUAS-IVS-mCD8::RFP}attP2*

*BL64089 w^1118^ P{GMR57C10-FLPG5.PEST}su(Hw)attP8; PBac{10xUAS(FRT.stop)myr::smGdP-*

*HA}VK00005 P{10xUAS(FRT.stop)myr::smGdP-V5-THS-10xUAS(FRT.stop)myr::smGdP-FLAG}su(Hw)att_P1 (MCFO)*

*BL36887 y^1^ sc* v^1^ sev^21^; P{y^+t7.7^ v^+t1.8^=TRiP.GL01056}attP2/TM3, Sb^1^*

*BL32194 w*; P{y^+t7.7^ w^+mC^=20XUAS-IVS-mCD8::GFP}attP2*

*BL36303 y^1^ v^1^; P{y^+t7.7^=CaryP}attP2*

*w^1118^* or *y^1^ v^1^; P{y^+t7.7^=CaryP}attP2* were used as control lines.

The stocks used in the screen are listed in Supplementary Table 1.

All other stocks were generated in this study as described below. Stocks are maintained at low densities at 18 °C in a 12-h light/dark cycle. All experiments were done at 25°C in a 12-h light/dark cycle.

### Immunofluorescence analyses

CNS of larvae L3 wandering were dissected in Schneider Medium (Biowest – cat. #L0207 or Gibco - cat. #21720-024), fixed for 30 min in 4% paraformaldehyde, rinsed with PBS with Triton (0.3%) (PBST), incubated with primary antibody for 24-48h and with fluorescently labeled secondary antibody for 2–24 h in PBST with 1% bovine serum albumin. Samples were washed 3× for 30 min each in PBST after each antibody incubation. Nuclei were counterstained with DAPI (Sigma) and tissues were mounted in Fluoromount-G (Southern Biotech). Primary antibodies used were: anti-Mip (*Periplaneta americana* MIP antiserum (Predel *et al*., 2001)); Rabbit Anti-GFP, 1:200 (Life Technologies, A11122); Mouse Anti-Fasciclin II-s, 1:10 (DSHB, 1DA); Mouse Living Colors DsRed, 1:500 (Clontech, 632496); Mouse Anti-GFP-4C9, 1:200 (DSHB, DSHB-GFP-4C9); Rat DN-Ex #8-s, 1:50 (DSHB, AB_528121, a-Cadherin), Mouse anti-V5 tag purified, 1:200 (eBioscience, 14-6796-82); Rabbit anti-HA tag, 1:800 (Cell Signalling Technology, 3724). Secondary antibodies were: Alexa Fluor 488, anti-mouse (Life Technologies, A11029); Alexa Fluor 488, anti-rabbit (Life Technologies, A11070); Alexa Fluor 568, anti-rabbit (Life Technologies, A11036); Alexa Fluor 594, anti-mouse (Life Technologies, A11020); Alexa Fluor 594, anti-rat (Life Technologies, A11007). Antibodies against the V5 and HA tags were used for MCFO.

### Generation of transgenic lines

#### UAS-Mip

Mip CDS (FBtr0075241) was amplified from genomic DNA (gDNA) from w[1118] *Drosophila melanogaster* using primers #1 and #2, containing BglII and NotI restrictions sites, generating a 656 bp PCR fragment. Mip 5’UTR-CDS-3’UTR construct was amplified by SOEing PCR using primers #3 to #6, generating a 1.3 Kb PCR fragment.

#1_Mip_Cloning_BglII F TGCTTAGCGGCCGCTTAGTTGCTGGGCAACTGGG

#2_Mip_Cloning_NotI R TAAGCAAGATCTACAGCTATGGCTCACACTAA

#3_ Cloning 5’Mip3’BglII F TAAGCAAGATCTGCGTATCATTAGCAGTTGAT

#4_ Cloning 5’Mip3’NotI R TGCTTAGCGGCCGCACTTTCAACGTATTTGTATT

#5_ Mipcds-lnk5utr F ATATAGTCAGCTATGGCTCACACTAAGA

#6_ Mipcds-lnk5utr R TGTGAGCCATAGCTGACTATATATCTCCT

All PCR fragments were checked by Sanger Sequencing, digested with BglII and NotI restriction enzymes and then ligated to pJFRC-MUH (Addgene #26213). The plasmids generated (pJFRC-MUH-mipCDS, and pJFRC-MUH-5’UTR_mipCDS_3’UTR) were then injected in y[1] w[1118]; Pbac{y[+]-attP-9A}VK00018 line (BL9736) using PhiC31 transgenesis program.

#### *Mip2M-GAL4* and *Mip3M-GAL4*

The putative *Mip* promoter region was amplified from genomic DNA by PCR using the primer pair GGAGGAATTCagcagcaaaaagtcggaaaa and GGAGAATTCGGATCCgtgaatttacgggcacgagt. This amplified 2.4 kb upstream of the Mip precursor coding sequence, including the *Mip* transcription initiation site. The resulting PCR product was cloned into pCR-TOPO (Thermofisher) and subsequently removed by digestion with *MunI* and *BamHI*, gel purified and exchanged for the *Akh* promotor in the *pAkh-GAL4* vector (Isabel *et al.,* 2005) after its digestion with *EcoRI* and *BamHI*. Transgenic flies were generated by BestGene (Chino Hills, CA, USA) using P-element transformation in *w^1118^* embryos. Lines were established from several individual transformants that carried the insert on different chromosomal localization. The lines *Mip2M-Gal4* (*w^1118^;;Mip-GAL4*) and *Mip3M-GAL4* (*w^1118^;Mip-GAL4*) were used in this study.

### Puparium aspect ratio measurement

Aspect ratio was measured as previously described (Heredia *et al*., 2021). Briefly, pictures of puparia taken under a dissecting scope were analyzed using ImageJ and/or Amscope software. Length was measured from the anteriormost edge of the pupa to the most anterior anal papilla. Width was measured in the widest part of the middle third of the pupa.

### Evaluation of GSB performance, quantification of muscle GCaMP fluctuations and behavior annotation

Wandering larvae filmed in the pupariation monitor device (Heredia *et al*., 2021) under white or blue light were evaluated for the occurrence of GSB. This behavior follows a stereotypical series of movements (Heredia *et al*., 2021). It starts with a prolonged contraction of the anterior segments, followed by back-and-forth peristaltic waves that initially involve the most anterior segments and move more posteriorly as the behavior progresses. One or two whole body contractions accompanied by a lateral “head waving” movement mark the end of the behavior. The performance of GSB was classified as normal when all these elements were present, and abnormal otherwise. See supplementary video 1 for reference.

Larvae were individually placed in 2×3 wells 3D printed arenas (Heredia *et al*., 2021) in the monitoring device for GCaMP fluorescent signal detection. Muscle GCaMP fluctuations from *mhc>>GCaMP* larvae were evaluated on behaving animals using custom written python scripts available in https://github.com/AndresGarelli/Larva_Tracking_OpenCV. Videos were recorded at 10 fps and GCaMP intensity measured once per second. The output was analyzed offline to calculate the following parameters:

- **Duration of pre-GSB contractions**: time in seconds during which the fluorescence intensity of GCaMP is above 50% of the difference between the baseline (F0) and maximum (Fmax) values of each peak, where F0 is the minimum value of the preceding 10 seconds.
- **Period**: is the time between the start of two consecutive peaks of GCaMP.
- **Duration of pre-GSB, GSB, and post-GSB**

For the evaluation of the sequence of behaviors during GSB at high resolution, up to twelve wandering larvae were individually tracked in two 1×6 3D-printed arenas using a python script that displays GCaMP fluctuations live. When pre-GSB contractions were detected, the arena was moved to a fluorescent stereomicroscope and behavior recorded at 10 fps using an Amscope MU300 camera. Note: on most occasions, larvae continued the behavior and smoothly transitioned to GSB upon moving the setup to the stereomicroscope. However, in a few opportunities, pre-GSB was interrupted and the larvae displayed a train of peristaltic waves or escape response, probably to high intensity illumination. This suggests that there seems to be a no-return point in the behavior, and also that prior to that point, the fixed action pattern can be interrupted.

Behavior during GSB was manually annotated offline using Anvil (http://www.anvil-software.de/) (Kipp, 2001) and graphically represented as a GSB behavioral sequence using custom python scripts. Muscle GCaMP traces were used to aid in the identification of the beginning of GSB in those conditions that lacked a tetanic contraction.

For the calculation of the “opposing transition index”, peristaltic waves were considered to be consecutive if they happened immediately one after the other or separated by a pause without contractions. Transitions between consecutive peristaltic waves were given a value of 1 if they moved in opposite directions and 0 if they traveled in the same direction.

The index was calculated by dividing the sum of transition values by the total number of transitions between consecutive peristaltic waves. An index of 1 is the result of a perfect back and forth alternation and, on the contrary, an index of 0 means that all consecutive waves move in the same direction.

### Glue expulsion

GFP fluorescence is preserved over the surface of the puparium after the salivary gland fluid has dried. To evaluate the release of glue, wandering larvae expressing *Sgs3::GFP* were transferred to empty vials and glue expulsion was evaluated the next day. The complete absence of green fluorescence was considered glue retention, while the observation of varying amounts of green signal over the puparium were scored as glue release.

### Optogenetic stimulation

Optogenetic stimulation experiments were performed in modified pupariation arenas (Heredia *et al*., 2021) consisting of six wells arranged in a 3 x 2 array individually back illuminated with green 5 mm high brightness through-hole LED controlled by the Raspberry Pi (Supplementary Figure 7). ReaChR response to light peaks at 590 nm, but it can also be effectively activated by blue and green wavelengths (Inagaki *et al*., 2014; Lin *et al*., 2013).

Flies were allowed to lay eggs on regular food and between 2-3 d later, larvae were transferred to food supplemented with 100 µM all-trans retinal and kept in the dark at 25 °C. When larvae reached wandering stage, they were individually placed in the wells of the optogenetics arena and the whole set up was placed inside an incubator at 25 °C illuminated with 940 nm IR LEDs from the front. Larvae were recorded with a PiNoIR camera for up to 24 h and subjected during the whole recording to ten light pulses of 0.5 s of duration at 1 Hz (0.5 s ON, 0.5 s OFF x 10 times) every 5 min. The intense light coming from the LEDs was filtered by placing two sheets of exposed photographic film in front of the camera lens. Larval behavior analysis was performed off-line. Time scale was synchronized at pupariation (*i.e.*, determined by anterior spiracle eversion, execution of GSB and initiation of post-GSB stage) for each larva and the behavior in response to light stimuli evaluated during 60-min periods starting either at 3 h or 1 h before pupariation (“3-2 h before post-GSB” and “1-0 h before post-GSB”). Response to light pulses was considered a positive, GSB-like behavior, when at least two consecutive back and forth peristaltic waves were observed.

### Statistical analysis

For all tests, alpha was set at 0.05 a priori. For quantitative data, comparison between multiple conditions was done using ANOVA when samples had normal distribution (Shapiro-Wilk test and equal variance). If these conditions were not met, Kruskal-Wallis One Way Analysis of Variance on Ranks was performed. If the result of these tests was statistically significant, then Tukey or Dunn’s post-hoc tests were applied for the assessment of statistical significance of pairwise comparisons.

Comparison between two conditions were done using one-tailed unpaired (unless noted otherwise) Student’s t-test, when samples had normal distribution and equal variance. Alternatively, a Mann-Whitney Rank Sum Test or Wilcoxon signed rank test (for paired samples) was performed. Statistically-significantly different comparisons were denoted with an asterisk.

For frequency (binary) data, like GSB execution and glue release, two-tailed Fisher’s exact test was used when comparing two conditions. Comparisons between more than two conditions when significant (at alpha 0.05), were followed by post-hoc pairwise two-tailed Fisher’s exact tests applying the conservative Bonferroni correction for multiple tests (adjusted alpha = alpha/*N* comparisons) (McDonald, 2014). Tests were performed using the online calculators available at https://astatsa.com/FisherTest/ and https://www.langsrud.com/fisher.htm.

Results from multiple comparisons were presented, except where explicitly denoted otherwise, with a letter scheme where conditions or genotypes sharing the same letter were not statistically significantly different. Statistical analyses were performed either using Graphpad Prism or the indicated online calculators.

## AUTHORS CONTRIBUTIONS

M.F-A., R.Z., F.H., Y.V., J.M., K.P., J.I., M.A., M.S.P., and A.G. performed genetic, phenotypic, molecular, and behavioral experiments. M.F-A., J.M., J.A.V., and C.W. developed critical reagents. F.H., A.M.G., and A.G. supervised the work. A.M.G. and A.G. contributed to image and data analyses and wrote the manuscript with the help of all authors.

## ACKNOWLEDGMENTS

We thank Drs. Carlos Ribeiro, Nara Muraro, Fernanda Ceriani, Rafael Pagani, and Mariana Melani for fly stocks and reagents. We thank lab members for discussions and/or comments on the manuscript. We thank Carolina Gomila for technical assistance. Stocks obtained from the Bloomington Drosophila Stock Center (NIH P40OD018537) and Vienna Drosophila Resource Center (VDRC; http://stockcenter.vdrc.at) were used in this study. Work in the Integrative Biomedicine Laboratory was supported by the FCT (PTDC/BIA-BID/31071/2017; PTDC/MED-NEU/30753/2017 (LISBOA-01-0145-FEDER-030753, with financing from the Programa Operacional Lisboa 2020); EXPL/BIA-BID/1524/2021; EXPL/BIA-COM/1296/2021; 10.54499/2022.03859.PTDC), by the UIDs iNOVA4Health (10.54499/UIDB/04462/2020) and cE3c (10.54499/UIDB/00329/2020), by LS4FUTURE (10.54499/LA/P/0087/2020) and CHANGE (10.54499/LA/P/0121/2020) financed by the FCT/MCTES (Portugal), and by Congento LISBOA-01-0145-FEDER-022170, co-financed by FCT/Lisboa2020; UID/Multi/04462/2019. Work in the Garelli lab was supported by ANPCyT (Agencia Nacional para la Promoción de la Ciencia y la Tecnología, PICT 2017-0254 and PICT 2020-01568), CONICET (PIP11220150100182CO) and Universidad Nacional del Sur PGI 24/B288. The generation of Mip-GAL4 lines was supported by the Deutsche Forschungsgemeinschaft (DFG WE 2652/4-1,2, to CW). AMG and FH were individually supported by grants 10.54499/CEECINST/00102/2018/CP1567/CT0031 and 10.54499/DL57/2016/CP1457/CT0016, respectively. AG and YV are CONICET researchers. MA held a CONICET doctoral fellowship and JI an undergraduate fellowship from Consejo Interuniversitario Nacional.

## Supplementary Material

**Supplementary Table 1.**
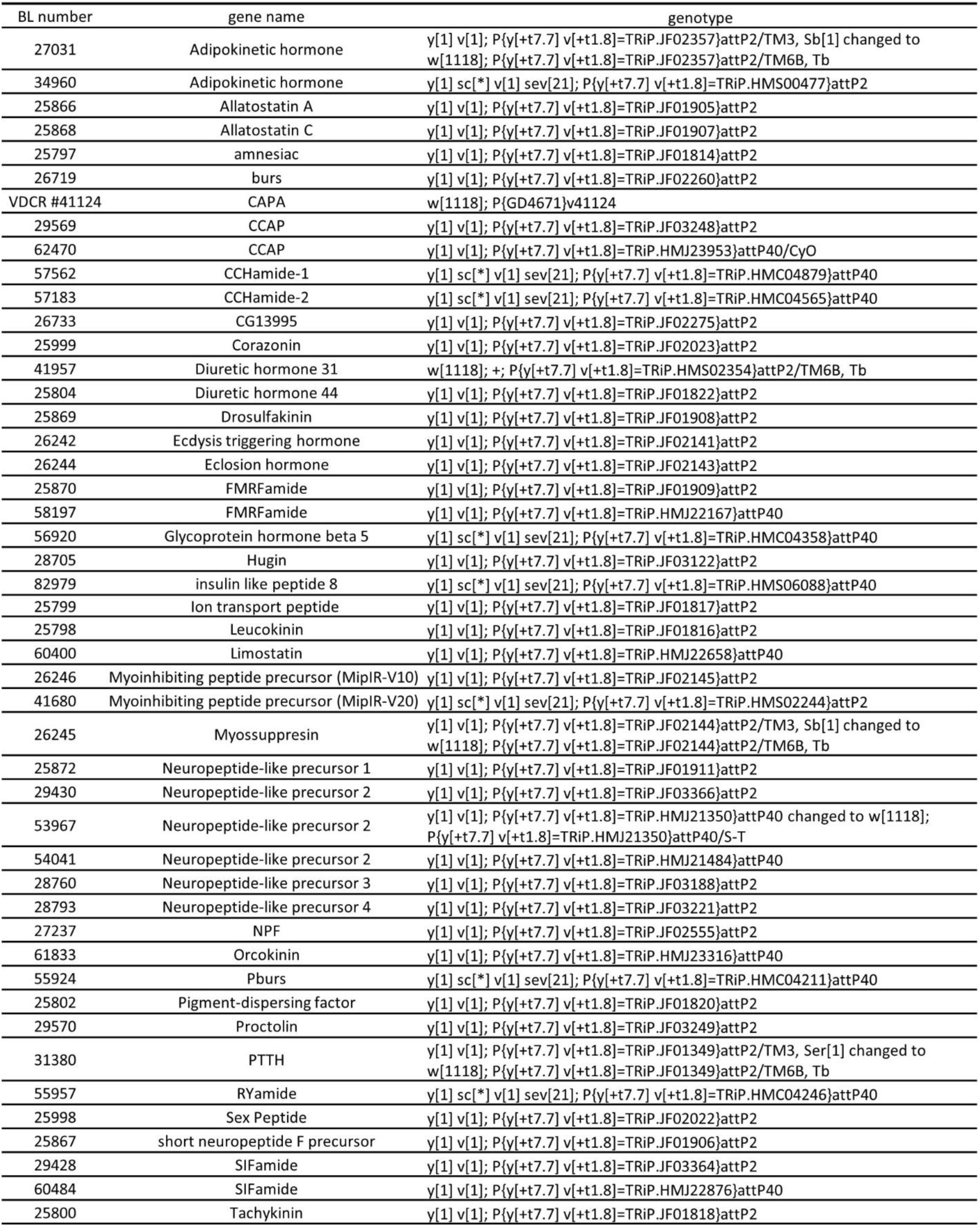
List of stocks used in the neuropeptide targeted screen, indicating the stock number of the Bloomington Stock Collection (or Vienna Drosophila Research Centre), gene targeted, and genotype.

**Supplementary Figure 1.**
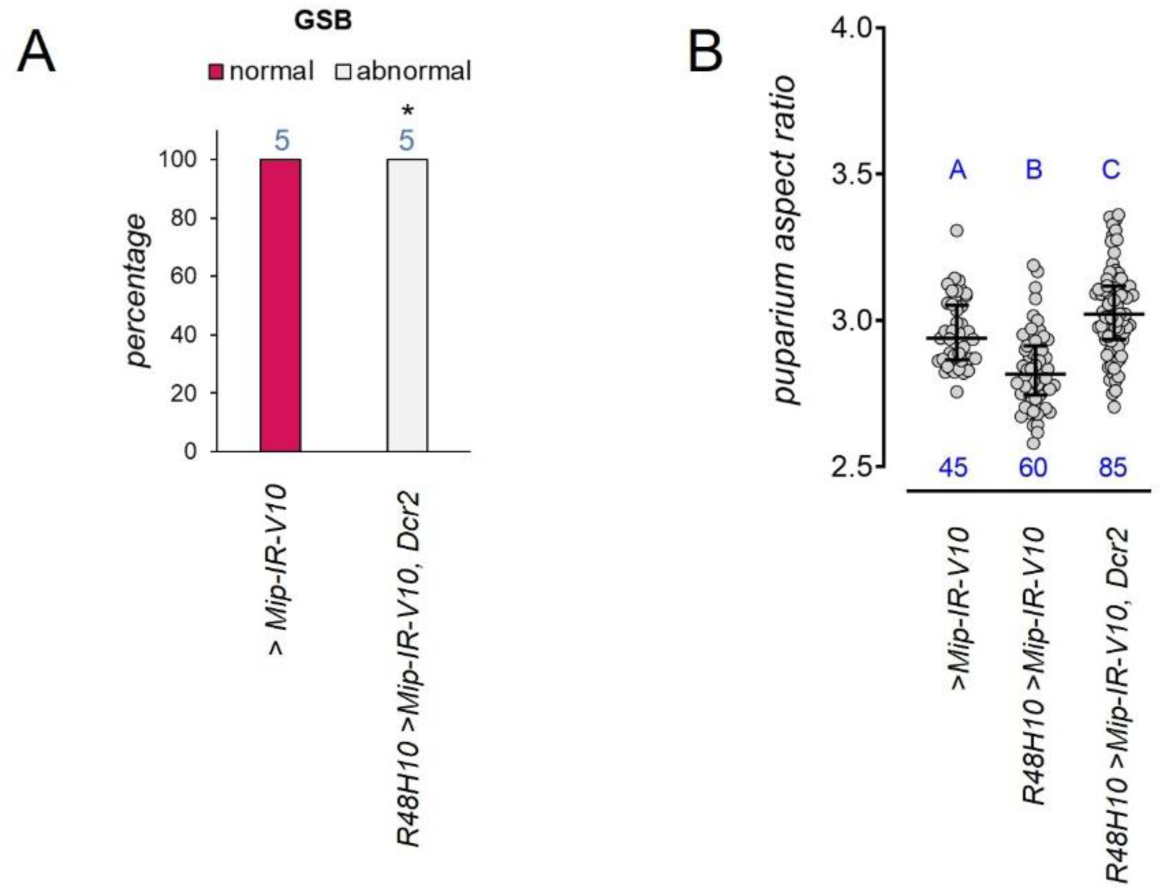
A second RNAi line against *Mip* alters GSB execution. A) Percentage of animals that perform a normal GSB. *P* = 0.0079, Fisher’s exact test. Blue number, N. B) Dot plot showing puparium aspect ratio of indicated genotypes. Dots: one animal. Horizontal bars, median, and 25-75 percentile. Blue numbers, N. Same blue letter, *P* > 0.05, Kruskal-Wallis and Dunn’s test.

**Supplementary Figure 2.**
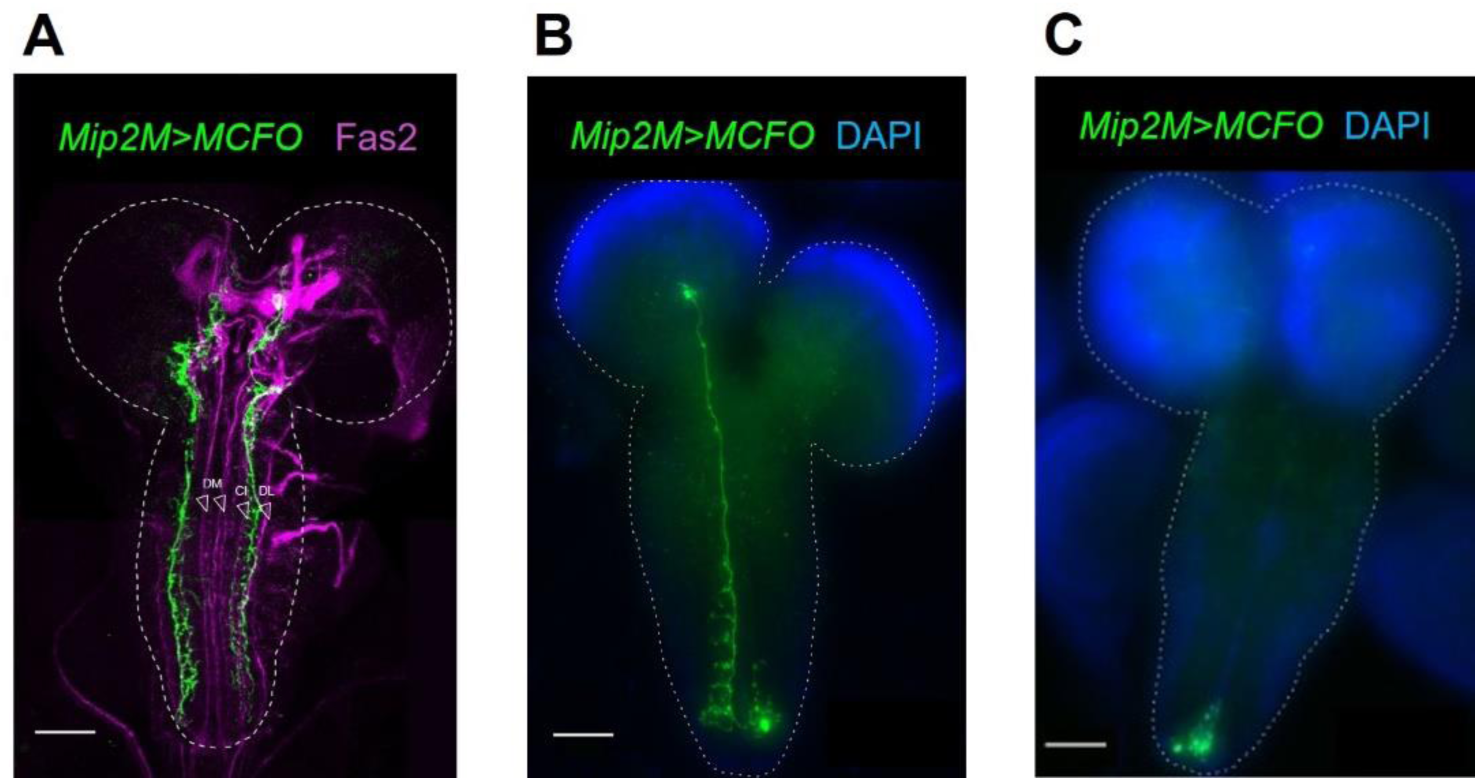
Morphology of Mip2M neurons using Multicolor Flip Out (MCFO) A) MCFO clones (green) labelling the two descending Mip neurons. Dorso medial (DM), Central Intermediate (CI) and Dorsolateral (DL) Fas2 (anti-Fas2, magenta) tracts serve as a reference. Dotted line, CNS contour. Scale bar, 50 µm. B) MCFO clone of the ascending interneuron with the cell body in abdominal segment A8/9. It has a small dendritic arborization and a single axon that crosses to the contralateral side and changes direction close to the midline, crossing the whole ventral nerve cord and reaching the protocerebrum. Along the first half of its trajectory, the single axon emits small ramifications arranged in a segmental fashion. Dotted line, CNS contour. Scale bar, 50 µm. C) MCFO clone of a local interneuron with the cell body in abdominal segment A8/9 and short ipsilateral projections to the contiguous neuromere. Dotted line, CNS contour. Scale bar, 50 µm.

**Supplementary Figure 3.**
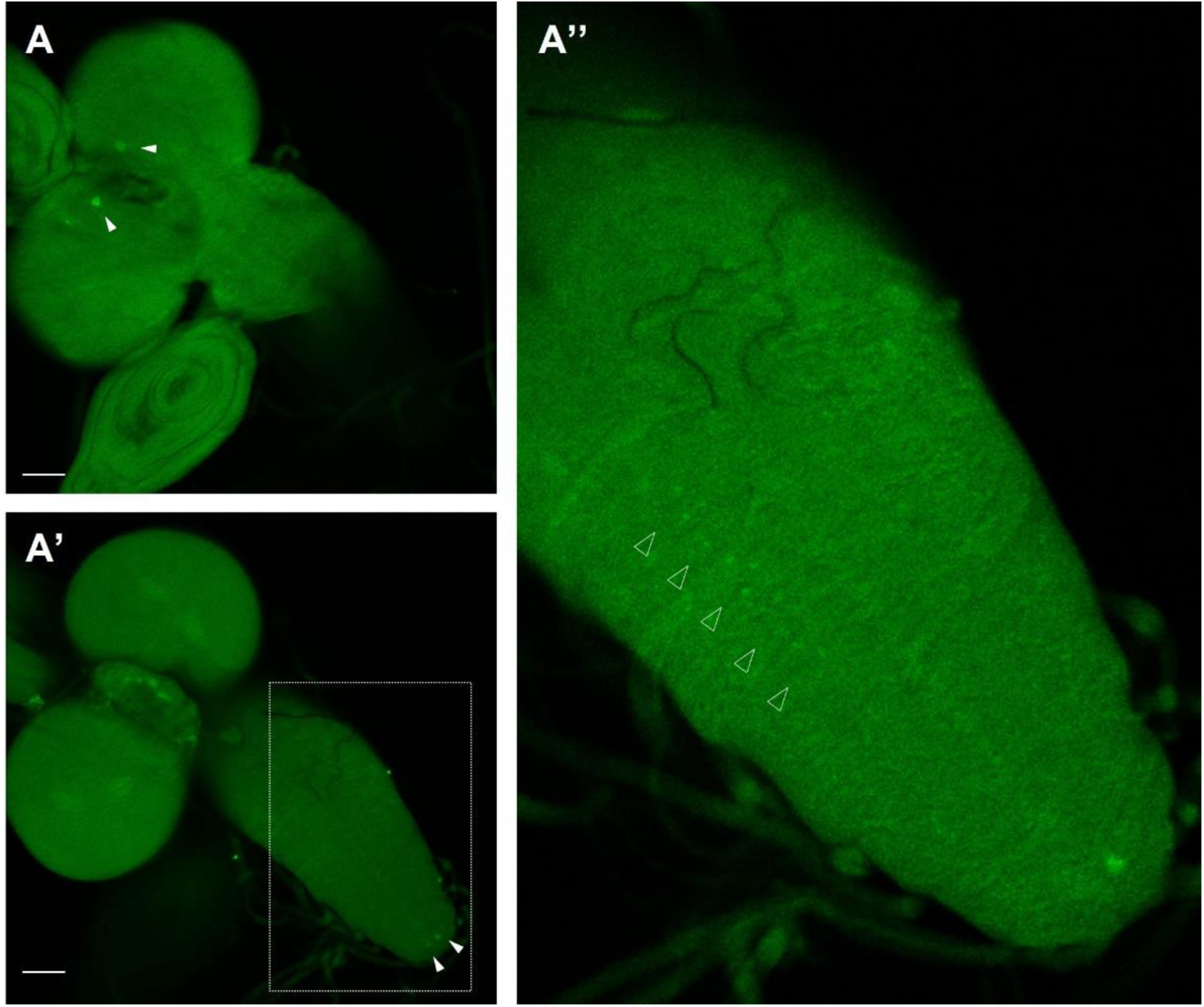
A) Confocal sections at different Z positions of another Mip1, Mip2M>Mip L3 larval depicting Mip immunoreactivity in the cell bodies of the Mip2M descending neurons (A, arrowheads), tip neurons (A’, arrowheads) and a detail of VNC showing a punctate pattern compatible with Mip detection in the axons of descending nuerons. Scale bar, 50 µm.

**Supplementary Figure 4.**
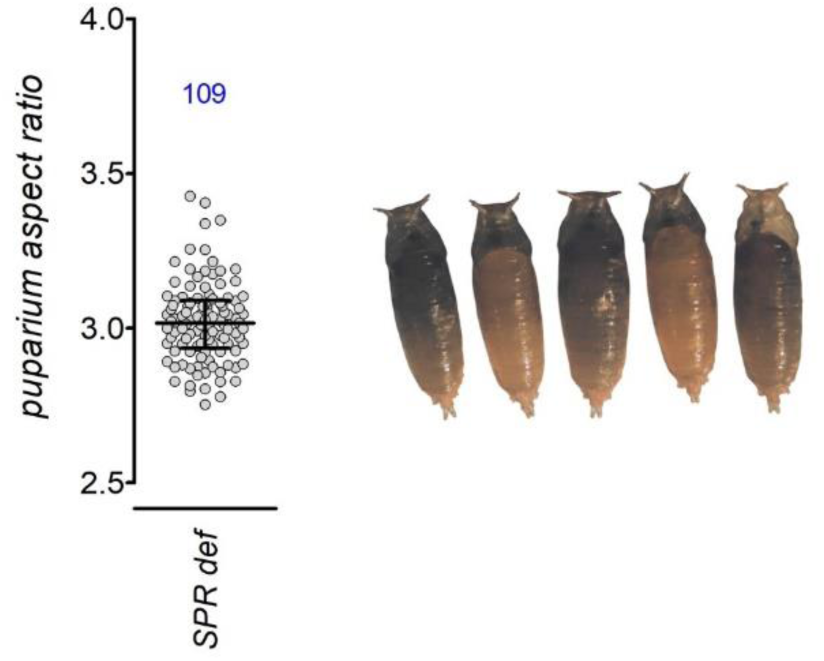
*SPR* mutants have slightly increased puparium aspect ratio but do not show major morphological alterations. Dot plot showing puparium aspect ratio of indicated genotypes (*SPR def* (full genotype *Df(1)Exel6234/Df(1)Exel6234*)) Representative images are shown on the right. Dots: one animal. Horizontal bars, median, and 25-75 percentiles. Blue numbers, N.

**Supplementary Figure 5.**
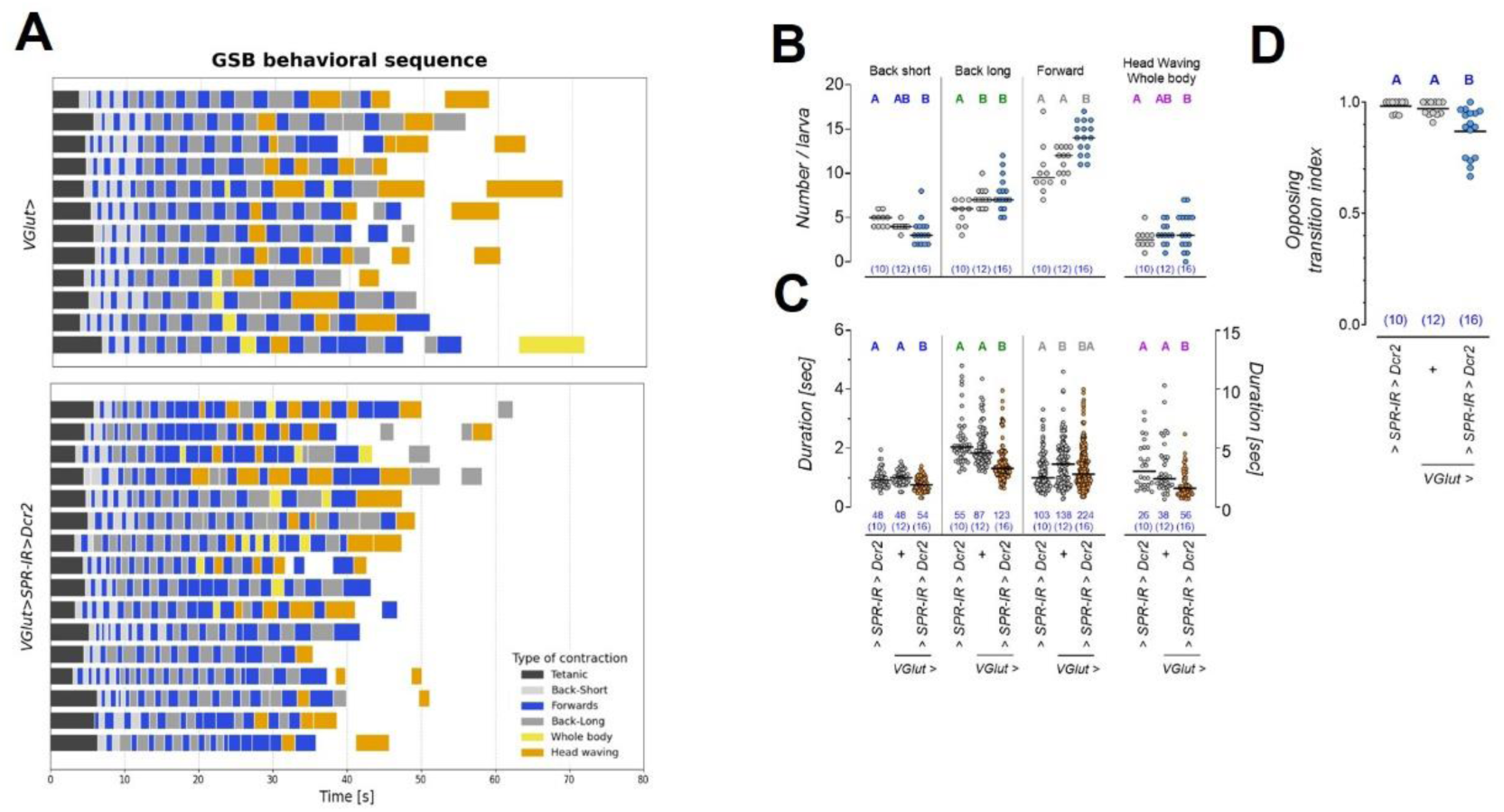
SPR is required in glutamatergic neurons to control aspects of GSB. A) Ethograms displaying the sequence and duration of GSB behavioral pattern. Knockdown of *SPR* in glutamatergic neurons, which comprise motor neurons, does not affect the back-and-forth alternate pattern of the initial short contractions. However, self-transition events between backwards long or forwards waves are frequent. B) The number of forward waves is increased in *VGlut>SPR-IR>Dcr2*. Dot plots showing the quantification of the occurrence of each type of movement. Dots represent one larva. Horizontal bar, mean value. Blue numbers, N. Same letters, *P* > 0.05, Kruskal Wallis and Dunn’s test. C) Knockdown of *SPR* accelerates backward peristaltic waves, which is reflected in their shorter duration. Whole body and head waving movements, represented on the second Y-axis, are also shorter. Dots represent one contraction. Horizontal bar, median value. Blue numbers, N (number of larvae). Same letters, *P* > 0.05, Kruskal Wallis and Dunn’s test. D) The increased frequency of self-transitions between peristaltic waves is reflected in a reduced opposing transition index. Dots represent one larva. Horizontal bar, mean value. Blue numbers, N. Same letters, *P* > 0.05, Kruskal Wallis and Dunn’s test.

**Supplementary Figure 6.**
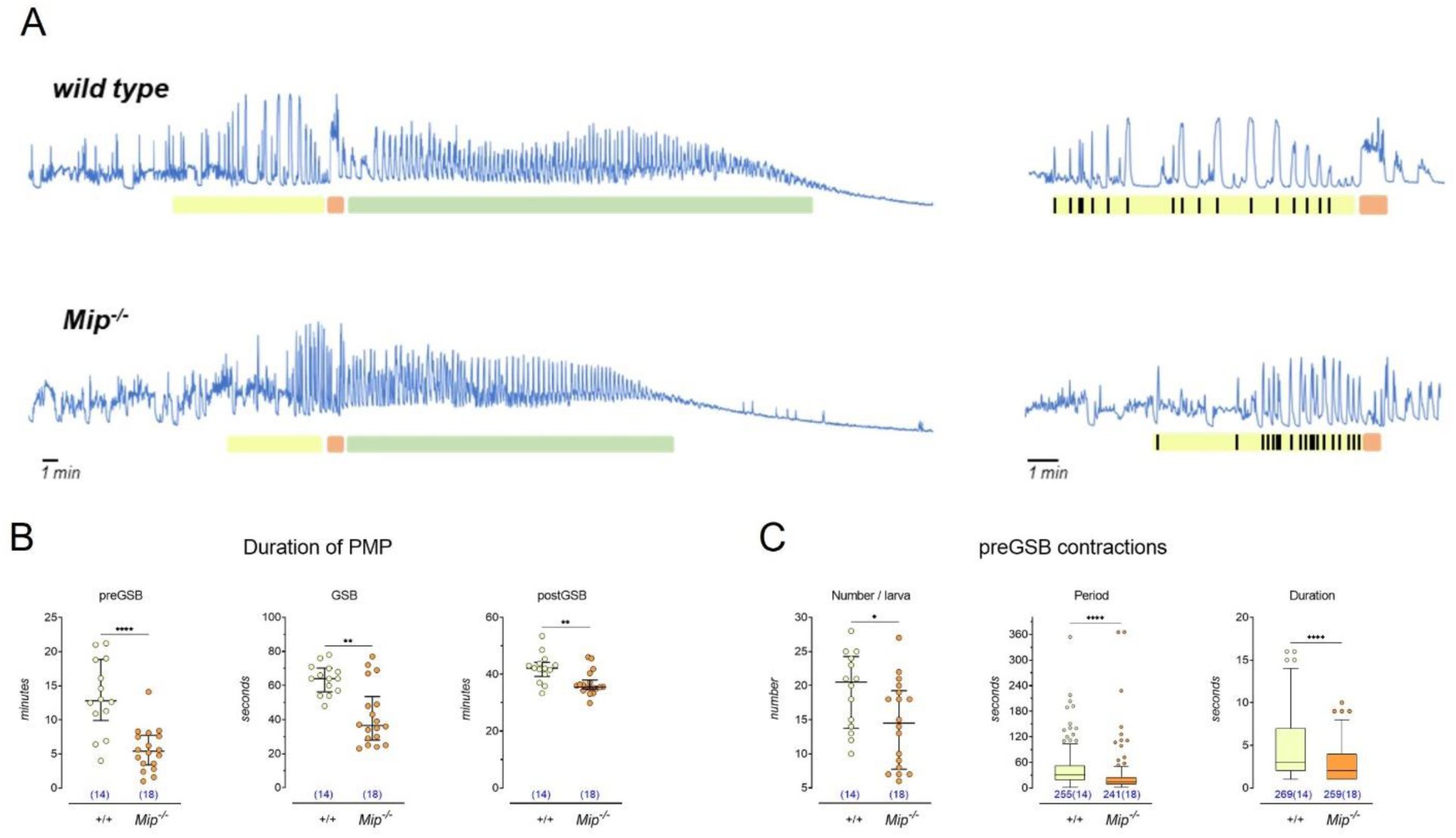
The Mip/SPR pathway modulates the other PMP stages. Muscle-GCaMP monitoring of *Mip^1^* animals showed that the three main stages of the PMP (pre-GSB, GSB, and post-GSB) were present with the typical sequential organization. This indicated that Mip activity is not required for the initiation of the behavioral subprograms nor the transitions between them. Shown are representative traces of *Mip^1^* mutant animals and the corresponding wild type background control (A). However, the three PMP stages were on average ∼60, 30, and 10% shorter, respectively, than those in control siblings (P < 0.0001, t-test; P = 0.0016 and P = 0.0031, Mann Whitney test respectively) (B). Further differences in motor activity patterns were unveiled by the characterization of pre-GSB contractions. The sustained whole-body contractions that lead to the retraction of the anterior segments and mouth hooks are typically separated by periods of inactivity that may occasionally be interrupted by short contractions. The pattern of the pre-GSB stage is abnormal in *Mip^1^* mutants. Example representative traces of GCaMP fluctuations during preGSB are shown in A-right. Namely, there is a decrease in the number (P = 0.0321 unpaired t-test) and the period of contractions, that is, the time between two consecutive contractions (P < 0.0001 Mann Whitney test). The duration of contractions also decreased (P < 0.0001 Mann Whitney test). The sustained whole-body contractions were less frequent. Only 5% of the total number of pre-GSB contractions measured was longer than 6 seconds, and the longest contraction ever observed for *Mip^1^* mutants lasted 10 seconds. N= 14 and 18 for background control and *Mip^1^* mutant larvae. Dot plot: one dot represents one larva. Lines are median and interquartile range. Box plots: one dot represents one contraction. The number of contractions is indicated between parentheses. Whiskers are plotted following the Tukey method (interquartile ranges x 1.5). Still, the anomalous activity displayed during the execution of pre-GSB was effective in remodeling larval shape and producing a visually normal puparium, apart from the slightly increased aspect ratio (Figure 5A). As a whole, mutant animals failed to organize a regular pattern of pre-GSB-related muscle activity, with an increased frequency of short contractions and the absence of the long-contractions/relaxation series. This data suggests that the Mip/SPR pathway is likely to modulate multiple aspects of the PMP beyond GSB

**Supplementary Figure 7.**
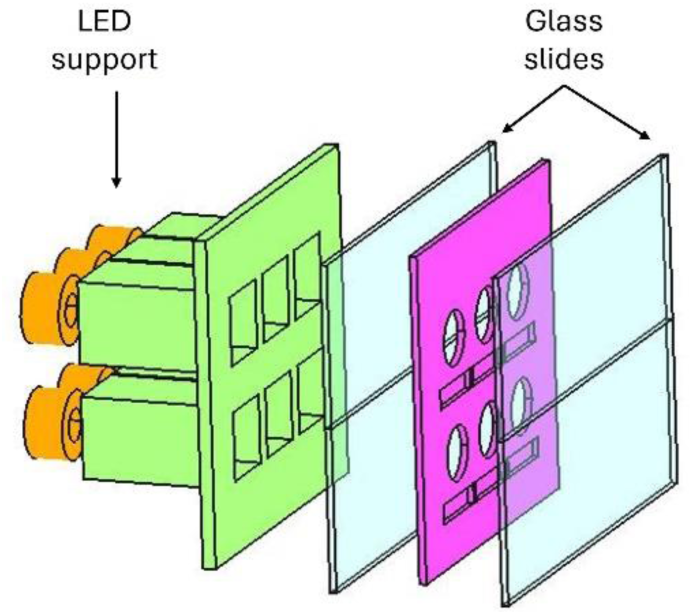
The Mip/SPR pathway modulates the other PMP stages. Schematic drawing of the device used for optogenetical stimulation of wandering larvae.

**Supplementary Video 1**

Example of videos of larvae performing a normal (top, *R48H10>+*) and abnormal GSB (bottom, *R48H10>Mip-IR*) recorded in the pupariation monitor device during the screen and synchronized at the beginning of GSB. Occurrence of pre-GSB contractions is indicated with a yellow hyphen and helps to identify the moment when GSB is expected to occur in those larvae with abnormal or absent GSB. The dashed silhouette marks the starting position of each larva. Larvae usually move forward about one third the length of their body during GSB. Video reproduced at 3X speed.

**Supplementary Video 2**

Wandering larvae fed with 100 µM all-trans retinal (ATR) were stimulated with 10 pulses of green light of 0.5 seconds of duration at 1 Hz. Control larvae not fed with ATR did not respond to light, while those supplemented did a GSB-like behavior.

**Supplementary Video 3**

Footage of Mip2M>ReaChR larvae doing a light induced GSB followed by a spontaneous GSB within the five minute-long unstimulated period, demonstrating that larval response to light constitute a separate behavioral event. Video reproduced at 4x speed.

Time of spontaneous GSB: Left, 01:52; center and right, 01:20

**Supplementary Video 4**

Example of videos of larvae showing muscle GCaMP fluctuations performing a normal (left, wild type) and abnormal GSB (right, *Mip^1^* null mutant). Wild type animals display the whole GSB behavioral sequence. It begins with a tetanic contraction of the anterior segments that expels glue from the salivary glands, followed by back-and-forth peristaltic waves that spread it over the ventral surface. GSB culminates with a "head waving" movement. Larvae that lack Mip neuropeptide can still initiate GSB, but it primarily consists of backwards movements.

## BIBLIOGRAPHY

1. Adams ME, Zitnan D. Identification of ecdysis-triggering hormone in the silkworm Bombyx mori. Biochem Biophys Res Commun. 1997 Jan 3;230(1):188–91. doi: 10.1006/bbrc.1996.5915. PMID: 9020043

2. Baggerman G, Boonen K, Verleyen P, De Loof A, Schoofs L. J. Peptidomic analysis of the larval Drosophila melanogaster central nervous system by two-dimensional capillary liquid chromatography quadrupole time-of-flight mass spectrometry. Mass Spectrom. 2005 Feb;40(2):250–60. doi: 10.1002/jms.744. PMID: 15706625.

3. Belgacem YH, Martin JR. Neuroendocrine control of a sexually dimorphic behavior by a few neurons of the pars intercerebralis in Drosophila. Proc Natl Acad Sci U S A. 2002 Nov 12;99(23):15154–8. doi: 10.1073/pnas.232244199. Epub 2002 Oct 24. PMID: 12399547; PMCID: PMC137559.

4. Berni J, Pulver SR, Griffith LC, Bate M. Autonomous circuitry for substrate exploration in freely moving Drosophila larvae. Curr Biol. 2012 Oct 23;22(20):1861–70. doi: 10.1016/j.cub.2012.07.048. Epub 2012 Aug 30. PMID: 22940472; PMCID: PMC4082562.

5. Blanco-Obregon D, El Marzkioui K, Brutscher F, Kapoor V, Valzania L, Andersen DS, Colombani J, Narasimha S, McCusker D, Léopold P, Boulan L. A Dilp8-dependent time window ensures tissue size adjustment in Drosophila. Nat Commun. 2022 Sep 26;13(1):5629. doi: 10.1038/s41467-022-33387-6. PMID: 36163439; PMCID: PMC9512784.

6. Burke KJ Jr, Bender KJ. Modulation of Ion Channels in the Axon: Mechanisms and Function. Front Cell Neurosci. 2019 May 17;13:221. doi: 10.3389/fncel.2019.00221. PMID: 31156397; PMCID: PMC6533529.

7. Biyasheva A, Do TV, Lu Y, Vaskova M, Andres AJ. Glue secretion in the Drosophila salivary gland: a model for steroid-regulated exocytosis. Dev Biol. 2001 Mar 1;231(1):234–51. doi: 10.1006/dbio.2000.0126. PMID: 11180965.

8. Calleja M, Moreno E, Pelaz S, Morata G. Visualization of gene expression in living adult Drosophila. Science. 1996 Oct 11;274(5285):252-5. doi: 10.1126/science.274.5285.252. PMID: 8824191.

9. Carreira-Rosario A, Zarin AA, Clark MQ, Manning L, Fetter RD, Cardona A, Doe CQ. MDN brain descending neurons coordinately activate backward and inhibit forward locomotion. Elife. 2018 Aug 2;7:e38554. doi: 10.7554/eLife.38554. PMID: 30070205; PMCID: PMC6097840.

10. Carvalho GB, Kapahi P, Anderson DJ, Benzer S. Allocrine modulation of feeding behavior by the Sex Peptide of Drosophila. Curr Biol. 2006 Apr 4;16(7):692–6. doi: 10.1016/j.cub.2006.02.064. PMID: 16581515; PMCID: PMC2745344.

11. Chapman T, Bangham J, Vinti G, Seifried B, Lung O, Wolfner MF, Smith HK, Partridge L. The sex peptide of Drosophila melanogaster: female post-mating responses analyzed by using RNA interference. Proc Natl Acad Sci U S A. 2003 Aug 19;100(17):9923–8. doi: 10.1073/pnas.1631635100. Epub 2003 Jul 31. PMID: 12893873; PMCID: PMC187888.

12. Chen PS, Stumm-Zollinger E, Aigaki T, Balmer J, Bienz M, Böhlen P. A male accessory gland peptide that regulates reproductive behavior of female D. melanogaster. Cell. 1988 Jul 29;54(3):291–8. doi: 10.1016/0092-8674(88)90192-4. PMID: 3135120.

13. Chen J, Kim SM, Kwon JY. A Systematic Analysis of Drosophila Regulatory Peptide Expression in Enteroendocrine Cells. Mol Cells. 2016 Apr 30;39(4):358–66. doi: 10.14348/molcells.2016.0014. Epub 2016 Mar 30. PMID: 27025390; PMCID: PMC4844944.

14. Clark MQ, Zarin AA, Carreira-Rosario A, Doe CQ. Neural circuits driving larval locomotion in Drosophila. Neural Dev. 2018 Apr 19;13(1):6. doi: 10.1186/s13064-018-0103-z. PMID: 29673388; PMCID: PMC5907184.

15. Clark AC, del Campo ML, Ewer J. Neuroendocrine control of larval ecdysis behavior in Drosophila: complex regulation by partially redundant neuropeptides. J Neurosci. 2004 Apr 28;24(17):4283–92. doi: 10.1523/JNEUROSCI.4938-03.2004. PMID: 15115824; PMCID: PMC6729283.

16. Cole SH, Carney GE, McClung CA, Willard SS, Taylor BJ, Hirsh J. Two functional but noncomplementing Drosophila tyrosine decarboxylase genes: distinct roles for neural tyramine and octopamine in female fertility. J Biol Chem. 2005 Apr 15;280(15):14948–55. doi: 10.1074/jbc.M414197200. Epub 2005 Feb 3. PMID: 15691831.

17. Conzelmann M, Williams EA, Tunaru S, Randel N, Shahidi R, Asadulina A, Berger J, Offermanns S, Jékely G. Conserved MIP receptor-ligand pair regulates Platynereis larval settlement. Proc Natl Acad Sci U S A. 2013 May 14;110(20):8224–9. doi: 10.1073/pnas.1220285110. Epub 2013 Apr 8. PMID: 23569279; PMCID: PMC3657792.

18. Cury KM, Axel R. Flexible neural control of transition points within the egg-laying behavioral sequence in Drosophila. Nat Neurosci. 2023 Jun;26(6):1054–1067. doi: 10.1038/s41593-023-01332-5. Epub 2023 May 22. PMID: 37217726; PMCID: PMC10244180.

19. Dao M, Stoveken HM, Cao Y, Martemyanov KA. The role of orphan receptor GPR139 in neuropsychiatric behavior. Neuropsychopharmacology. 2022 Mar;47(4):902–913. doi: 10.1038/s41386-021-00962-2. Epub 2021 Jan 21. PMID: 33479510; PMCID: PMC8882194.

20. Deng B, Li Q, Liu X, Cao Y, Li B, Qian Y, Xu R, Mao R, Zhou E, Zhang W, Huang J, Rao Y. Chemoconnectomics: Mapping Chemical Transmission in Drosophila. Neuron. 2019 Mar 6;101(5):876–893.e4. doi: 10.1016/j.neuron.2019.01.045. Epub 2019 Feb 21. PMID: 30799021.

21. de Velasco B, Erclik T, Shy D, Sclafani J, Lipshitz H, McInnes R, Hartenstein V. Specification and development of the pars intercerebralis and pars lateralis, neuroendocrine command centers in the Drosophila brain. Dev Biol. 2007 Feb 1;302(1):309–23. doi: 10.1016/j.ydbio.2006.09.035. Epub 2006 Sep 26. PMID: 17070515.

22. Dewey EM, McNabb SL, Ewer J, Kuo GR, Takanishi CL, Truman JW, Honegger HW. Identification of the gene encoding bursicon, an insect neuropeptide responsible for cuticle sclerotization and wing spreading. Curr Biol. 2004 Jul 13;14(13):1208–13. doi: 10.1016/j.cub.2004.06.051. PMID: 15242619.

23. Diao F, Ironfield H, Luan H, Diao F, Shropshire WC, Ewer J, Marr E, Potter CJ, Landgraf M, White BH. Plug-and-play genetic access to drosophila cell types using exchangeable exon cassettes. Cell Rep. 2015 Mar 3;10(8):1410–21. doi: 10.1016/j.celrep.2015.01.059. Epub 2015 Feb 26. PMID: 25732830; PMCID: PMC4373654.

24. Dietzl G, Chen D, Schnorrer F, Su KC, Barinova Y, Fellner M, Gasser B, Kinsey K, Oppel S, Scheiblauer S, Couto A, Marra V, Keleman K, Dickson BJ. A genome-wide transgenic RNAi library for conditional gene inactivation in Drosophila. Nature. 2007 Jul 12;448(7150):151-6. doi: 10.1038/nature05954. PMID: 17625558.

25. Denlinger DL. Metamorphosis behavior of flies. Annu Rev Entomol. 1994;39:243–66. doi: 10.1146/annurev.en.39.010194.001331. PMID: 8135500.

26. Erwin PM, Szmant, AM. Settlement induction of Acropora palmata planulae by a GLW-amide neuropeptide. Coral Reefs. 2010; 29, 929–939. doi: 10.1007/s00338-010-0634-1

27. Ewer J. Behavioral actions of neuropeptides in invertebrates: insights from Drosophila. Horm Behav. 2005 Nov;48(4):418–29. doi: 10.1016/j.yhbeh.2005.05.018. Epub 2005 Jul 5. PMID: 15996666

28. Ewer J, Gammie SC, Truman JW. Control of insect ecdysis by a positive-feedback endocrine system: roles of eclosion hormone and ecdysis triggering hormone. J Exp Biol. 1997 Mar;200(Pt 5):869–81. doi: 10.1242/jeb.200.5.869. PMID: 9100362.

29. Fraenkel, G., & Bhaskaran, G. Pupariation and pupation in cyclorrhaphous flies (Diptera): terminology and interpretation. Annals of the Entomological Society of America. 1973; 66(2), 418–422. 10.1093/aesa/66.2.418

30. Fushiki A, Zwart MF, Kohsaka H, Fetter RD, Cardona A, Nose A. A circuit mechanism for the propagation of waves of muscle contraction in Drosophila. Elife. 2016 Feb 15;5:e13253. doi: 10.7554/eLife.13253. PMID: 26880545; PMCID: PMC4829418.

31. Garelli A, Heredia F, Casimiro AP, Macedo A, Nunes C, Garcez M, Dias ARM, Volonte YA, Uhlmann T, Caparros E, Koyama T, Gontijo AM. Dilp8 requires the neuronal relaxin receptor Lgr3 to couple growth to developmental timing. Nat Commun. 2015 Oct 29;6:8732. doi: 10.1038/ncomms9732. PMID: 26510564; PMCID: PMC4640092.

32. Gowda SBM, Salim S, Mohammad F. Anatomy and Neural Pathways Modulating Distinct Locomotor Behaviors in Drosophila Larva. Biology (Basel). 2021 Jan 25;10(2):90. doi: 10.3390/biology10020090. PMID: 33504061; PMCID: PMC7910854.

33. Guo L, Zhang N, Simpson JH. Descending neurons coordinate anterior grooming behavior in Drosophila. Curr Biol. 2022 Feb 28;32(4):823–833.e4. doi: 10.1016/j.cub.2021.12.055. Epub 2022 Feb 3. PMID: 35120659.

34. Hauser F, Neupert S, Williamson M, Predel R, Tanaka Y, Grimmelikhuijzen CJ. Genomics and peptidomics of neuropeptides and protein hormones present in the parasitic wasp Nasonia vitripennis. J Proteome Res. 2010 Oct 1;9(10):5296–310. doi: 10.1021/pr100570j. PMID: 20695486.

35. Heredia F, Volonté Y, Pereirinha J, Fernandez-Acosta M, Casimiro AP, Belém CG, Viegas F, Tanaka K, Menezes J, Arana M, Cardoso GA, Macedo A, Kotowicz M, Prado Spalm FH, Dibo MJ, Monfardini RD, Torres TT, Mendes CS, Garelli A, Gontijo AM. The steroid-hormone ecdysone coordinates parallel pupariation neuromotor and morphogenetic subprograms via epidermis-to-neuron Dilp8-Lgr3 signal induction. Nat Commun. 2021 Jun 7;12(1):3328. doi: 10.1038/s41467-021-23218-5. PMID: 34099654; PMCID: PMC8184853.

36. Hoogstraaten RI, van Keimpema L, Toonen RF, Verhage M. Tetanus insensitive VAMP2 differentially restores synaptic and dense core vesicle fusion in tetanus neurotoxin treated neurons. Sci Rep. 2020 Jul 2;10(1):10913. doi: 10.1038/s41598-020-67988-2. PMID: 32616842; PMCID: PMC7331729.

37. Inagaki HK, Jung Y, Hoopfer ED, Wong AM, Mishra N, Lin JY, Tsien RY, Anderson DJ. Optogenetic control of Drosophila using a red-shifted channelrhodopsin reveals experience-dependent influences on courtship. Nat Methods. 2014 Mar;11(3):325–32. doi: 10.1038/nmeth.2765. Epub 2013 Dec 22. PMID: 24363022; PMCID: PMC4151318.

38. Isabel G, Martin JR, Chidami S, Veenstra JA, Rosay P. AKH-producing neuroendocrine cell ablation decreases trehalose and induces behavioral changes in Drosophila. Am J Physiol Regul Integr Comp Physiol. 2005 Feb;288(2):R531–8. doi: 10.1152/ajpregu.00158.2004. Epub 2004 Sep 16. PMID: 15374818.

39. Isberg V, Andersen KB, Bisig C, Dietz GP, Bräuner-Osborne H, Gloriam DE. Computer-aided discovery of aromatic l-α-amino acids as agonists of the orphan G protein-coupled receptor GPR139. J Chem Inf Model. 2014 Jun 23;54(6):1553–7. doi: 10.1021/ci500197a. Epub 2014 May 30. PMID: 24826842.

40. Jékely G. Global view of the evolution and diversity of metazoan neuropeptide signaling. Proc Natl Acad Sci U S A. 2013 May 21;110(21):8702–7. doi: 10.1073/pnas.1221833110. Epub 2013 May 1. PMID: 23637342; PMCID: PMC3666674.

41. Jenett A, Rubin GM, Ngo TT, Shepherd D, Murphy C, Dionne H, Pfeiffer BD, Cavallaro A, Hall D, Jeter J, Iyer N, Fetter D, Hausenfluck JH, Peng H, Trautman ET, Svirskas RR, Myers EW, Iwinski ZR, Aso Y, DePasquale GM, Enos A, Hulamm P, Lam SC, Li HH, Laverty TR, Long F, Qu L, Murphy SD, Rokicki K, Safford T, Shaw K, Simpson JH, Sowell A, Tae S, Yu Y, Zugates CT. A GAL4-driver line resource for Drosophila neurobiology. Cell Rep. 2012 Oct 25;2(4):991–1001. doi: 10.1016/j.celrep.2012.09.011. Epub 2012 Oct 11. PMID: 23063364; PMCID: PMC3515021.

42. Jovanic T, Schneider-Mizell CM, Shao M, Masson JB, Denisov G, Fetter RD, Mensh BD, Truman JW, Cardona A, Zlatic M. Competitive Disinhibition Mediates Behavioral Choice and Sequences in Drosophila. Cell. 2016 Oct 20;167(3):858–870.e19. doi: 10.1016/j.cell.2016.09.009. Epub 2016 Oct 6. PMID: 27720450.

43. Kim DH, Han MR, Lee G, Lee SS, Kim YJ, Adams ME. Rescheduling Behavioral Subunits of a Fixed Action Pattern by Genetic Manipulation of Peptidergic Signaling. PLoS Genet. 2015 Sep 24;11(9):e1005513. doi: 10.1371/journal.pgen.1005513. PMID: 26401953; PMCID: PMC4581697.

44. Kim YJ, Zitnan D, Galizia CG, Cho KH, Adams ME. A command chemical triggers an innate behavior by sequential activation of multiple peptidergic ensembles. Curr Biol. 2006 Jul 25;16(14):1395–407. doi: 10.1016/j.cub.2006.06.027. PMID: 16860738.

45. Kim YJ, Bartalska K, Audsley N, Yamanaka N, Yapici N, Lee JY, Kim YC, Markovic M, Isaac E, Tanaka Y, Dickson BJ. MIPs are ancestral ligands for the sex peptide receptor. Proc Natl Acad Sci U S A. 2010 Apr 6;107(14):6520–5. doi: 10.1073/pnas.0914764107. Epub 2010 Mar 22. PMID: 20308537; PMCID: PMC2851983.

46. Kipp, M. (2001) Anvil - A Generic Annotation Tool for Multimodal Dialogue. Proceedings of the 7th European Conference on Speech Communication and Technology (Eurospeech), pp. 1367–1370.

47. Kohsaka H, Okusawa S, Itakura Y, Fushiki A, Nose A. Development of larval motor circuits in Drosophila. Dev Growth Differ. 2012 Apr;54(3):408–19. doi: 10.1111/j.1440-169X.2012.01347.x. PMID: 22524610.

48. Kohsaka H, Zwart MF, Fushiki A, Fetter RD, Truman JW, Cardona A, Nose A. Regulation of forward and backward locomotion through intersegmental feedback circuits in Drosophila larvae. Nat Commun. 2019 Jun 14;10(1):2654. doi: 10.1038/s41467-019-10695-y. PMID: 31201326; PMCID: PMC6572865.

49. Kupfermann, Irving, and Klaudiusz R. Weiss. "The command neuron concept." Behavioral and Brain Sciences 1.1 (1978): 3–10.

50. Lam G, Thummel CS. Inducible expression of double-stranded RNA directs specific genetic interference in Drosophila. Curr Biol. 2000 Aug 24;10(16):957–63. doi: 10.1016/s0960-9822(00)00631-x. PMID: 10985382.

51. Lechable M, Jan A, Duchene A, Uveira J, Weissbourd B, Gissat L, Collet S, Gilletta L, Chevalier S, Leclère L, Peron S, Barreau C, Lasbleiz R, Houliston E, Momose T. An improved whole life cycle culture protocol for the hydrozoan genetic model Clytia hemisphaerica. Biol Open. 2020 Nov 5;9(11):bio051268. doi: 10.1242/bio.051268. PMID: 32994186; PMCID: PMC7657476.

52. Leitz T, Morand K, Mann M. Metamorphosin A: a novel peptide controlling development of the lower metazoan Hydractinia echinata (Coelenterata, Hydrozoa). Dev Biol. 1994 Jun;163(2):440–6. doi: 10.1006/dbio.1994.1160. PMID: 7911112.

53. Li HH, Kroll JR, Lennox SM, Ogundeyi O, Jeter J, Depasquale G, Truman JW. A GAL4 driver resource for developmental and behavioral studies on the larval CNS of Drosophila. Cell Rep. 2014 Aug 7;8(3):897–908. doi: 10.1016/j.celrep.2014.06.065. Epub 2014 Jul 31. PMID: 25088417.

54. Lin JY, Knutsen PM, Muller A, Kleinfeld D, Tsien RY. ReaChR: a red-shifted variant of channelrhodopsin enables deep transcranial optogenetic excitation. Nat Neurosci. 2013 Oct;16(10):1499–508. doi: 10.1038/nn.3502. Epub 2013 Sep 1. PMID: 23995068; PMCID: PMC3793847.

55. Liu H, Kubli E. Sex-peptide is the molecular basis of the sperm effect in Drosophila melanogaster. Proc Natl Acad Sci U S A. 2003 Aug 19;100(17):9929–33. doi: 10.1073/pnas.1631700100. Epub 2003 Aug 1. PMID: 12897240; PMCID: PMC187889.

56. Liu C, Bonaventure P, Lee G, Nepomuceno D, Kuei C, Wu J, Li Q, Joseph V, Sutton SW, Eckert W, Yao X, Yieh L, Dvorak C, Carruthers N, Coate H, Yun S, Dugovic C, Harrington A, Lovenberg TW. GPR139, an Orphan Receptor Highly Enriched in the Habenula and Septum, Is Activated by the Essential Amino Acids L-Tryptophan and L-Phenylalanine. Mol Pharmacol. 2015 Nov;88(5):911–25. doi: 10.1124/mol.115.100412. Epub 2015 Sep 8. PMID: 26349500.

57. Lorenz K.Z. The Foundations of Ethology. Springer-Verlag, New York 1981.

58. Lorenz MW, Kellner R, Hoffmann KH. A family of neuropeptides that inhibit juvenile hormone biosynthesis in the cricket, Gryllus bimaculatus. J Biol Chem. 1995 Sep 8;270(36):21103–8. doi: 10.1074/jbc.270.36.21103. PMID: 7673141.

59. Mahr A, Aberle H. The expression pattern of the Drosophila vesicular glutamate transporter: a marker protein for motoneurons and glutamatergic centers in the brain. Gene Expr Patterns. 2006 Mar;6(3):299–309. doi: 10.1016/j.modgep.2005.07.006. Epub 2005 Dec 27. PMID: 16378756.

60. Marder E. Neuromodulation of neuronal circuits: back to the future. Neuron. 2012 Oct 4;76(1):1–11. doi: 10.1016/j.neuron.2012.09.010. PMID: 23040802; PMCID: PMC3482119.

61. Marder E, Kedia S, Morozova EO. New insights from small rhythmic circuits. Curr Opin Neurobiol. 2022 Oct;76:102610. doi: 10.1016/j.conb.2022.102610. Epub 2022 Aug 17. PMID: 35986971.

62. Marder E, Bucher D, Schulz DJ, Taylor AL. Invertebrate central pattern generation moves along. Curr Biol. 2005 Sep 6;15(17):R685–99. doi: 10.1016/j.cub.2005.08.022. PMID: 16139202.

63. McDonald, J.H. 2014. Handbook of Biological Statistics, 3rd ed. Sparky House Publishing, Baltimore, Maryland.

64. Monastirioti M, Gorczyca M, Rapus J, Eckert M, White K, Budnik V. Octopamine immunoreactivity in the fruit fly Drosophila melanogaster. J Comp Neurol. 1995 May 29;356(2):275–87. doi: 10.1002/cne.903560210. PMID: 7629319; PMCID: PMC4664080.

65. Min S, Chae HS, Jang YH, Choi S, Lee S, Jeong YT, Jones WD, Moon SJ, Kim YJ, Chung J. Identification of a Peptidergic Pathway Critical to Satiety Responses in Drosophila. Curr Biol. 2016 Mar 21;26(6):814–20. doi: 10.1016/j.cub.2016.01.029. Epub 2016 Mar 3. PMID: 26948873.

66. Mirabeau O, Joly JS. Molecular evolution of peptidergic signaling systems in bilaterians. Proc Natl Acad Sci U S A. 2013 May 28;110(22):E2028-37. doi: 10.1073/pnas.1219956110. Epub 2013 May 13. PMID: 23671109; PMCID: PMC3670399.

67. Namiki S, Ros IG, Morrow C, Rowell WJ, Card GM, Korff W, Dickinson MH. A population of descending neurons that regulates the flight motor of Drosophila. Curr Biol. 2022 Mar 14;32(5):1189–1196.e6. doi: 10.1016/j.cub.2022.01.008. Epub 2022 Jan 31. PMID: 35090590; PMCID: PMC9206711.

68. Nässel DR, Winther AM. Drosophila neuropeptides in regulation of physiology and behavior. Prog Neurobiol. 2010 Sep;92(1):42–104. doi: 10.1016/j.pneurobio.2010.04.010. Epub 2010 May 4. PMID: 20447440.

69. Nern A, Pfeiffer BD, Rubin GM. Optimized tools for multicolor stochastic labeling reveal diverse stereotyped cell arrangements in the fly visual system. Proc Natl Acad Sci U S A. 2015 Jun 2;112(22):E2967–76. doi: 10.1073/pnas.1506763112. Epub 2015 May 11. PMID: 25964354; PMCID: PMC4460454.

70. Ng M, Roorda RD, Lima SQ, Zemelman BV, Morcillo P, Miesenböck G. Transmission of olfactory information between three populations of neurons in the antennal lobe of the fly. Neuron. 2002 Oct 24;36(3):463–74. doi: 10.1016/s0896-6273(02)00975-3. PMID: 12408848.

71. Ni JQ, Liu LP, Binari R, Hardy R, Shim HS, Cavallaro A, Booker M, Pfeiffer BD, Markstein M, Wang H, Villalta C, Laverty TR, Perkins LA, Perrimon N. A Drosophila resource of transgenic RNAi lines for neurogenetics. Genetics. 2009 Aug;182(4):1089–100. doi: 10.1534/genetics.109.103630. Epub 2009 Jun 1. PMID: 19487563; PMCID: PMC2728850.

72. Ni JQ, Zhou R, Czech B, Liu LP, Holderbaum L, Yang-Zhou D, Shim HS, Tao R, Handler D, Karpowicz P, Binari R, Booker M, Brennecke J, Perkins LA, Hannon GJ, Perrimon N. A genome-scale shRNA resource for transgenic RNAi in Drosophila. Nat Methods. 2011 May;8(5):405–7. doi: 10.1038/nmeth.1592. Epub 2011 Apr 3. PMID: 21460824; PMCID: PMC3489273.

73. Nohr AC, Jespers W, Shehata MA, Floryan L, Isberg V, Andersen KB, Aqvist J, Gutiérrez de Terán H, Brauner-Osborne H, Gloriam DE. The GPR139 reference agonists 1a and 7c, and tryptophan and phenylalanine share a common binding site. Sci Rep. 2017; 7, 1128. 10.1038/s41598-017-01049-z

74. Nusbaum MP, Blitz DM, Marder E. Functional consequences of neuropeptide and small-molecule co-transmission. Nat Rev Neurosci. 2017 Jul;18(7):389–403. doi: 10.1038/nrn.2017.56. Epub 2017 Jun 8. PMID: 28592905; PMCID: PMC5547741.

75. Owald D, Lin S, Waddell S. Light, heat, action: neural control of fruit fly behaviour. Philos Trans R Soc Lond B Biol Sci. 2015 Sep 19;370(1677):20140211. doi: 10.1098/rstb.2014.0211. PMID: 26240426; PMCID: PMC4528823.

76. Park D, Veenstra JA, Park JH, Taghert PH. Mapping peptidergic cells in Drosophila: where DIMM fits in. PLoS One. 2008 Mar 26;3(3):e1896. doi: 10.1371/journal.pone.0001896. PMID: 18365028; PMCID: PMC2266995.

77. Pauls D, von Essen A, Lyutova R, van Giesen L, Rosner R, Wegener C, Sprecher SG. Potency of transgenic effectors for neurogenetic manipulation in Drosophila larvae. Genetics. 2015 Jan;199(1):25–37. doi: 10.1534/genetics.114.172023. Epub 2014 Oct 29. PMID: 25359929; PMCID: PMC4286689.

78. Peabody NC, Pohl JB, Diao F, Vreede AP, Sandstrom DJ, Wang H, Zelensky PK, White BH. Characterization of the decision network for wing expansion in Drosophila using targeted expression of the TRPM8 channel. J Neurosci. 2009 Mar 18;29(11):3343–53. doi: 10.1523/JNEUROSCI.4241-08.2009. PMID: 19295141; PMCID: PMC2717795.

79. Perkins LA, Holderbaum L, Tao R, Hu Y, Sopko R, McCall K, Yang-Zhou D, Flockhart I, Binari R, Shim HS, Miller A, Housden A, Foos M, Randkelv S, Kelley C, Namgyal P, Villalta C, Liu LP, Jiang X, Huan-Huan Q, Wang X, Fujiyama A, Toyoda A, Ayers K, Blum A, Czech B, Neumuller R, Yan D, Cavallaro A, Hibbard K, Hall D, Cooley L, Hannon GJ, Lehmann R, Parks A, Mohr SE, Ueda R, Kondo S, Ni JQ, Perrimon N. The Transgenic RNAi Project at Harvard Medical School: Resources and Validation. Genetics. 2015 Nov;201(3):843–52. doi: 10.1534/genetics.115.180208. Epub 2015 Aug 28. PMID: 26320097; PMCID: PMC4649654.

80. Pfeiffer BD, Jenett A, Hammonds AS, Ngo TT, Misra S, Murphy C, Scully A, Carlson JW, Wan KH, Laverty TR, Mungall C, Svirskas R, Kadonaga JT, Doe CQ, Eisen MB, Celniker SE, Rubin GM. Tools for neuroanatomy and neurogenetics in Drosophila. Proc Natl Acad Sci U S A. 2008 Jul 15;105(28):9715–20. doi: 10.1073/pnas.0803697105. Epub 2008 Jul 9. PMID: 18621688; PMCID: PMC2447866.

81. Pfeiffer BD, Ngo TT, Hibbard KL, Murphy C, Jenett A, Truman JW, Rubin GM. Refinement of tools for targeted gene expression in Drosophila. Genetics. 2010 Oct;186(2):735–55. doi: 10.1534/genetics.110.119917. Epub 2010 Aug 9. PMID: 20697123; PMCID: PMC2942869.

82. Poels J, Van Loy T, Vandersmissen HP, Van Hiel B, Van Soest S, Nachman RJ, Vanden Broeck J. Myoinhibiting peptides are the ancestral ligands of the promiscuous Drosophila sex peptide receptor. Cell Mol Life Sci. 2010 Oct;67(20):3511–22. doi: 10.1007/s00018-010-0393-8. Epub 2010 May 11. PMID: 20458515.

83. Predel R, Rapus J, Eckert M. Myoinhibitory neuropeptides in the American cockroach. Peptides. 2001 Feb;22(2):199–208. doi: 10.1016/s0196-9781(00)00383-1. PMID: 11179813.

84. Predel R, Wegener C, Russell WK, Tichy SE, Russell DH, Nachman RJ. Peptidomics of CNS-associated neurohemal systems of adult Drosophila melanogaster: a mass spectrometric survey of peptides from individual flies. J Comp Neurol. 2004 Jun 28;474(3):379–92. doi: 10.1002/cne.20145. PMID: 15174081.

85. Reiher W, Shirras C, Kahnt J, Baumeister S, Isaac RE, Wegener C. Peptidomics and peptide hormone processing in the Drosophila midgut. J Proteome Res. 2011 Apr 1;10(4):1881–92. doi: 10.1021/pr101116g. Epub 2011 Feb 22. PMID: 21214272.

86. Röder L, Vola C, Kerridge S. The role of the teashirt gene in trunk segmental identity in Drosophila. Development. 1992 Aug;115(4):1017–33. doi: 10.1242/dev.115.4.1017. PMID: 1360402.

87. Sahoo PK, Smith DS, Perrone-Bizzozero N, Twiss JL. Axonal mRNA transport and translation at a glance. J Cell Sci. 2018 Apr 13;131(8):jcs196808. doi: 10.1242/jcs.196808. PMID: 29654160; PMCID: PMC6518334.

88. Santos JG, Vömel M, Struck R, Homberg U, Nässel DR, Wegener C. Neuroarchitecture of peptidergic systems in the larval ventral ganglion of Drosophila melanogaster. PLoS One. 2007 Aug 1;2(8):e695. doi: 10.1371/journal.pone.0000695. PMID: 17668072; PMCID: PMC1933254.

89. Sanyal S. Genomic mapping and expression patterns of C380, OK6 and D42 enhancer trap lines in the larval nervous system of Drosophila. Gene Expr Patterns. 2009 Jun;9(5):371–80. doi: 10.1016/j.gep.2009.01.002. PMID: 19602393.

90. Scheller RH, Jackson JF, McAllister LB, Schwartz JH, Kandel ER, Axel R. A family of genes that codes for ELH, a neuropeptide eliciting a stereotyped pattern of behavior in Aplysia. Cell. 1982 Apr;28(4):707–19. doi: 10.1016/0092-8674(82)90050-2. PMID: 6284369.

91. Schoofs L, Holman GM, Hayes TK, Nachman RJ, De Loof A. Isolation, identification and synthesis of locustamyoinhibiting peptide (LOM-MIP), a novel biologically active neuropeptide from Locusta migratoria. Regul Pept. 1991 Oct 1;36(1):111–9. doi: 10.1016/0167-0115(91)90199-q. PMID: 1796179.

92. Song W, Veenstra JA, Perrimon N. Control of lipid metabolism by tachykinin in Drosophila. Cell Rep. 2014 Oct 9;9(1):40–47. doi: 10.1016/j.celrep.2014.08.060. Epub 2014 Sep 25. Erratum in: Cell Rep. 2020 Feb 18;30(7):2461. PMID: 25263556; PMCID: PMC4325997.

93. Suver MP, Huda A, Iwasaki N, Safarik S, Dickinson MH. An Array of Descending Visual Interneurons Encoding Self-Motion in Drosophila. J Neurosci. 2016 Nov 16;36(46):11768–11780. doi: 10.1523/JNEUROSCI.2277-16.2016. PMID: 27852783; PMCID: PMC5125229.

94. Sweeney ST, Broadie K, Keane J, Niemann H, O’Kane CJ. Targeted expression of tetanus toxin light chain in Drosophila specifically eliminates synaptic transmission and causes behavioral defects. Neuron. 1995 Feb;14(2):341–51. doi: 10.1016/0896-6273(95)90290-2. PMID: 7857643.

95. Takagi S, Cocanougher BT, Niki S, Miyamoto D, Kohsaka H, Kazama H, Fetter RD, Truman JW, Zlatic M, Cardona A, Nose A. Divergent Connectivity of Homologous Command-like Neurons Mediates Segment-Specific Touch Responses in Drosophila. Neuron. 2017 Dec 20;96(6):1373–1387.e6. doi: 10.1016/j.neuron.2017.10.030. Epub 2017 Nov 30. PMID: 29198754.

96. Thum AS, Knapek S, Rister J, Dierichs-Schmitt E, Heisenberg M, Tanimoto H. Differential potencies of effector genes in adult Drosophila. J Comp Neurol. 2006 Sep 10;498(2):194–203. doi: 10.1002/cne.21022. PMID: 16856137.

97. Truman JW. Hormonal control of insect ecdysis: endocrine cascades for coordinating behavior with physiology. Vitam Horm. 2005;73:1–30. doi: 10.1016/S0083-6729(05)73001-6. PMID: 16399406.

98. van den Pol AN. Neuropeptide transmission in brain circuits. Neuron. 2012 Oct 4;76(1):98–115. doi: 10.1016/j.neuron.2012.09.014. PMID: 23040809; PMCID: PMC3918222.

99. Vargas JNS, Sleigh JN, Schiavo G. Coupling axonal mRNA transport and local translation to organelle maintenance and function. Curr Opin Cell Biol. 2022 Feb;74:97–103. doi: 10.1016/j.ceb.2022.01.008. Epub 2022 Feb 24. PMID: 35220080; PMCID: PMC10477965.

100. Veenstra JA. Peptidergic paracrine and endocrine cells in the midgut of the fruit fly maggot. Cell Tissue Res. 2009 May;336(2):309–23. doi: 10.1007/s00441-009-0769-y. Epub 2009 Mar 25. PMID: 19319573.

101. Vömel M, Wegener C. Neuroarchitecture of aminergic systems in the larval ventral ganglion of Drosophila melanogaster. PLoS One. 2008 Mar 26;3(3):e1848. doi: 10.1371/journal.pone.0001848. PMID: 18365004; PMCID: PMC2268740.

102. Warren JT, Yerushalmi Y, Shimell MJ, O’Connor MB, Restifo LL, Gilbert LI. Discrete pulses of molting hormone, 20-hydroxyecdysone, during late larval development of Drosophila melanogaster: correlations with changes in gene activity. Dev Dyn. 2006 Feb;235(2):315–26. doi: 10.1002/dvdy.20626. PMID: 16273522; PMCID: PMC2613944.

103. Williamson M, Lenz C, Winther AM, Nässel DR, Grimmelikhuijzen CJ. Molecular cloning, genomic organization, and expression of a B-type (cricket-type) allatostatin preprohormone from Drosophila melanogaster. Biochem Biophys Res Commun. 2001 Feb 23;281(2):544–50. doi: 10.1006/bbrc.2001.4402. Erratum in: Biochem Biophys Res Commun 2001 Mar 23;282(1):368. Winther ME [corrected to Winther AM]. PMID: 11181081.

104. Yapici N, Kim YJ, Ribeiro C, Dickson BJ. A receptor that mediates the post-mating switch in Drosophila reproductive behavior. Nature. 2008 Jan 3;451(7174):33-7. doi: 10.1038/nature06483. PMID: 18066048.

105. Zarin AA, Mark B, Cardona A, Litwin-Kumar A, Doe CQ. A multilayer circuit architecture for the generation of distinct locomotor behaviors in Drosophila. Elife. 2019 Dec 23;8:e51781. doi: 10.7554/eLife.51781. PMID: 31868582; PMCID: PMC6994239.

106. Zdarek J, Fraenkel G. The mechanism of puparium formation in flies. Journal of Experimental Zoology, 1972, 179(3), 315–323. doi: 10.1002/jez.1401790304

107. Zitnan D, Kingan TG, Hermesman JL, Adams ME. Identification of ecdysis-triggering hormone from an epitracheal endocrine system. Science. 1996 Jan 5;271(5245):88-91. doi: 10.1126/science.271.5245.88. PMID: 8539606

